# Not only a matter of age: Machine learning-based characterization of the differential effect of brain stimulation on skill acquisition

**DOI:** 10.1101/2023.06.14.544579

**Authors:** Pablo Maceira-Elvira, Traian Popa, Anne-Christine Schmid, Andéol Cadic-Melchior, Henning Müller, Roger Schaer, Leonardo G. Cohen, Friedhelm C. Hummel

## Abstract

Brain stimulation shows potential at enhancing cognitive and motor functions in humans. However, multiple studies assessing its effects on behavior show heterogeneous results, especially in healthy older subjects. We propose a new method to predict an individual’s likelihood and the magnitude of the benefit from stimulation, based on the baseline performance of a sequential motor task, framed in the context of their age. Our results show a differential effect of stimulation, in which individuals with less efficient learning mechanisms benefit from stimulation, while those possessing optimal learning strategies resent a detrimental effect. Importantly, this differential effect was determined by one’s ability to integrate task-relevant information at the early stages of training, and not the age. This study paves the way towards the personalized application of stimulation to maximize its effects, and constitutes the first steps to implement an individualized translational clinical intervention, based on the state of the neural system.

**Teaser:** Age notwithstanding, brain stimulation is most effective in deficient neural systems, while being detrimental to optimal systems

**Visual abstract:** 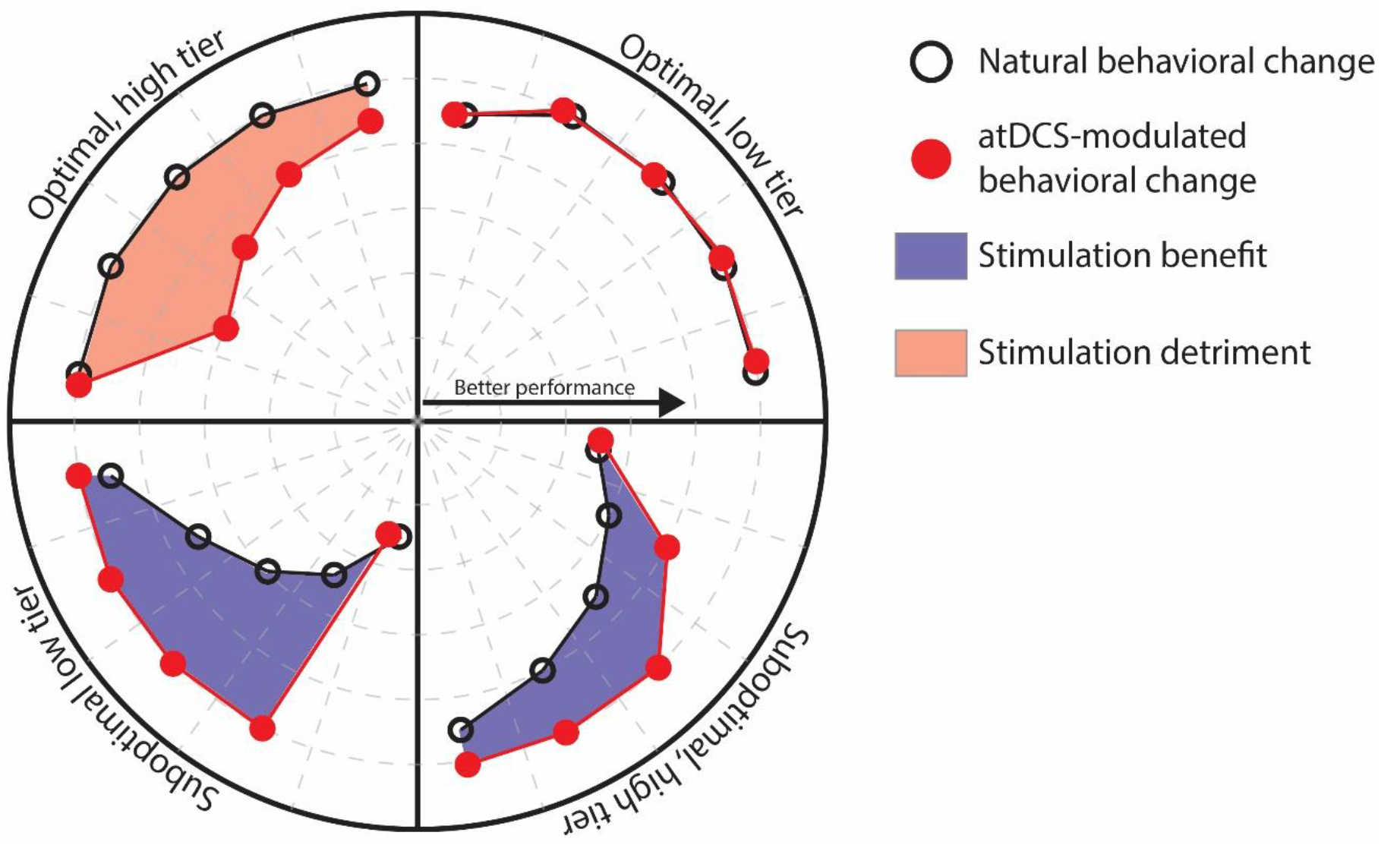

*Main finding:* Anodal transcranial direct current stimulation (atDCS), applied over the hand representation of the motor cortex concomitant to the training of a sequential motor sequence, has differential effects as a function of the recipient’s ability to integrate task-relevant information at the early stages of training. Stimulation benefits individuals with seemingly less efficient learning mechanisms, enabling the rapid storage of the spatial coordinates of the motor sequence and an accelerated optimization of the accuracy of execution. In contrast, individuals possessing optimal learning mechanisms experience detrimental effects of stimulation, leading to drops in the accuracy of execution.

## Introduction

The acquisition of sequential motor skills constitutes a central object of study due to its widespread presence in the completion of activities of daily living. When encountered for the first time, the execution of a sequential motor task entails a speed-accuracy tradeoff (Wickelgren, 1977), in which performing the task at higher speeds often leads to a drop in accuracy. With practice, individuals experience a shift in this tradeoff (Reis et al., 2009; Shmuelof et al., 2012), allowing for an accurate execution of the task at increasing speeds. Optimal motor skill acquisition, characterized by the aforementioned shift in the speed-accuracy tradeoff occurring at the early stages of training, depends on prioritizing the improvement of the accuracy over that of the speed (Maceira-Elvira et al., 2022). Having optimized the accuracy by storing the spatial coordinates of the sequence in memory (Ghilardi et al., 2009), individuals are able to focus on the improvement of their speed (Hikosaka et al., 1999). In the absence of external temporal cues, individuals tend to group easier effector transitions (e.g., adjacent finger movements in a finger-tapping task) and to execute them in close temporal proximity (Popp et al., 2020) giving rise to temporal patterns commonly referred to as motor chunks (Rosenbaum et al., 2007; Sakai et al., 2003; Verwey, 1996)). The prime placed on the speed of execution influences the rate at which chunking patterns are generated (Maceira-Elvira et al., 2022), with a higher prime placed on speed leading to such patterns emerging sooner.

The acquisition of sequential motor tasks is often less efficient in older adults, who frequently depict an overall lower performance in the execution of such tasks (Maceira-Elvira et al., 2022; Shea et al., 2006; Verwey, 2010; Zimerman et al., 2013). As such, the plasticity-augmenting properties of non-invasive brain stimulation (NIBS) (Bolognini et al., 2009), constitute a promising option to improve motor skill acquisition in older adults (Zimerman & Hummel, 2010). Notably, anodal transcranial direct current stimulation (atDCS) applied over the motor cortex can improve motor skill acquisition in older adults (Fujiyama et al., 2017; Summers et al., 2016; Zimerman et al., 2013) by accelerating the shift in the speed-accuracy tradeoff, streamlining the acquisition of sequential motor tasks (Maceira-Elvira et al., 2022). This effect of atDCS appears to be reserved to the early stages of training (i.e., mostly to the first training session), and exclusive to individuals with age-related diminished learning abilities (Maceira-Elvira et al., 2022). However, some previous studies have shown beneficial effects of atDCS in young adults as well (Hashemirad et al., 2016; Karok & Witney, 2013; Saucedo-Marquez et al., 2013; Wessel et al., 2021). This raises the question of what are the determinant factors behind an individual’s response to a NIBS intervention, such as atDCS, possibly accounting for the large variability in its effects on motor skill acquisition (Hashemirad et al., 2016; Summers et al., 2016).

Here, we studied the effects of atDCS applied concomitant to motor training in middle-aged and older adults over the course of ten days. After finding the same effect of atDCS on the early optimization of the accuracy as reported before (Maceira-Elvira et al., 2022), we estimated each individual’s ability to acquire a sequential motor task, as well as their likelihood to benefit from atDCS, based on their initial performance in this task. Our results suggest a differential effect of atDCS, granting larger benefits to individuals with less efficient learning mechanisms, while being detrimental to individuals possessing optimal learning strategies. These effects were neither necessarily characteristic of, nor reserved to, individuals of a certain age.

## Results

### The effect of atDCS applied during extensive motor training

We studied the effects of applying atDCS over the left-hand representation of the primary motor cortex (M1) during the acquisition of a well-established finger-tapping task (Draaisma et al., 2022; Maceira-Elvira et al., 2022; Walker et al., 2003; Wessel et al., 2021; Zimerman et al., 2013) in middle-aged (50-65 y/o; n = 20, 11 female; age*μ* = 59.05 y/o) and older adults (>65 y/o; n = 20, 10 female; age*μ* = 71.7 y/o), as they practiced over the course of ten days. The participants received either verum stimulation (*i*.*e*., active; middle-aged = 10, age*μ* = 58.9; older = 10, age*μ* = 71.4) or placebo stimulation (middle-aged = 10, age*μ* = 59.2; older = 10, age*μ* = 72.1) for 20 minutes daily, as they practiced the finger-tapping task using their left hand. The task consisted in replicating an explicitly shown, nine-digit numerical sequence as fast and as accurately as possible using four buttons. To simplify the report and the discussion of results for each group, we will refer to these groups as “middle-verum”, “middle-placebo”, “older-verum” and “older-placebo” henceforth. Each training day consisted of seven 90-second practice blocks interleaved with 90 second resting periods; six of the practice blocks contained a fixed sequence (i.e., training sequence), while the seventh contained a different sequence used to assess the generalization of the participants’ performance (i.e., “catch blocks”). We applied the same analytical pipeline developed for our previous study (Maceira-Elvira et al., 2022), describing the performance of our participants in terms of the main drivers for this task (i.e., the speed and the accuracy), and the influence of these parameters in the mechanical execution of the sequence. Our previous quantification of general performance in this task consisted of a combination of speed and accuracy, which was useful to describe aspects such as online (within training sessions) and offline (between training sessions) behavioral changes (Dayan & Cohen, 2011)). Nevertheless, as our previous findings (Maceira-Elvira et al., 2022) revealed an effect of atDCS exclusive to the accuracy (and not the speed), and no effect on offline learning, we chose to focus our present discussion on the evolution of both the speed and the accuracy separately. Please refer to the *Supplementary Materials* to find the scores obtained by the participants (both in the training and the “catch blocks”), as well as the contribution of online and offline learning to the overall improvement seen towards the end of training.

*Figure 1* shows the speed and the accuracy, as well as the dynamics of these two parameters, for the groups of middle-aged and older adults. Please note that these data depict the participants’ performance in the trained sequence only (i.e., and not the catch blocks, as there was a consistent drop in performance for these blocks, please see the *Supplementary Materials*). Speed (*Figure 1A*), quantified as the number of sequences generated by the participants on each training block, was generally higher in the middle-placebo group compared to the older-placebo. However, the statistical testing of this difference showed only a trend towards significance (F_[1,18]_ = 3.12, η^2^ = 0.15, p = 0.09), and we did not find evidence of their initial speed being different either (T_[18]_ = 1.16, d = 0.52, p = 0.25). We did not find evidence for an effect of atDCS on the speed of execution in middle-aged adults, neither in absolute terms (F_[1,18]_ = 0.43, η^2^ = 0.02, p = 0.51) nor in the rate of improvement (F_[1,1118]_ = 1.19, η^2^ = 0.001, p = 0.27). In older-adults, the older-verum group was initially slower compared to the older-placebo (although the evidence had only a trend towards significance (T_[12]_ = 1.84, d = 0.82, p = 0.09)), and the difference in the rate of speed change (i.e., slope) was not statistically significant (F_[1,1108]_ = 0.39, η^2^ = 0.0003, p = 0.53). As for the overall speed dynamics (*Figure 1C*), the evolution of the speed over the course of training was similar among the groups. As such, these results do not provide evidence for an effect of stimulation on the speed of execution in neither middle-aged nor older adults, matching our previous findings (Maceira-Elvira et al., 2022).

**Fig. 1.**
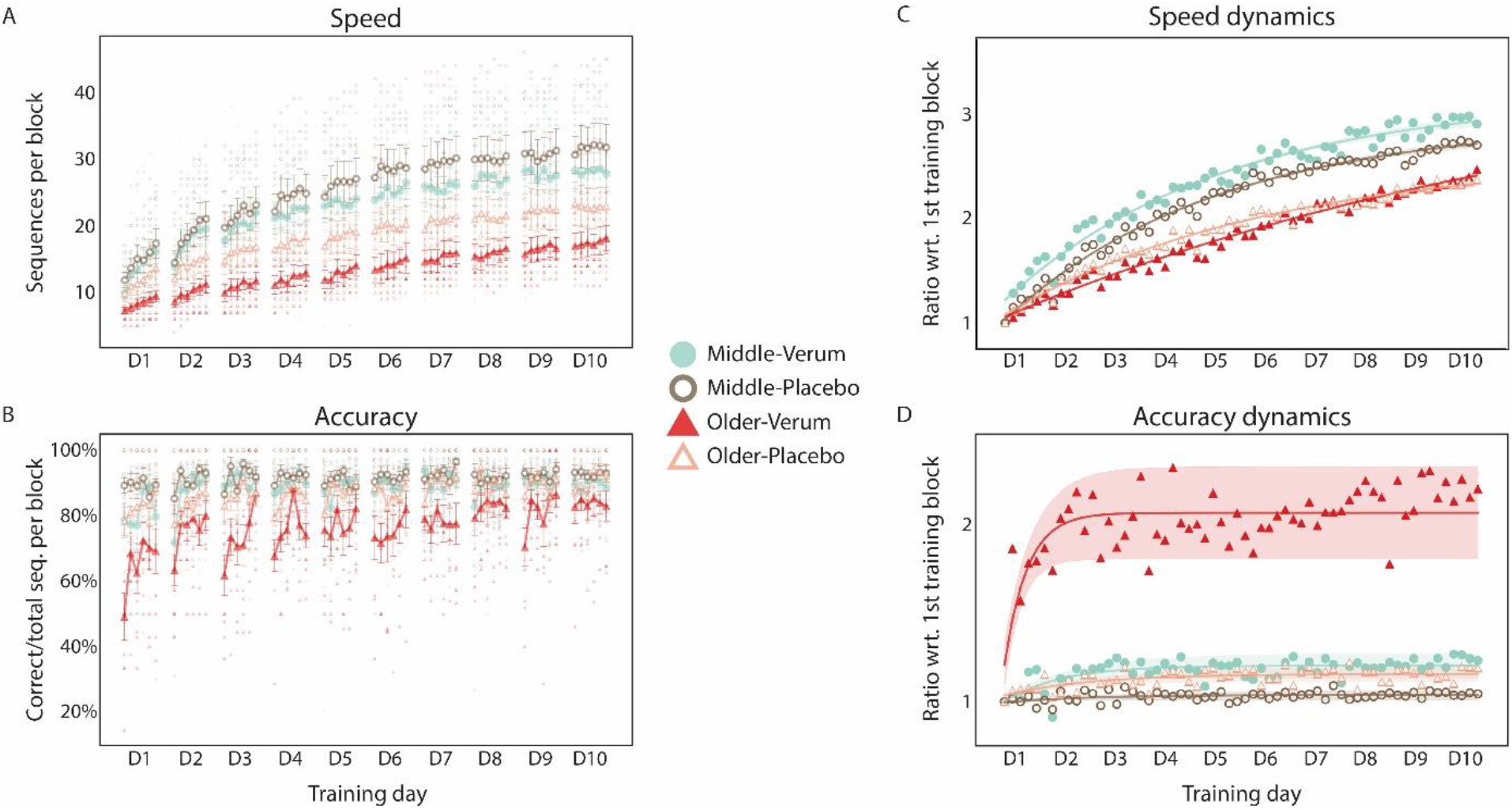
Speed and accuracy over training of the sequence-tapping task. **(A)** Speed in middle-aged and older adults under verum and placebo stimulation, quantified as the total number of generated sequences in each block. The smaller markers portray individual data, while the larger markers reflect the average for each group, and the error bars represent the standard error of the mean. **(B)** Accuracy in the training blocks, calculated as the ratio of correct to total sequences, with the smaller markers portraying individual data and the larger markers reflecting the average for each group. The error bars represent the standard error of the mean. **(C)** Speed dynamics, calculated as the ratio of the average speed of each training block, divided by the average speed of the first block of training within each group. Please note that in spite of the differences in the magnitude of the speed among the age groups, the general improvement follows similar trajectories across all groups. **(D)** Accuracy dynamics, calculated as the ratio of the average accuracy of each training block, divided by the average accuracy of the first block of training within each group. Please note the stark difference between the older-verum, characterized by having initially low accuracies, and the other three groups, starting off at much higher accuracy values. The dynamics in the older-verum group, depicting a sharp increase and stabilization occurring at the early stages of training, match our previous findings in older adults (Maceira-Elvira et al., 2022). The shaded area represents the 95% confidence interval for the fitted curve.

*Figure 1B* shows the accuracy, calculated as the ratio of correct sequences to total sequences generated by each participant. The middle-placebo group was significantly more accurate than the older-placebo throughout training (F_[1,18]_ = 10.06, η^2^ = 0.36, p = 0.005). In middle-aged adults, we did not find the accuracy to be different when comparing both stimulation groups (F_[1,18]_ = 2.97, η^2^ = 0.14, p = 0.10). In older adults, the accuracy was significantly higher in the placebo group compared to verum throughout training (F_[1,18]_ = 21.08, η^2^ = 0.54, p = 0.0002), starting from the first training block (T_[15]_ = 3.44, d = 1.54, p = 0.003, *Figure 1B*). The slopes were significantly different (F_[1,1108]_ = 9.30, η^2^ = 0.008, p = 0.002) between the two stimulation groups, with a steeper improvement of the verum group (T_[1108]_ = 3.05, d = 13.61, p = 0.002). These differences are illustrated in *Figure 1D*. The older-verum group, with lower initial accuracy, improved sharply early in training, reaching a plateau and maintaining similar levels for the remainder of training.

The results obtained from both groups of middle-aged adults and the older-verum group were in correspondence with our previous findings (Maceira-Elvira et al., 2022). However, the initial values for speed and accuracy seen in the older-placebo group were higher, more similar to those seen in middle-aged adults. In view of this unexpected difference in baseline performance, we conducted additional comparisons to verify the apparent effect of the stimulation on the accuracy of older adults was not an artifact derived from their initially low skill level, and to better understand the underlying surrogate.

### The effect of atDCS in comparably inaccurate individuals

The dynamics we observe in the accuracy of the older-verum group are consistent with our previous findings (Maceira-Elvira et al., 2022). However, in that study both the verum and the placebo groups of older adults had similar accuracies initially, whereas in the present study the accuracy in the placebo group was much higher in the beginning. Therefore, a possible explanation for the sharp increase in accuracy in the verum group could be that they started from much lower values and had, thus, more space for improvement. To test whether this was true, we selected individuals from the older-placebo group with initial accuracies within the range of accuracies seen in the older-verum group. The range of accuracies in the first training block of this group went from 14% to 62.5%, excluding one individual with an initial accuracy of 100%, so we considered a range of 0 to 62.5 for this comparison. As there were very few participants with initial accuracies within this range in the older-placebo group in the present study (n = 2), we included individuals from the older-placebo group (n = 4), as well as the unstimulated cohort (n = 5) from our previous study (Maceira-Elvira et al., 2022). In total, we compared nine individuals from the older-verum group of this study to eleven older adults under either placebo or no stimulation (older-placebo-unstimulated group). *Figure 2* shows the accuracy dynamics for these two groups, contrasting the sharp increase in accuracy in the older-verum to a gradual increase in the older-placebo-unstimulated group, even though individuals in both groups possessed similar accuracies on the first block of training. For this comparison, we included only the first five days of training of the present study, as in our previous study, participants trained for only five days (Maceira-Elvira et al., 2022).

**Fig. 2.**
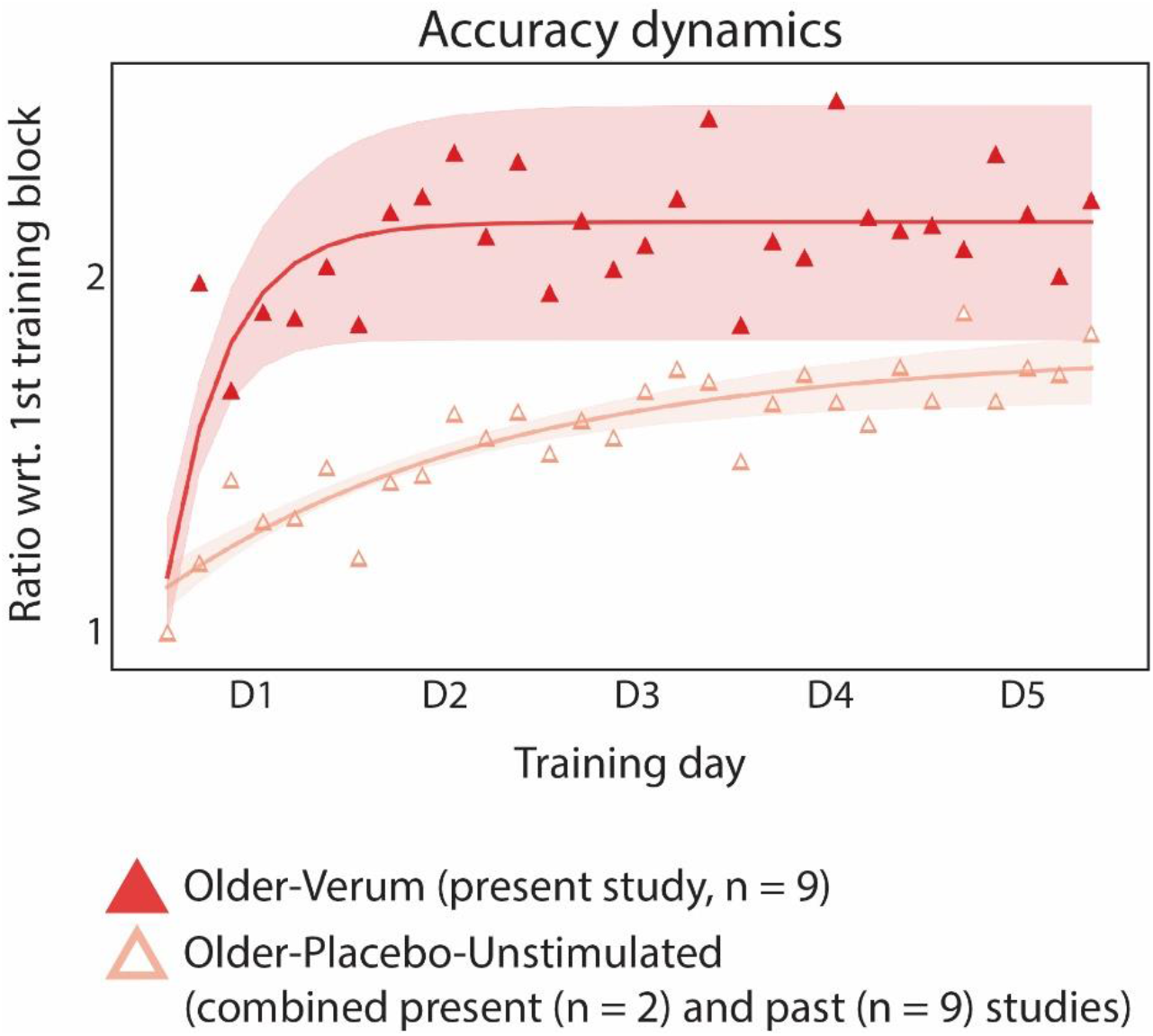
Accuracy dynamics in older adults with initial accuracies equal or lower than 62.5%. We compared individuals from the older-verum group of this study (red, full triangles, n = 9) to individuals from the older-placebo groups of this study (n = 2) and an earlier study (n = 4), as well as unstimulated older individuals from that same study (n = 5) (Maceira-Elvira et al., 2022) (older-placebo-unstimulated, pink, hollow triangles). Of note is that even though all older adults start off at similar accuracies, the older-verum group experiences a sharp accuracy increase on the first training day and reaches a plateau, while the older-placebo-unstimulated group increases their accuracy gradually over the course of training. The shaded area represents the 95% confidence interval for the fitted curve. Please note there are only five days of training displayed, as opposed to the ten days of training shown in Figure 1. The reason is that in the past studies we conducted, participants were required to practice for five days instead of ten.

### Streamlining the mechanical execution of the motor sequence

In a previous study (Maceira-Elvira et al., 2022), we found that optimizing the accuracy at the early stages of training enabled the streamlining of the execution of the sequence, leading to the generation of efficient temporal patterns commonly referred to as motor chunks (Rosenbaum et al., 2007; Sakai et al., 2003; Verwey, 1996). In turn, the generation of efficient chunking patterns seemed to depend on the prime placed on speed, with a higher prime leading to an earlier formation of chunking patterns, and a lower prime resulting in such patterns appearing later in training. Based on those observations, we expected individuals from the older-placebo group of the present study to generate efficient chunking patterns sooner than those in the older-verum group, as they were significantly more accurate and faster from the beginning. *Figure 3* shows the efficiency of generated chunking patterns by all individuals of this study. We quantified the efficiency in the same way and using the same model as in our previous study (Maceira-Elvira et al., 2022).

**Fig. 3.**
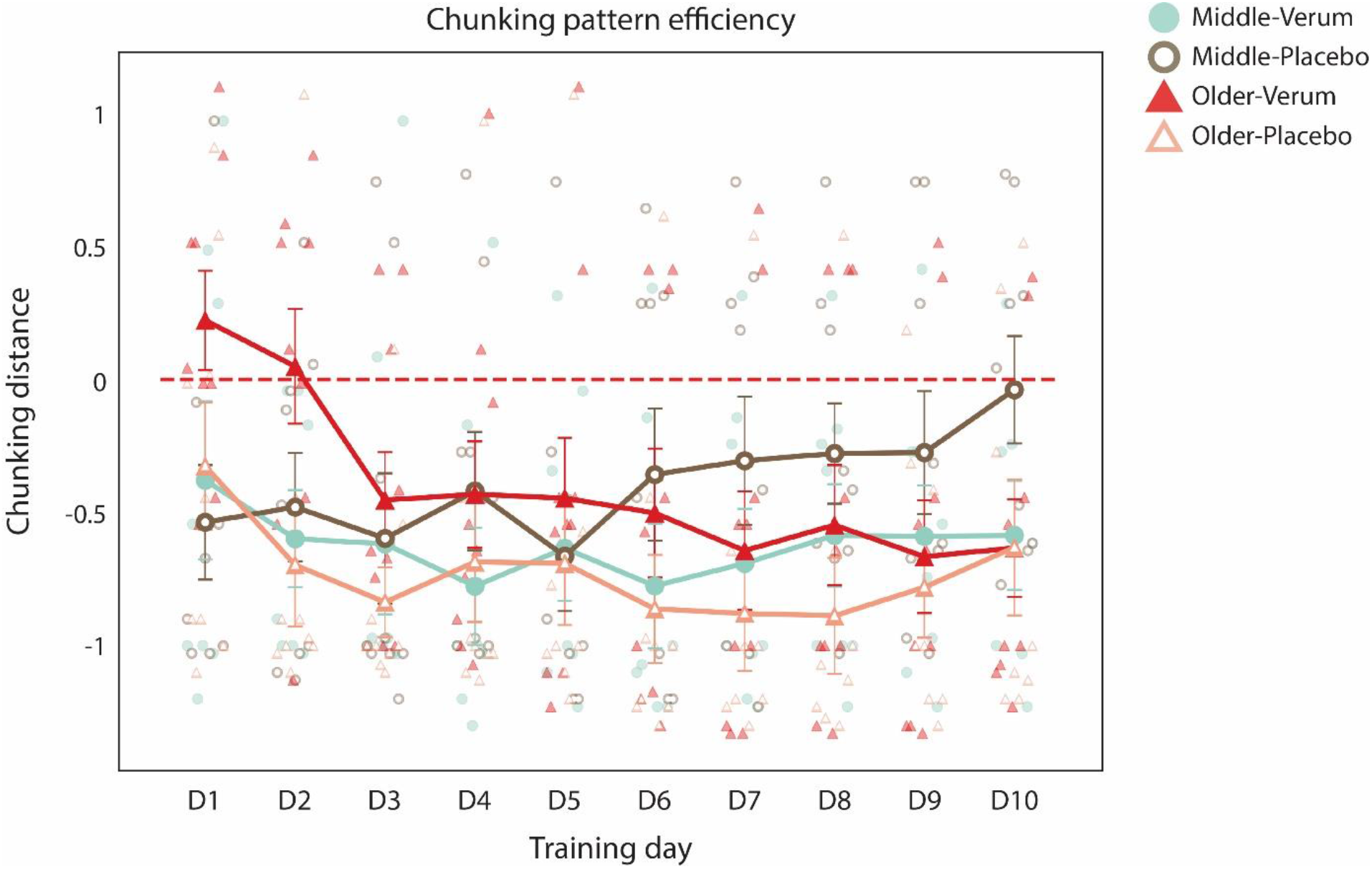
Chunking pattern efficiency. Evolution of the chunking patterns generated by all participants over the course of training. The measure of efficiency is the “chunking distance” described in a previous paper (Maceira-Elvira et al., 2022), which values represent more efficient (i.e., negative values) or less efficient (i.e., positive values) chunking patterns emerging during the execution of the finger-tapping task. The smaller markers depict the values assigned to each individual on each day of training, while the larger markets illustrate the average values per day. The error bars represent the standard error of the mean. Please note that most individuals in both groups of middle-aged adults and the older-placebo group, possessing relatively high speeds and accuracies at the beginning of training (see Figure 1A and 1B), generate efficient chunking patterns on the first training day, which do not seem to vary much over the remainder of training. In contrast the older-verum group, with initially lower speed and accuracy compared to the other three groups, generate efficient chunking patterns at a later stage during training. However, matching our previous results (Maceira-Elvira et al., 2022), half of the participants in this group generate efficient chunking patterns on the first day, with most of the others doing so by the third day.

Our results show that almost all individuals from the middle-verum, middle-placebo and older-placebo groups generated efficient chunking patterns from the first day of training. This meets our expectations, as the three groups had initially high accuracy and speed. In the older-verum group, about half of the participants generated efficient patterns on the first training day, with most of them doing so by the third day of training; this is the same behavior we found in our previous study for the older-verum (Maceira-Elvira et al., 2022).

### The likelihood to benefit from stimulation and its differential effect

In a previous study (Maceira-Elvira et al., 2022), we found atDCS not to have an effect in neither young nor middle-aged adults, with its beneficial effects seemingly reserved to older adults. Our interpretation of these results was that atDCS had a restorative rather than an enhancing effect (at least for the motor task in question), which could explain why it only appeared to benefit individuals with seemingly diminished learning mechanisms. However, the results of the present study showed that being above a certain age does not necessarily result in a less efficient neural system; indeed, the average performance of the older-placebo group in the present study was similar to the one seen in middle-aged adults, both initially and throughout training. Therefore, applying atDCS to its full potential may require knowledge on the status of the neural mechanisms supporting skill acquisition, for which age alone does not provide enough information. With this in mind, we implemented an algorithm to determine whether an individual would be more or less likely to benefit from atDCS in the acquisition of the finger-tapping task.

Our working hypothesis is that the optimal acquisition of the finger-tapping task relies on prioritizing the improvement of the accuracy at the early stages of training. Under this hypothesis, we would consider individuals reaching a plateau in their accuracy on the first training session to be “optimal learners”, while those reaching a plateau at a later stage would be considered “suboptimal”. Further, in the context of our interpretation of the differential effects of stimulation, we would expect optimal learners not to benefit from atDCS, while suboptimal learners would profit from stimulation, with larger boons expected for worse learners. In regard to the prioritization of the accuracy over the speed, we cannot entirely rule out the possibility of it being a matter of personal preference. However, the marked accuracy dynamics related to the application of atDCS suggest this process may be driven by the system’s ability to manage neural resources, with the goal of maximizing performance while minimizing cognitive load.

Following this reasoning, we implemented an algorithm to determine the likelihood of an individual to reach a plateau in accuracy on the first day of training, based on baseline parameters. Specifically, we trained a classifier to label individuals as either “optimal” or “suboptimal” learners based on their age, and measures of their initial speed and initial accuracy (please refer to *Methods* for details on the features drawn). We trained the classifier with both unstimulated or placebo-stimulated participants from our previously published dataset (Maceira-Elvira et al., 2022), using 80% of the data for training and 20% for testing, achieving a prediction accuracy of 97% (i.e., F1 score). Please note that we intentionally used all the aforementioned data for training the model (i.e., without leaving a validation set aside), so this score only indicates the separability of the training features. After training the classifier, we projected the predictions from the model into a sigmoid function (i.e., logistic function) to estimate the likelihood of each individual to reach a plateau on the first training day. *Figure 4* illustrates the logistic function, with values above 50% classified as “Suboptimal” (i.e., with above-chance likelihood to benefit from stimulation).

**Fig. 4.**
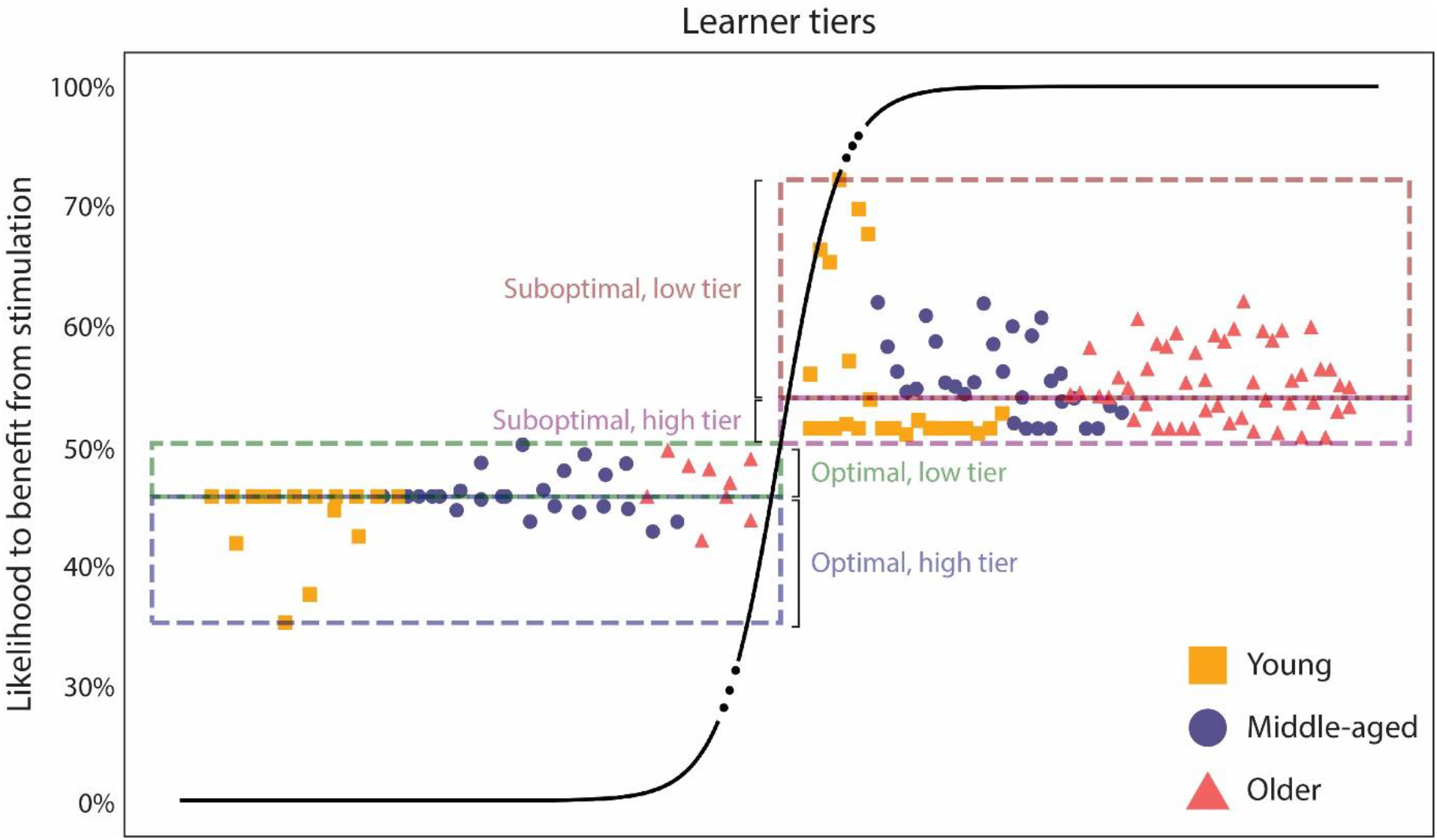
The likelihood of an individual’s response to stimulation. We trained a classifier to discriminate between optimal and suboptimal learners, using data from a previously published dataset (Maceira-Elvira et al., 2022). Then, we extracted the distance between each individual and the hyperplane separating both classes of learners, and projected it onto a sigmoid function to generate a map of probabilities, with those above 50% belonging to the “suboptimal” class, thus having a chance to benefit from stimulation. Each class is further divided into two segments (i.e., tiers), based on the median split of the training data. Of note is that each tier contains individuals from the three age groups (i.e., young, middle-aged and older), challenging the general notion of older adults being worse learners than young adults. Please note that there is no meaning to the x-axis, so the age clusters forming along the horizontal axis bear no meaning.

This model provides a continuous range of the likelihood of a person to benefit from stimulation. We proceeded to segment this range into four learner tiers, namely higher-optimal tier, lower-optimal tier, higher-suboptimal tier and lower-suboptimal tier, expecting the effect of stimulation to be more pronounced as a function of the distance from the higher-tier of optimal learners. In other words, we expected the effect of stimulation to be apparent in both tiers of suboptimal learners, with larger effects present in the lower-tier. The definition of the boundary between adjacent tiers in both optimal and suboptimal learners was based on the median split of the probabilities estimated from the training set (i.e., from previously published data (Maceira-Elvira et al., 2022)). Please refer to *Methods* for more details on this model.

Once we had built the classifier and defined the different categories of learners, we applied the model to the data from the present study to estimate the likelihood of each participant to benefit from stimulation. *Figure 5* shows the accuracy dynamics for the four categories of learners. Please note that each category contains data from participants belonging to all age groups of our previous (Maceira-Elvira et al., 2022) and present studies (please refer to *Table 1* to find the demographics describing the participants of each category).

**Fig. 5.**
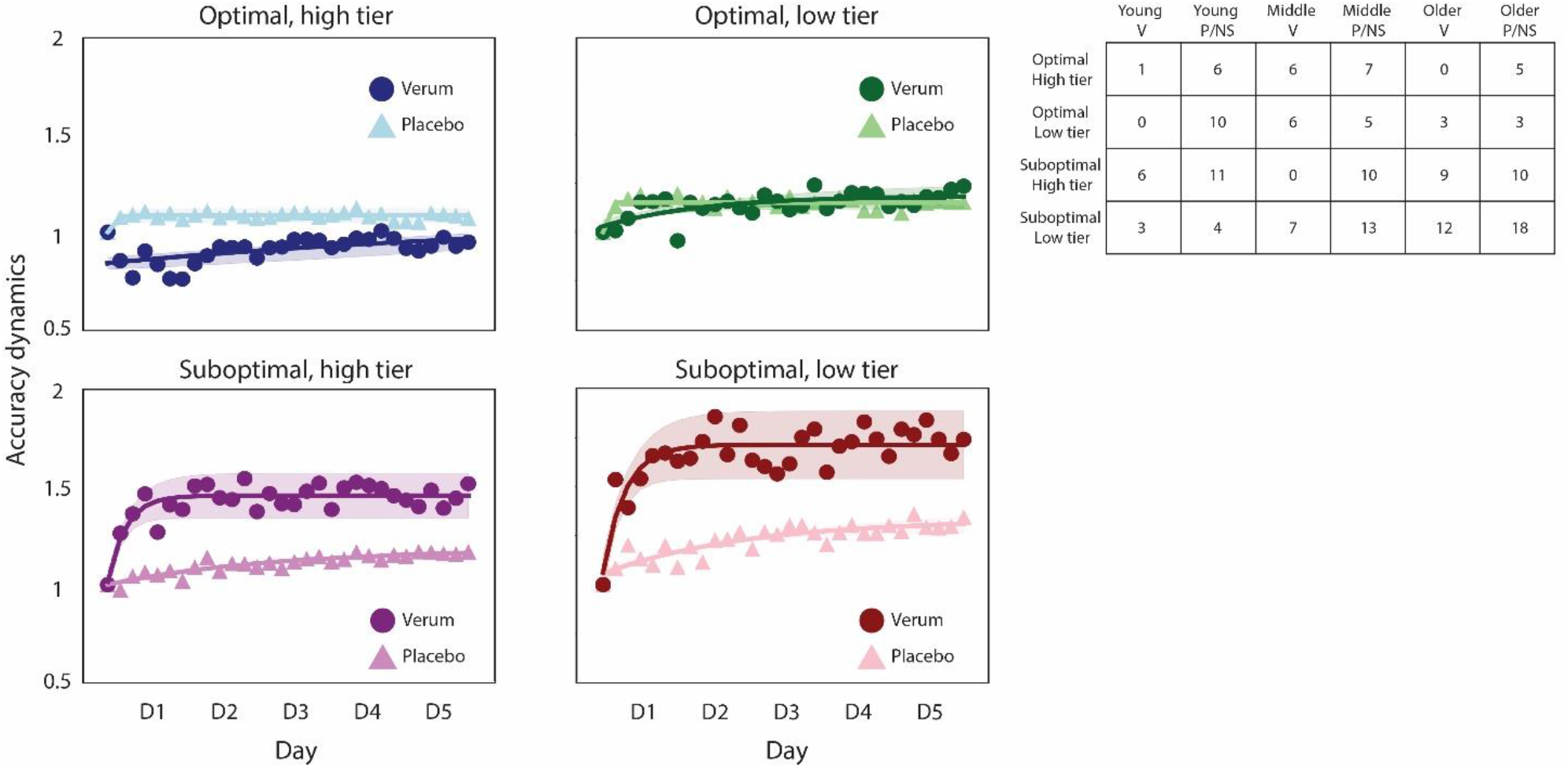
Accuracy dynamics for the different learner tiers. Accuracy dynamics, calculated as the ratio of the average accuracy of each training block, divided by the average accuracy of the first block of training within each group. The data depicted combines the dataset of the present study with our previously-published datasets (Maceira-Elvira et al., 2022), so each group contains individuals from all age groups (see Figure 4 and the table in this figure). In the table, each column corresponds to young, middle-aged and older adults receiving verum (V), placebo (P) or no stimulation (NS). Please note the sharp increase in accuracy in the verum groups of suboptimal learners, reminiscent of the dynamics we had seen before in the older-verum group, with more pronounced increases in the low tier of suboptimal learners, suggesting stronger effects in less skilled learners. Of note is also the evolution of the accuracy in the group of most skilled learners (i.e., optimal, high tier) receiving verum stimulation, as they seem to worsen at the early stages of training, suggesting a detrimental effect of stimulation in individuals with optimally-tuned neural systems. The shaded area represents the 95% confidence interval for the fitted curve.

Our results confirmed the differential effect of stimulation we had hypothesized, with more pronounced effects in the fourth category (i.e., the lower-tier of suboptimal learners) than in higher-tier suboptimal learners. The sharp improvement in accuracy we have proposed to be derived from the application of atDCS was present in both groups of suboptimal learners, supporting our previous findings on the effects of atDCS in the acquisition of the finger-tapping task (Maceira-Elvira et al., 2022). Please note that while these two categories have a majority of individuals over the age of 65 (i.e., older adults), there are both young and middle-aged adults in both categories of suboptimal learners.

In regards to both categories of optimal learners, our expectation of having little or no effect of stimulation was confirmed by the lower-tier of optimal learners, for which the dynamics appear to be quite similar. In contrast, individuals in the higher-tier of optimal learners seemed to even worsen at the early stages of training, suggesting a detrimental effect of stimulation for individuals with optimally-functioning neural systems.

### Electrophysiology

We used the SICI paradigm (Heise et al., 2013; Kujirai et al., 1993; Wessel et al., 2019) of transcranial magnetic stimulation (TMS) to quantify GABAergic intracortical inhibition within M1, and to assess any changes related to the application of stimulation after the ten training days. We did not find statistically significant changes in the SICI recordings performed at rest in any of the groups, providing no evidence for neither training-derived nor stimulation-derived effects. The modulation of SICI, assessed by applying this paradigm during movement preparation (please refer to the *Methods* for more details), changed significantly in the middle-verum group (T_[27]_ = 3.01, d = 1.56, p = 0.005), and was significantly higher than in the middle-placebo group (T_[27]_ = 3.65, d = 1.95, p = 0.001). In older adults, there was only a trend towards a significant difference between verum and placebo (F_[1]_ = 3.61, η^2^ = 0.1, p = 0.06), and no evidence for a significant change after training (F_[1]_ = 0.46, η^2^ = 0.01, p = 0.5).

## Discussion

The present work adds to the understanding of the differential, individual effect of atDCS on the acquisition of a previously unknown sequential motor task, which was more pronounced in individuals with less efficient learning mechanisms, while being detrimental to those expected to learn the task most efficiently.

Previous research shows atDCS can improve motor skill acquisition (Hashemirad et al., 2016; Summers et al., 2016), but the direction and magnitude of this effect varies among reports and is difficult to predict for each individual. Therefore, we implemented a machine learning method to label individuals as either optimal or suboptimal learners, and studied the effect of stimulation in each of the resulting categories. Our results showed the least capable learners among our cohorts to benefit from stimulation to the largest extent, with this benefit being less pronounced as initial skill levels increased. At the other extreme of the spectrum, the most skilled learners showed even detrimental effects of atDCS, suggesting an interfering effect of stimulation on optimal neural systems.

### Support for the model of efficient motor skill acquisition

In a previous study (Maceira-Elvira et al., 2022), we found evidence in favor of a model initially proposed by Hikosaka and colleagues (Hikosaka et al., 1999), in which the acquisition of sequential tasks consists of two stages: the storage of spatial coordinates (i.e., the ordered elements of the sequence) early during training, followed by the storage of motor coordinates involved in processing the timing of execution (i.e., temporal coordinates) later on. We found the efficient execution of the sequence (i.e., the generation of efficient motor chunks), assumed to minimize the computational costs in the neural system while maximizing motor efficacy (Ramkumar et al., 2016), depends on the previous storage of spatial coordinates in memory. However, the emergence of efficient chunking patterns did not seem to be a direct consequence of storing the spatial coordinates, but rather the result of an optimization process on the mechanical execution of the sequence, driven by the prime placed on speed (Maceira-Elvira et al., 2022). In other words, we found that once the spatial coordinates of the sequence were stored in memory, individuals replicating the motor sequence at a higher speed produced efficient chunking patterns sooner and required fewer repetitions of the sequence (i.e., fewer practice) to do so.

In the context of this model, the expected pattern for an individual capable of learning the finger-tapping task optimally would be the following: prioritization of the accuracy at the early stages of training with speed improvements thereafter, with an efficient mechanical execution of the sequence achieved sooner in faster individuals. In the present study, both groups of middle-aged adults and the group of older adults receiving placebo stimulation followed this pattern. Their accuracy reached a plateau on the first training day, their speed of execution was relatively high (i.e., compared to the one seen in the group of older adults receiving verum stimulation), and the chunking patterns they generated on the first training day were recognized as “efficient” by our classifier. As such, these observations further support our proposed model for the optimal acquisition of sequential motor tasks, which can serve as a reference for the detection of diminished learning mechanisms and the identification of likely targets for interventions.

### The neural correlates of the effect of atDCS on motor skill acquisition and its differential effect

atDCS is assumed to facilitate the firing of neuronal populations within the stimulated area by altering the neurons’ resting membrane potential, effectively lowering the neuronal threshold and thus increasing the likelihood of neurons depolarizing (Otmakhov et al., 1993), resulting in an increased cortical excitability (Nitsche & Paulus, 2000). Further, pharmacological studies (Ziemann et al., 1996) have revealed a decrease in GABAergic intracortical inhibition resulting from the application of atDCS (Nitsche et al., 2005), thought to induce long-term potentiation-(LTP-) like plasticity (Fritsch et al., 2010). However, we did not find such a change, as measured by the SICI paradigm; this matched previous reports, in which no change in GABAergic intracortical inhibition was observed when atDCS was applied concomitant to motor training (Amadi et al., 2015; Maceira-Elvira et al., 2022). An interpretation for the absence of such a change could be that LTP-like plasticity is not being induced within M1, or like Amadi and colleagues propose (Amadi et al., 2015), that the LTP induced by atDCS competes with the one induced by motor training itself, leaning towards a state of long-term depression (LTD). If this latter interpretation were correct and the LTP-like plasticity inducing effect of both processes cancelled out, it could explain why we do not see any effects of stimulation on the actual execution of the movements (e.g., on the speed). In which case the question remains: where does the observed effect on the accuracy come from?

We have proposed that atDCS streamlines the transmission of task-relevant information upstream within the motor network. This seems to be limited to the early stages of acquisition, which is assumed to be dominated by processes involving the mapping of stimuli to responses and the storage of spatial coordinates in memory. In addition, we propose that being able to discriminate optimal from suboptimal learners based on the performance of two different motor sequences suggests optimal learners are able to integrate task-general, sequence-unspecific information in a short amount of time. This task-general information would be used to build an internal model of the task, which would be orchestrated by the cerebellum and the basal ganglia (Shadmehr & Krakauer, 2008). These structures are far from the site of stimulation (i.e., M1), so it is unlikely the induced electric field would be able to directly reach them with sufficient intensity. However, Harvey and Svoboda (Harvey & Svoboda, 2007) showed that a “crosstalk” effect exists, in which LTP at one synapse influences the plasticity threshold at neighboring synapses. Depending on the stage of acquisition, this threshold shift could result in LTP being induced at synapses leading upstream of the motor network; with stimulation acting on metaplasticity (i.e., altering the “plasticity of synaptic plasticity” (Abraham & Bear, 1996)), it could facilitate the relay of information and prepare the system to encode the incoming information more efficiently (Hulme et al., 2013). Please note that in Harvey and Svoboda’s study (Harvey & Svoboda, 2007), subthreshold stimulation did not induce LTP; it was only when combined with the LTP crosstalk from the suprathreshold stimulation that synaptic efficiency increased. In the context of our study, it would be through the combination of atDCS (which is subthreshold) and motor training-derived changes in synaptic efficiency, that LTP would be induced. The need for this aggregated interaction would also explain why the induced electric field, spanning across a broad area, has such specific effects.

The differential effect of atDCS could then be related to the native metaplastic properties of the motor network. Aberrant metaplasticity has been related to diminished learning abilities, with its emergence being associated to substance addiction (e.g., ethanol (Bernier et al., 2011)) and neurological disorders (e.g., dystonia (Hulme et al., 2013)). As such, atDCS could help restore metaplasticity to a more normal state. On the other side of the spectrum, the most efficient learners, possibly possessing highly efficient brain networks, could be driven off balance by the external input.

### The specificity of atDCS and its implications for the advancement of the application of neuromodulation techniques

The present findings extend the previous interpretation of the effects of atDCS in acquiring the finger-tapping task, suggesting it facilitates the storage of the spatial coordinates of the task in memory at the early stages of training (Maceira-Elvira et al., 2022). This benefit appears to be reserved for individuals with less efficient neural systems, characterized by an initially low accuracy at relatively low speed.

We did not find evidence for additional effects of stimulation related to a more prolonged exposure to this technique in any of the groups. Under the present hypothesis, in which we propose stimulation can have a restorative effect on individuals with seemingly diminished learning mechanisms, and considering the clear deterioration of the mechanisms supporting other aspects of skill acquisition, such as offline learning (Spencer et al., 2007; Yan et al., 2010), this is perhaps unexpected. However, it may be that the motor cortex is not the best target to enhance this component, or that applying stimulation during practice fails to target the mechanisms supporting offline learning.

M1 has been reported to be a hub where the influx of multimodal sensory information (e.g., somatosensory and visual, (Cross et al., 2021)) converges to be relayed to higher brain areas, and it has been found to play a protagonist role in motor skill acquisition primarily at the early stages of training (Kawai et al., 2015; Picard et al., 2013). Our results show that applying atDCS over M1 enables the rapid improvement of the accuracy towards a stable state, thought to indicate the storage of the elements of the sequence (i.e., the spatial coordinates) in memory (Shmuelof et al., 2012). Previous research shows the spatial coordinates are stored upstream of the motor network, with neural representations emerging in the premotor and parietal cortices (Kornysheva & Diedrichsen, 2014; Yokoi & Diedrichsen, 2019). The lack of such a representation within M1 (Yokoi & Diedrichsen, 2019), and the lack of an effect of stimulation on the speed of execution of the sequences (Maceira-Elvira et al., 2022), suggests atDCS applied at M1 is neither influencing the organization of task-relevant information nor the execution of actions. Instead, a possible explanation is that atDCS is facilitating the sampling of task-relevant information to be relayed back upstream of the motor network. If this were the case, acting on other stages of motor skill acquisition would require targeting alternative brain areas.

The diagram on *Figure 6* illustrates the acquisition time course for the finger-tapping task, with different stages associated to different brain regions. This diagram draws on the stages of acquisition proposed within the Cognitive Framework for Sequential Motor Behavior (C-SMB) proposed by Verwey and colleagues (Verwey et al., 2015), although the interpretation offered in the present discussion differs from the one proposed originally by the authors. At the early stages of training, individuals need to integrate information on the properties of the system they are interacting with. Aspects such as the correspondence of the elements of the sequence to each button, the stiffness of the buttons to be pressed and the spacing between them, need to be sampled and used to generate an internal model for the task (Kawato, 1999). This information is general to the task and unspecific to any particular sequence, indicating that the model generated while performing a certain sequence would, in principle, be applicable to different sequences.

**Fig. 6.**
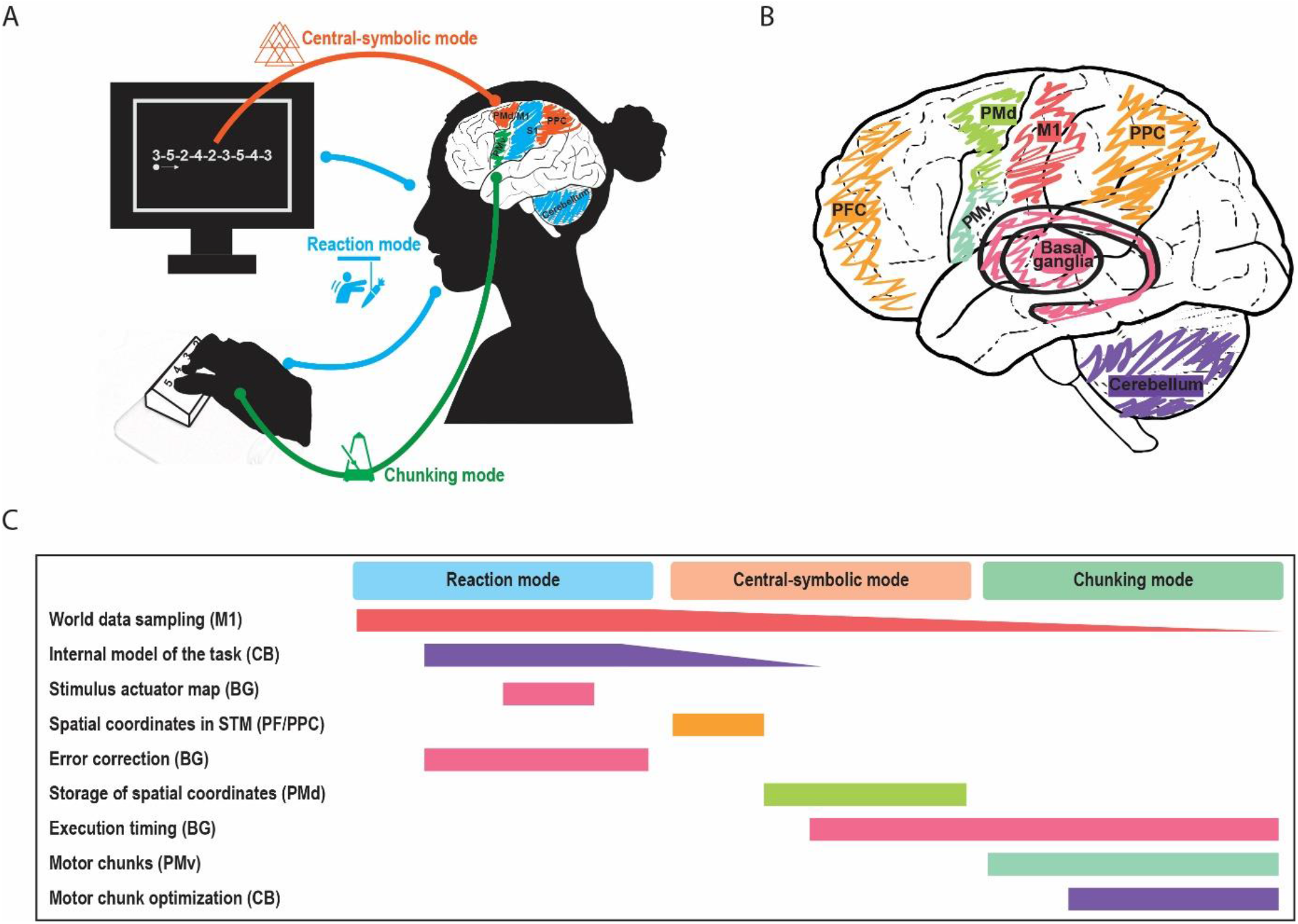
Skill acquisition time course for the finger-tapping task. **(A)** Model for the different steps involved in the acquisition and optimization of the finger-tapping task, based on the stages described within the cognitive framework for sequential motor behavior (C-SMB) (Verwey et al., 2015). **(B)** Brain structures attributed with distinct functions in the acquisition of the finger-tapping task. **(C)** Time line for the acquisition of the finger-tapping task, with distinct stages attributed to different neural structures. The fading bars in the Gantt chart (e.g., world data sampling) indicate processes that remain active but become less dominant with the improvement of skill. Please note that the colors in **(B)** are mapped to those in **(C)**.

Then, participants need to establish a map based on the correspondence between the numerical elements of the sequence and the buttons assigned to each number. In addition, each button is assigned to a different actuator (i.e., each finger of the hand used for training), which ultimately maps each finger to the corresponding numerical element of the sequence. The generation of the model for the task may be supported by the cerebellum, with error-based updates to the model being managed by the basal ganglia (Shadmehr & Krakauer, 2008). In the context of the acquisition of sequential motor tasks, the striatal system has been attributed the role of storing the stimulus-response associations (i.e., the map of numbers to fingers), while the cerebellum is thought to contain the optimal internal model for performing the task (Penhune & Steele, 2012). As such, these regions could constitute potential targets for stimulation in individuals presenting a deficient implementation of these cognitive elements. However, the timing of the stimulation would be crucial, as its effect would depend on the stage of acquisition in which individuals found themselves. For example, the enhancement of plasticity within the cerebellum may improve skill acquisition when reaching a stage at which the role of M1 is less critical, as increased excitability of the cerebellar cortex can lead to decreased heterosynaptic plasticity within M1 (Popa et al., 2013). Indeed, atDCS applied to the cerebellum has been found to be detrimental to the accuracy (Ballard et al., 2019), the speed (Voegtle et al., 2022) and action selection (Jongkees et al., 2019) during early acquisition. Stimulating deeper brain structures, such as the striatum, would be difficult using atDCS, as this and other non-invasive stimulation techniques (such as TMS) present a marked depth-focality tradeoff (Deng et al., 2013). However, new techniques like temporal interference (TI) electrical stimulation show promise in focally modulating deep structures like the hippocampus (Violante et al., 2022) and the striatum (Vassiliadis et al., 2022; Wessel et al., 2022), with encouraging results to enhance the acquisition of sequential motor tasks (Wessel et al., 2022).

After establishing this map, neural resources previously dedicated to choosing an adequate response to each numerical stimulus are likely released and become available for other purposes. As individuals transition from generating individual responses to each presented stimulus (i.e., “Reaction mode”) to performing the sequence from memory as a result of storing the spatial coordinates (i.e., “Central-symbolic mode”), they rely less on sensory feedback (Yeo et al., 2016). A deficient storage of the elements of the sequence in short-term memory could be potentially improved by targeting the prefrontal and parietal cortices. Indeed, transcranial alternate current stimulation (tACS) applied to these regions has reportedly had beneficial effects to improve short-term memory (Reinhart & Nguyen, 2019).

The repeated execution of the sequence results in the emergence of motor chunks (Acuna et al., 2014; Maceira-Elvira et al., 2022; Sakai et al., 2003; Verwey, 2001). The structure of motor chunks depends on the ease of transition between subsequent key presses (Popp et al., 2020), resulting in a pattern that is gradually optimized to balance motor efficacy and computational complexity (Ramkumar et al., 2016). The striatum has been attributed a critical role in the timely generation of actions, playing a major role in the generation of motor chunks (Penhune & Steele, 2012), which would make it a likely target for individuals with a diminished ability to generate efficient chunking patterns. Finally, the absence of offline learning could be targeted by stimulating the hippocampus, as it has been shown to play a major role in the replay of cortical activity during sleep, possibly supporting the consolidation of learning (Albouy et al., 2015; Buch et al., 2021).

### The future of non-invasive brain stimulation

Non-invasive brain stimulation shows promise for the enhancement of failing motor (Zimerman et al., 2013; Zimerman & Hummel, 2010) and cognitive (Indahlastari et al., 2021; Martins et al., 2017; Perceval et al., 2016) functions in healthy older individuals, thus constituting an exciting option for therapeutic and rehabilitative applications (Alonzo et al., 2019; Hummel & Cohen, 2005; Wessel et al., 2015). However, the corpus of literature involving the use of NIBS for this intent report highly variable results (Summers et al., 2016), obstructing the quantification and estimation of the potential benefits of these techniques. The present study provides an illustration of what the deployment of stimulation techniques such as atDCS could look like in a clinical setting. In such a scenario, healthcare providers could assess the likelihood of an individual to benefit from stimulation based on a set of parameters related to the mechanism to be treated (e.g., rehabilitation, memory), which will necessitate the prior identification of such parameters. Here, we have identified a small set of parameters that may jointly represent the ability of an individual to integrate task-relevant information in a well-delimited context, constituted by the finger-tapping task. However, additional investigations are necessary to identify parameters that may relate to other aspects of behavior and cognitive functions and that may be influenced by brain stimulation, such that these techniques may be used to their full potential for a personalized, therapeutic approach.

## Materials and Methods

### Participants

The present work includes data from a total of 153 healthy volunteers. Forty healthy adults volunteered specifically for this study, while the rest were volunteers of previous studies. We grouped them, according to their age, as middle-aged (50-65 y/o; n = 20, 11 female; age*μ* = 59.1 y/o), and older (>65 y/o; n = 20, 10 female; age*μ* = 71.7 y/o). They were randomly assigned within each age group to receive either real stimulation (i.e., verum; middle-aged = 10, age*μ* = 58.9; older = 10, age*μ* = 71.4) or placebo stimulation (middle-aged = 10, age*μ* = 59.2; older = 10, age*μ* = 72.1). The in-depth analyses we implemented for the characterization and classification of individuals as a function of their initial performance in the finger-tapping task was built on a previously-published dataset of 113 individuals (4). These participants were also grouped according to their age as young (18 to 30 years old; n = 41, 27 female; ageμ = 24.5 years old), middle-aged (50 to 65 years old; n = 34, 20 female; ageμ = 57.7 years old), and older (>65 years old; n = 38, 21 female; ageμ = 72.3 years old) adults.

All participants were right-handed, confirmed using the Edinburgh Handedness Inventory (70). The participants reported not having a previous history of serious medical conditions (General Health Questionnaire, GHQ) or contraindications for tDCS (questionnaire based on safety recommendations for these techniques (71)). We performed a neurological examination on all participants to ensure they were healthy and performed the Mini-Mental State Exam (MMSE, (72)) to verify that all participants scored at least 26 out of 30 points. All participants gave their informed consent under protocol guidelines approved by the cantonal ethics committee Vaud, Switzerland (project No. 2017-00301), according to the Declaration of Helsinki. All participants completed training except for one older adult, who chose to leave the study after six days of training for finding the steps involved in setting up the stimulation and motor training to be too overwhelming.

### Motor training

We used the same finger-tapping task as in our previous study (4), requiring participants to replicate a nine-digit sequence as fast and as accurately as possible (Figure 7B). The participants trained their left (non-dominant) hand for 20 min each day for two sets of five consecutive days (i.e., total ten days), with two rest days in between (Figure 7C). Each training session consisted of six 90-second training blocks and one 90-second “catch block” (presenting a sequence different from the training sequence) halfway through training to test for sequence-unspecific learning.

**Fig. 7.**
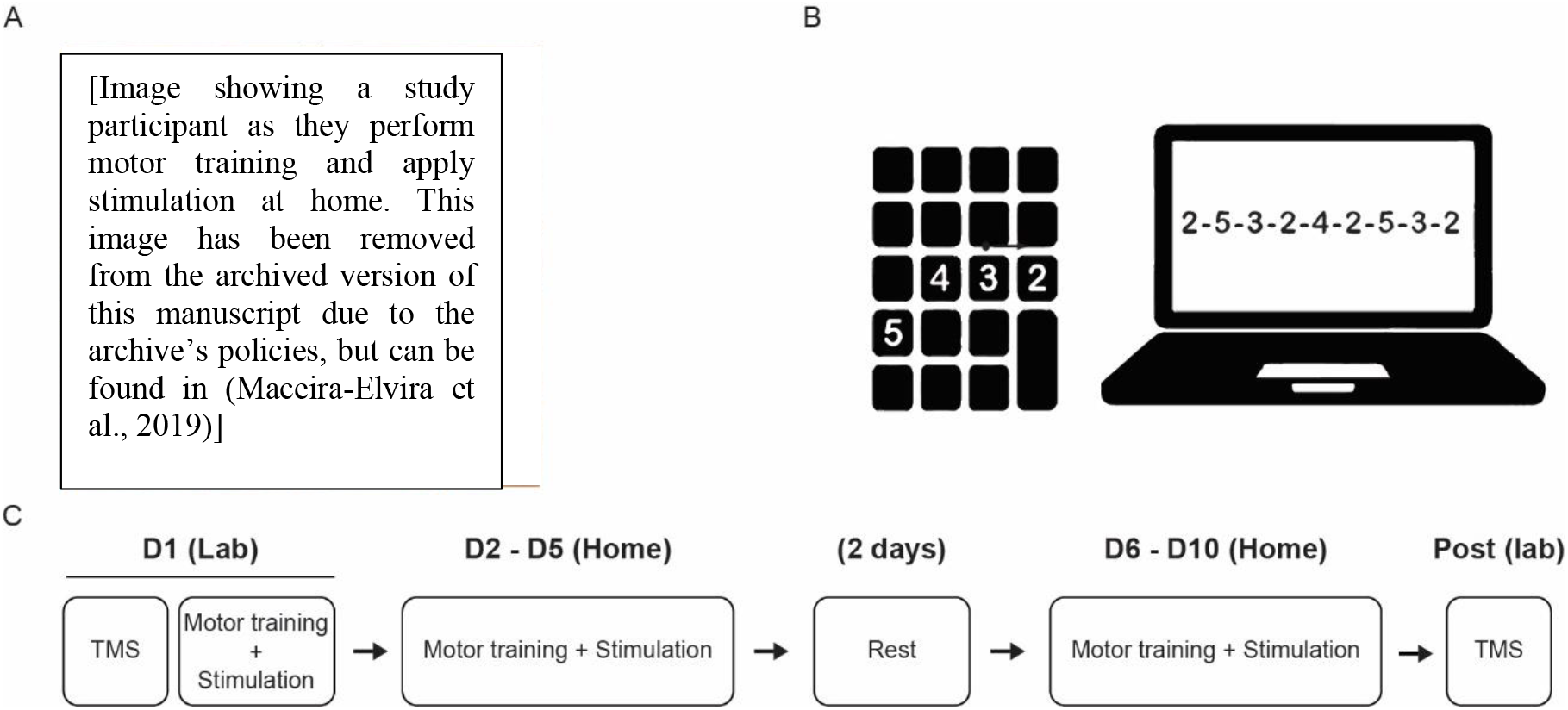
Experimental design. **(A)** Study participant training at home. **(B)** Sequence-tapping task, performed on a laptop using a portable numpad keyboard. **(C)** Training structure. On the first training day, we started by performing neurophysiological investigations using the SICI paradigm, assumed to reflect GABAergic intracortical inhibition within the motor cortex. Immediately after, we provided a thorough explanation on how to prepare the equipment for motor training and stimulation, as detailed in our previous report (Maceira-Elvira et al., 2019). Participants performed the first training session at our research laboratory and the remaining nine days of training independently at home. After the tenth day of training, participants returned to our laboratory for post-training TMS measurements.

### Electrical stimulation of M1 during training

We applied stimulation using the setup we described in an early report of this study (73), with the anode and the cathode placed at the C4 and FP1 locations respectively, according to the 10-20 EEG system. The verum stimulation consisted of 20 min of stimulation with 1 mA direct current, while the placebo stimulation consisted of 32 seconds of stimulation delivered at the beginning of training, with ramp-up/down times of 8 seconds in each case.

### Electrophysiology measurements

We conducted a series of electrophysiological investigations to study intracortical inhibition and its modulation during movement preparation using the well-established SICI paradigm (25–27, 74, 75) in transcranial magnetic stimulation (TMS). We performed measurements at rest and during a reaction time task before and after training to look for training-and/or stimulation-related changes in SICI, using the same protocol as in our previous study (4), targeting the FDI muscle of the left hand. We measured the TMS motor evoked potential using EMG, adjusting the test pulse intensity to 120% resting motor threshold (RMT) and the conditioning pulse intensity to 80% RMT. During the reaction-time task, we applied pulses at 20% and at 90% of the time window between the visual cue and the average reaction time.

### Experimental design

The study followed a randomized, parallel, placebo-controlled, double-blinded design. To facilitate extensive training, taking place over the course of ten days, we asked participants to perform their training with concomitant stimulation at home (Figure 7A), for which we provided all the necessary equipment. The participants came for a first visit, during which we explained how to setup the stimulation and launch the training at home, providing a detailed manual describing the whole procedure (please refer to the Supplementary Materials to find this manual). Participants performed the first training session at our lab, to ensure they had understood the steps involved, and completed the rest of training (i.e., nine days) at home (Figure 7C).

### Data analysis

We used the same analytical pipeline we implemented for our previous study (4) to characterize the execution of the sequence for every participant, extracting the temporal execution patterns (i.e., chunking patterns (8–10)) and quantifying their efficiency. We defined the efficiency of execution in terms of how similar the chunking patterns were to those seen in healthy young adults in our previous study (4) considering young-like chunking patterns to be more efficient.

All statistical tests mentioned below were performed in R (76), using the lme4 package (77) to fit LME models to our data, and the emmeans toolbox (78) for post hoc testing. We obtained the effect sizes from emmeans, and fitted all models using restricted maximum likelihood (REML). We tested the significance of fixed effects by means of ANOVA Type III on the model using Satterthwaite’s method, and obtained p-values using the lmerTest package (79). We performed post hoc tests on significant fixed effects and corrected for multiple comparisons using Tukey’s HSD method (80). We ran two-tailed post hoc tests on the estimated marginal means (i.e., least-squares means) from our fitted models, with degrees of freedom estimated using the Kenward-Roger method (81). The present manuscript discusses, with a few exceptions, significant results only (with a cutoff for statistical significance of p < 0.05). Please refer to the Report on Statistics for the results of all statistical tests.

### Optimal/suboptimal learner stratification

The classification of participants as either “optimal” or “suboptimal” learners was based on our previously proposed model (4), in which the optimal acquisition of the finger-tapping task relies on the prioritization of the improvement of the accuracy at the early stages of training. Based on our observations in various cohorts of healthy young adults, we propose that the optimal acquisition of the finger-tapping task is characterized by the stabilization of the accuracy (i.e., reaching an accuracy plateau) by the end of the first training session. Under this hypothesis, we would consider individuals reaching this plateau on the first training day to be optimal learners, and those reaching the plateau at a later stage to be suboptimal. Following this rationale, we trained a classifier to predict whether participants would reach a plateau in accuracy on the first training day or not based on baseline parameters. Specifically, we used the individual’s age, and their speed and accuracy in the baseline block (performed before training) and the first block of training. Please note that even though we propose age is not the only determining factor in an individual’s ability to acquire new motor skills, we think there are relevant implications to having a certain speed and a certain accuracy, relative to one’s age. In other words, we think there are important differences in being relatively slow and inaccurate at a younger age, compared to being similarly slow and inaccurate at an advanced age. We determined the time point (i.e., the training session) when participants reached a plateau in accuracy by fitting a logarithmic function to the accuracy and detecting the bending point of the curve (please see the Supplementary Materials for an illustration). We assigned a label of “optimal” to participants whenever a bending point was detected on the first training day, and a label of “suboptimal” otherwise.

We trained the model exclusively on data from a previous study (4), and we included only participants that had either received no stimulation or placebo stimulation (i.e., we excluded the participants receiving verum stimulation from the training set). We applied a 2nd degree polynomial expansion to add non-linear relationships among the features, and we used 10-fold cross-validation to select the best model parameters, using 80% of the data for training and 20% for testing in each iteration. After the 10 iterations, we selected the model yielding the highest F1-score, which was a support-vector classifier. After training, we applied the classifier to the training set and extracted the distance of each participant to the hyperplane separating the “optimal” and “suboptimal” classes, and projected these distances on a logistic regression (i.e., sigmoid) function. The sigmoid function has a range between 0 and 1, inclusive, and the interpretation for classification is that values above 0.5 belong to one class and the those below belong to a different class. In the context of our application, participants whose estimated distance from the hyperplane was smaller than 0.5 when being projected onto the sigmoid function were classified as optimal learners, while those above 0.5 were suboptimal. After defining this map, we applied the classifier to the rest of the data (i.e., the data of the present study, with participants receiving either placebo or verum stimulation, and the data of the verum groups in the published dataset) and labeled all participants as either “optimal” and “suboptimal” learners. Further, we segmented each class based on the median of the projected distances, using only the training data. As a result, the optimal and suboptimal classes were split into a higher and a lower tier of learners within each class.

## Supporting information

Report on statistical tests

## Acknowledgments

We would also like to thank the NMOD platform at the Campus Biotech Foundation in Geneva (FCBG) for their support.

## Reporting of results

The reporting of results was done providing all the relevant information recommended in the CONSORT 2010 checklist for the transparent reporting of trials.

## Funding

The project was partially funded by the Defitech Foundation (StS project, Morges, CH), the Bertarelli Foundation -Catalyst program (‘Deep-MCI-T’, Gstaad, CH) and by the SNSF (320030L_197899, NiBS-iCog).

## Author contributions

The co-authors of this work satisfy the recommended requirements offered by the International Committee of Medical Journal Editors (ICMJE), each of which contributing specifically with the elements listed next.

Conception: FCH

Protocol development: PME, TP, ACS, FCH

Data acquisition: PME, TP, ACM, ACS

Data analyses: PME

Interpretation of results: PME, LGC, FCH

Manuscript: PME, FCH

Manuscript critical revision: PME, TP, LGC, FCH

Funding: FCH

## Competing interests

The authors declare no competing interests.

## Data and materials availability

All data needed to evaluate the conclusions in the paper are present in the paper and/or the *Supplementary Materials*. All data necessary to generate the figures within this manuscript and the Supplementary Materials will be available in the Zenodo repository upon peer-reviewed publication. The code written to classify individuals into the four groups of learners will be available in the Zenodo repository upon peer-reviewed publication.

## Notes

### Competing Interest Statement

The authors have declared no competing interest.

