## Supplementary Materials for "Not only a matter of age: Machine learning-based characterization of the differential effect of brain stimulation on skill acquisition"

#### *Behavioral data corrections*

The participants of the present study performed the motor training and applied stimulation at home, as detailed in our previous report (Maceira-Elvira et al., 2019). In a few cases, the participants experienced issues that had to be accounted for before running the analyses detailed in the main text. The problems encountered and the actions we took to account for them are detailed next:

- Participant 217207: The training of this participant was interrupted on two occasions (i.e., on the third block of day 3 and on the third block of day 5) due to stimulation being stopped by the stimulation software. Upon this event, the participant called to get support and was able to resume training. As the score of the blocks in question was affected, we replaced them with the scores of the preceding block (i.e., the second block of the third and the fifth days).
- Participant 217208: On the first block of the first training day, during which the experimenters were present, the participant started training without having connected the keyboard, which resulted in their responses not being recorded by the program. The experimenter noted this problem and added the number of sequences produced while the keyboard was unplugged to the number of sequences detected by the computer. The addition was of three correct sequences.
- Participant 217257: This participant reported doing the last training session (i.e., day 10) while being heavily intoxicated with alcohol. We excluded this training session from the analysis.
- Participant 217309: While performing the training on the third day and during the last two blocks of training, the participant accidentally pressed the “Num Lock” key on the keyboard used to perform the training, which resulted in their

responses not being recorded. These two blocks were excluded from the analysis.

- Participant 217357: While performing the training on the second day and during the first two blocks of training, the participant accidentally pressed the “Num Lock” key on the keyboard used to perform the training, which resulted in their responses not being recorded. These two blocks were excluded from the analysis.

### Task scores and the contribution of online/offline learning to total learning

We quantified the general performance of our participants using the same scoring criteria used in our previous study (Maceira-Elvira et al., 2022). *Figure S1a and S1b* show the scores obtained by our participants, calculated as the number of correct sequences weighted by the ratio of correct to total sequences (i.e., percent correct) generated on each training block. The smaller markers portray individual data, while the larger markers reflect the average for each group, and the error bars represent the standard error of the mean. Please note these scores are not corrected for baseline performance, as we were interested only on the improvement dynamics (i.e., improvement slope) and on the contribution of each acquisition stage to the overall performance change seen towards the end of training.

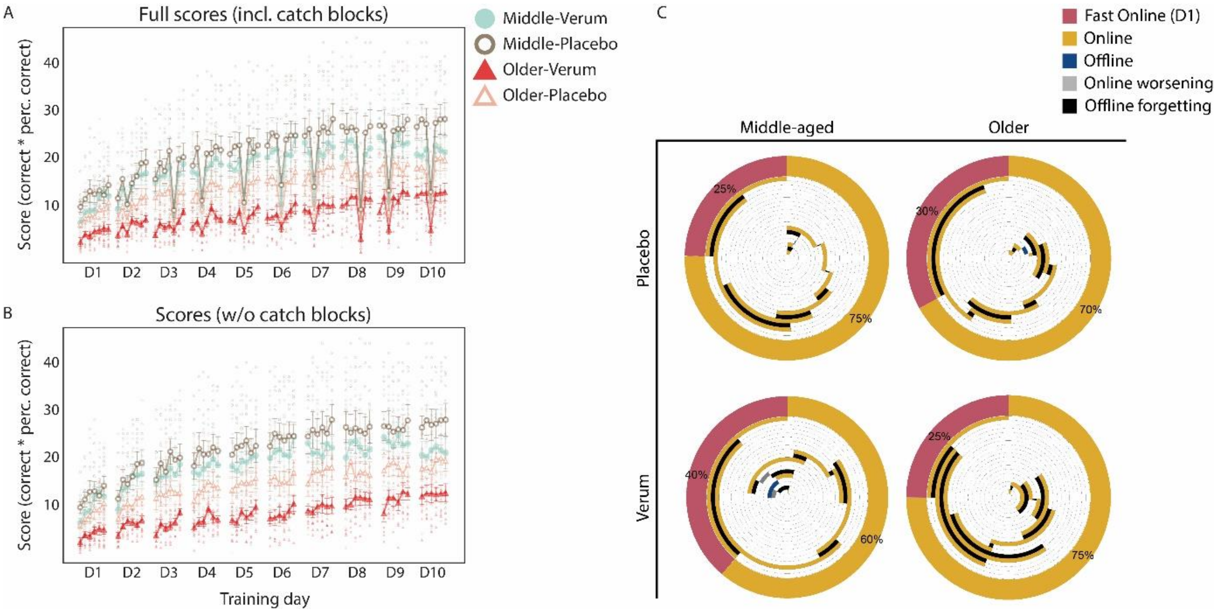


***Fig. S1. Scores obtained by all participants in the finger-tapping task and contribution of the different stages of acquisition to the overall performance change. (A)*** *Participant scores in all blocks within each training session, including the “catch blocks” used to check for generalization of learning. The translucent points depict individual scores, while the opaque points illustrate the mean score of each block in each group. The error bars illustrate the standard error of the mean. Please note the*

*consistent drop in performance within each session, marking the appearance of each catch block.* ***(B)*** *Participant scores in all training blocks, excluding the catch blocks.* ***(C)*** *Contribution of different stages of learning to total learning over the course of the training. The outermost, thickest ring represents total learning, and is segmented to show the percentage of total learning that was derived from performance changes taking place within the first training session (i.e., fast online learning), performance changes taking place during the training sessions from the second day onwards (i.e., online learning), and performance changes taking place overnight (i.e., offline learning). Please note the absence of offline learning in all groups. The inner circles describe the time course of the different stages of acquisition, in which the last ring corresponds to online learning of the first day (thus matching the outer portion assigned to fast online learning), the one-before-last ring corresponds to performance changes taking place the first and second sessions, and so on.*

*Figure S1c* shows the contribution of different stages of acquisition to total learning. Matching our previous findings (Maceira-Elvira et al., 2022), neither middle-aged nor older adults seem to experience overnight improvements (i.e., offline learning); instead, they seem to worsen overnight (offline forgetting). Please note that the amount of offline forgetting in the group of older adults receiving placebo stimulation resembles that seen in both groups of middle-aged adults, while the group of older adults receiving verum stimulation shows a much more dominant presence of offline forgetting.

### “Optimal” and “suboptimal” label creation

The definition we used for optimal and suboptimal learners was based on our previous findings (Maceira-Elvira et al., 2022), in which the groups of individuals acquiring the finger-tapping task most efficiently where able to optimize their accuracy at the early stages of training (i.e., by the end of the first day). Accordingly, we estimated the time point at which individuals reached a plateau in their accuracy. Please note this was not necessarily at 100%; indeed, a stable accuracy at 100% was rare. Rather, the participants seem to settle for an “acceptable” error rate, around which their performance appears to stabilize. Figure S2 shows an example of two participants, with IDs 115104 (Figure S2a) and 115304 (Figure S2b); these participants to a previously published dataset (Maceira-Elvira et al., 2022), in which participants practiced over the course of five consecutive days. In the figure, we show the accuracy values per block of training, as well as a logarithmic function fitted to these values. The red, dashed lines indicate the estimated stabilization value for the accuracy (horizontal), and the block at which the fitted line describes the bend or the “knee” (Satopää et al., 2011) of the fitted line (vertical). The point where these two lines intersect marks the knee point for the fitted function, and constitutes the block at which the accuracy stabilizes. In the context of our interpretation, when this bend occurs within the first 6 training blocks (i.e., the first training day), participants are able to acquire the finger-tapping task most efficiently.

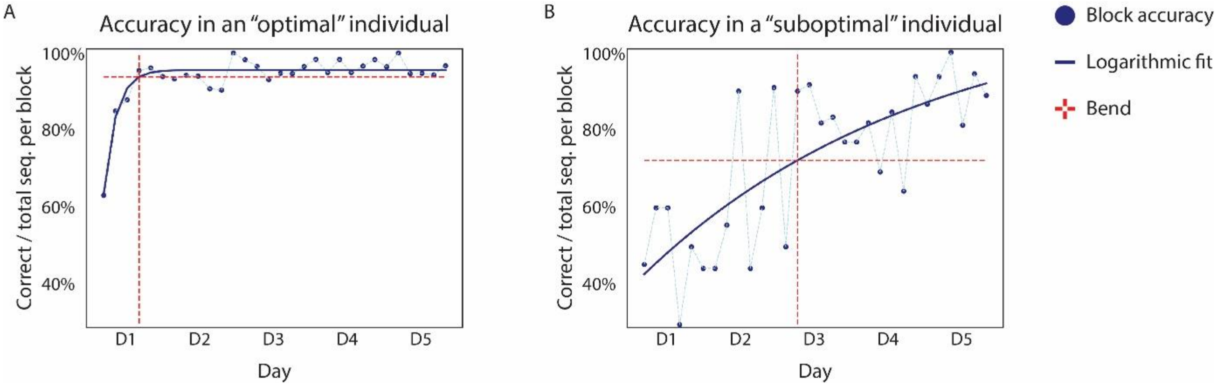


***Figure S2 Label assignment for optimal and suboptimal individuals based on the estimated accuracy stabilization time point.*** *Estimation of the block at which the accuracy stabilizes based on the knee point of a logarithmic function adjusted to the accuracy of each block. The blue dots represent the accuracy on each training block, while the blue, solid line illustrates the fitted function. The red, dashed lines represent the value around which the accuracy was estimated to settle (horizontal) and the training block when this occurred (vertical).* ***(A)*** *Accuracy in an "optimal" learner, whose accuracy stabilized by the end of the first training session.* ***(B)*** *Accuracy in a "suboptimal" learner, whose accuracy stabilized halfway through training.*

### Neurophysiological investigations on GABAergic intracortical inhibition at rest and during movement preparation

*Figure S4* shows the SICI measurements at rest (i.e., SICIrest, (a)) and during movement preparation (i.e., SICImodulation, (b)), averaged per stimulation group. The SICI ratio variable was calculated by dividing the average MEP amplitude obtained when applying the SICI paradigm by the average MEP amplitude obtained when applying the test pulse. The test pulse was delivered at an intensity close to 120% of the resting motor threshold (i.e., RMT), but was adjusted to obtain an MEP amplitude of 1 mV in 5 out of 10 consecutive single pulses. Modulation was calculated as the difference in MEP amplitude when the SICI paradigm was delivered at 90% and 20% of reaction time, as measured during a simple reaction time task.

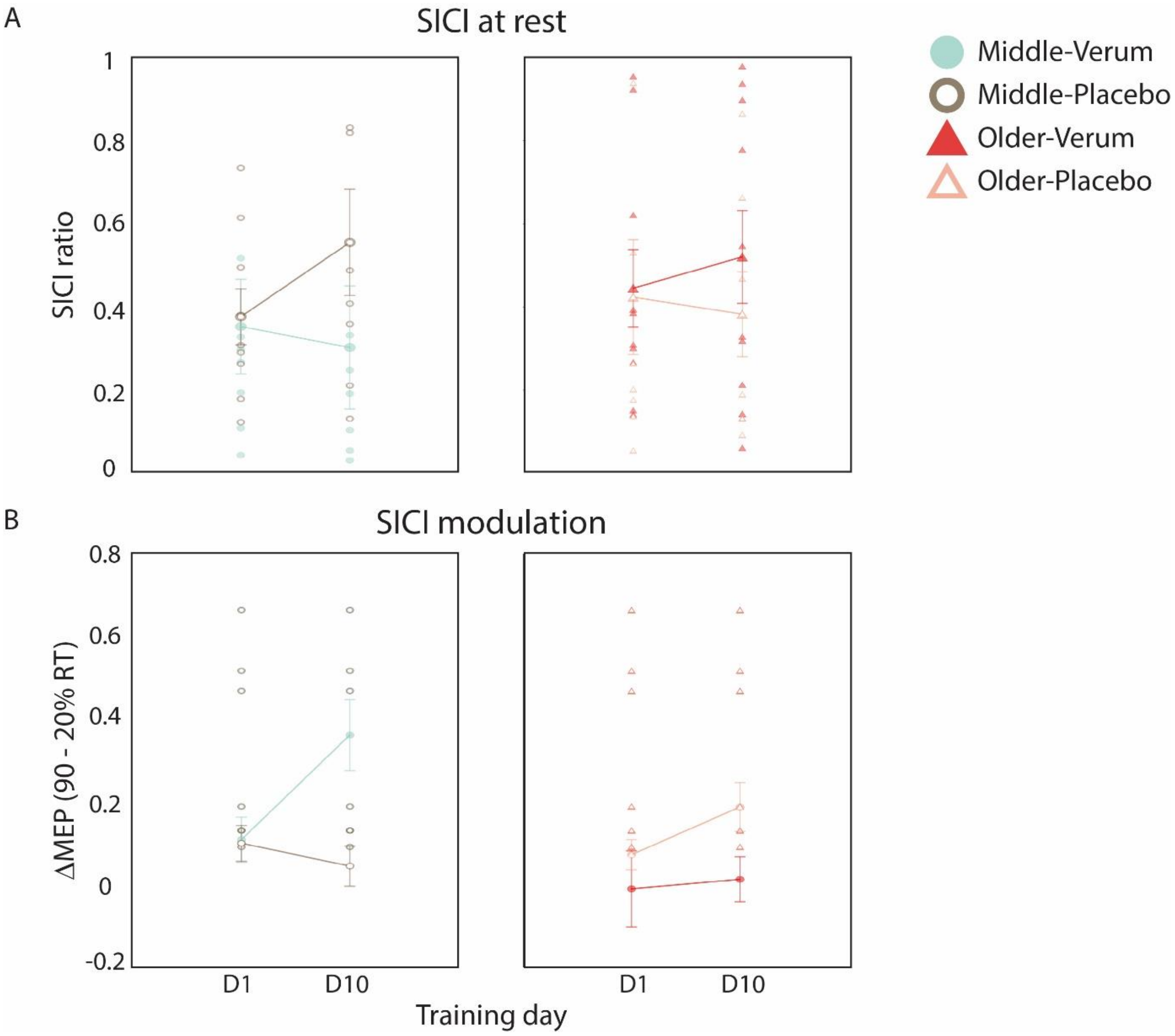


*Fig. S3. **GABAergic intracortical inhibition measured at M1 using the SICI TMS paradigm. (a)** SICI recordings measured while participants were at rest and **(b)** performing a reaction-time task. GABAergic intracortical inhibition was estimated as the ratio between the motor evoked potential (MEP) amplitude in SICI trials to the amplitude obtained when applying the test stimulus alone (calibrated to ~1 mV). Modulation was calculated as the difference in SICI ratio when applying the SICI paradigm at 90% of reaction time with respect to applying the SICI paradigm at 20% of reaction time. The translucent markers depict individual data points, while the solid markers represent the average per age and per stimulation group. The error bars represent the standard error of the mean.*

### Speed dynamics of the learner groups

We found atDCS to have an effect on the accuracy and not on the speed of execution. As further verification, we compared the change in speed (i.e., the speed dynamics) in the four learner categories, for which we reported marked differences in terms of accuracy in the main text. As shown in *Figure S4*, the speed dynamics were similar among groups, with no apparent stimulation-related differences.

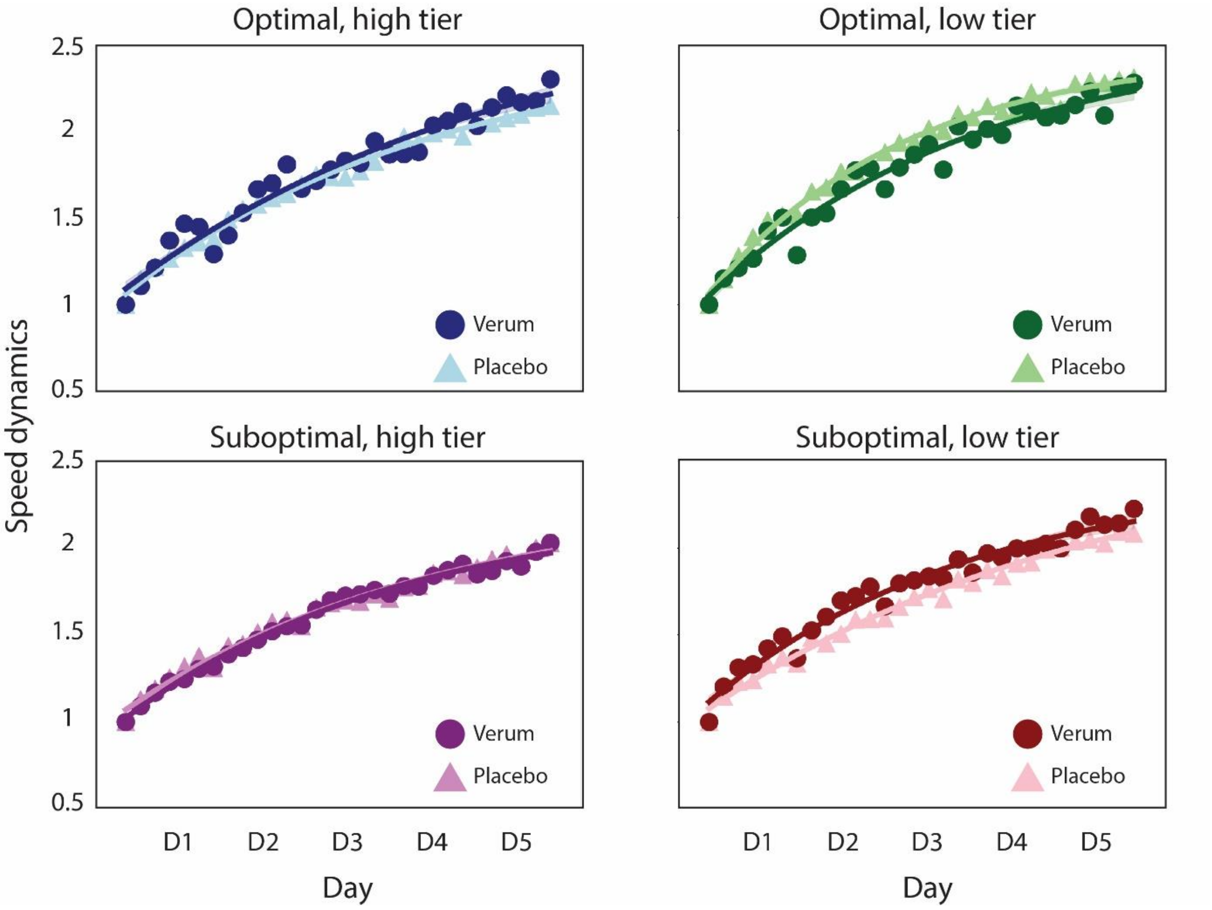


***Fig. S4. Speed dynamics for the different learner tiers.*** *Speed dynamics, calculated as the ratio of the average speed of each training block, divided by the average speed of the first block of training within each group. The data depicted combines the dataset of the present study with our previously-published datasets* (Maceira-Elvira et al., 2022), *so each group contains individuals from all age groups (see Figure 4 and the table in Figure 5 of the main text). Please note the similarity among all learner groups and stimulation groups.*

### Report on statistical testing

As a complement to this document, we have generated a R notebook in HTML format detailing all the statistical tests we conducted, leading to the results detailed in the main text. This notebook is stored in the Zenodo repository (repository url).
