## Supplementary material for "Not only a matter of age: Machine learning-based characterization of the differential effect of brain stimulation on skill acquisition": Report on statistical tests: Differential_stim_effect_Maceira-Elvira_etal_Report_on_statistics.html


Code 

- Show All Code
- Hide All Code
- Download Rmd

### Not only a matter of age: Machine learning-based characterization of the differential effect of brain stimulation on skill acquisition

### Report on Statistics

#### Author: Pablo Maceira-Elvira


```
# Import necessary libraries
library(car)
library(Rmisc)
library(Matrix)
library(dplyr)
library(lme4)
library(lmerTest)
library(emmeans)
library(effectsize)
```


```
#Set Working directory
setwd("C:/Users/maceira/switchdrive/SD_Cosmos/My_documents/Projects/TrainStim/Analyses/1-Statistics/R_code/Main")
curr_dir <- getwd()
```


##### General information

The present report details all the statistical tests performed on the
data discussed in (Maceira-Elvira et al., 202X). For each of the
different parameters of interest (e.g., the speed, the accuracy, etc.),
we present various models of gradually-increasing complexity, seeking
the best representation of the data as quantified using the amount of
variance explained by the model and, when relevant, the Bayesian
Information Criterion (BIC).


```
# Data folder
python_data_directory <- file.path(paste(curr_dir, 'Data', 'AP3_stats', 'MS', sep='/'))
```

##### Speed of execution in the finger-tapping task

The following statistical tests concern the speed of execution in the
finger-tapping task, quantified as the total number of sequences
generated in each training block.


```
speed_file_name <-  "AP3_speed_MS_27_03_2023.txt"
file_path <- file.path(paste(python_data_directory, speed_file_name, sep='/'))
df_speed = read.delim(file_path, header = TRUE, na.strings = "NN")
head(df_speed)
```


```
df_speed$Code <- as.factor(df_speed$Code)
df_speed$AG <- as.factor(df_speed$AG)
df_speed$S <- as.factor(df_speed$S)
df_speed$Group <- as.factor(df_speed$Group)
df_speed$Speed <- as.numeric(df_speed$Speed)
df_speed$D <- as.factor(df_speed$D)
df_speed$B <- as.numeric(df_speed$B)
```


```
Warning: NAs introduced by coercion
```


```
levels(df_speed$Code)
```


```
 [1] "217201" "217202" "217203" "217205" "217206" "217207" "217208" "217209" "217210" "217251" "217252" "217253" "217254" "217255" "217257" "217258" "217259" "217260"
[19] "217301" "217302" "217303" "217304" "217305" "217306" "217307" "217308" "217309" "217310" "217311" "217312" "217351" "217352" "217353" "217354" "217355" "217356"
[37] "217357" "217358" "217359" "217360"
```


```
levels(df_speed$AG)
```


```
[1] "2" "3"
```


```
levels(df_speed$S)
```


```
[1] "1" "2"
```


```
levels(df_speed$Group)
```


```
[1] "Middle-Placebo" "Middle-Verum"   "Older-Placebo"  "Older-Verum"
```


```
levels(df_speed$D)
```


```
 [1] "0"  "1"  "2"  "3"  "4"  "5"  "6"  "7"  "8"  "9"  "10"
```


###### Compare initial training speeds


```
data_subset <- subset(df_speed, D == 1 & B == 1)
data_subset <- droplevels(data_subset)
levels(data_subset$D)
```


```
[1] "1"
```


```
levels(data_subset$Group)
```


```
[1] "Middle-Placebo" "Middle-Verum"   "Older-Placebo"  "Older-Verum"
```


```
m_speed_b1 <- lm(formula = Speed ~ AG*S, data=data_subset)
summary(m_speed_b1)
```


```
Call:
lm(formula = Speed ~ AG * S, data = data_subset)

Residuals:
   Min     1Q Median     3Q    Max 
-6.800 -1.800 -0.100  1.325 10.200 

Coefficients:
            Estimate Std. Error t value Pr(>|t|)    
(Intercept)   9.9000     0.9919   9.981 6.54e-12 ***
AG3          -2.6000     1.4028  -1.853    0.072 .  
S2            1.9000     1.4028   1.354    0.184    
AG3:S2        0.6000     1.9838   0.302    0.764    
---
Signif. codes:  0 ‘***’ 0.001 ‘**’ 0.01 ‘*’ 0.05 ‘.’ 0.1 ‘ ’ 1

Residual standard error: 3.137 on 36 degrees of freedom
Multiple R-squared:  0.2239,    Adjusted R-squared:  0.1593 
F-statistic: 3.462 on 3 and 36 DF,  p-value: 0.02616
```


```
anova(m_speed_b1)
```


```
Analysis of Variance Table

Response: Speed
          Df Sum Sq Mean Sq F value  Pr(>F)  
AG         1   52.9  52.900  5.3766 0.02620 *
S          1   48.4  48.400  4.9193 0.03296 *
AG:S       1    0.9   0.900  0.0915 0.76405  
Residuals 36  354.2   9.839                  
---
Signif. codes:  0 ‘***’ 0.001 ‘**’ 0.01 ‘*’ 0.05 ‘.’ 0.1 ‘ ’ 1
```


```
eta_squared(m_speed_b1)
```


```
# Effect Size for ANOVA (Type I)

Parameter | Eta2 (partial) |       95% CI
-----------------------------------------
AG        |           0.13 | [0.01, 1.00]
S         |           0.12 | [0.01, 1.00]
AG:S      |       2.53e-03 | [0.00, 1.00]

- One-sided CIs: upper bound fixed at [1.00].
```

###### Note:

As there is a significant difference among the age groups, we will
test the difference in speed among the unstimulated cohorts (i.e.,
placebo groups) and the stimulation-related difference within each age
group.

###### Initial speed comparison among the placebo groups


```
data_subset <- subset(df_speed, D == 1 & B == 1 & S == 2)
data_subset <- droplevels(data_subset)
levels(data_subset$D)
```


```
[1] "1"
```


```
levels(data_subset$S)
```


```
[1] "2"
```


```
levels(data_subset$Group)
```


```
[1] "Middle-Placebo" "Older-Placebo"
```


```
m_speed_b1_placebo <- lm(formula = Speed ~ AG, data=data_subset)
summary(m_speed_b1_placebo)
```


```
Call:
lm(formula = Speed ~ AG, data = data_subset)

Residuals:
   Min     1Q Median     3Q    Max 
 -6.80  -1.80  -0.30   1.45  10.20 

Coefficients:
            Estimate Std. Error t value Pr(>|t|)    
(Intercept)   11.800      1.209   9.758  1.3e-08 ***
AG3           -2.000      1.710  -1.170    0.257    
---
Signif. codes:  0 ‘***’ 0.001 ‘**’ 0.01 ‘*’ 0.05 ‘.’ 0.1 ‘ ’ 1

Residual standard error: 3.824 on 18 degrees of freedom
Multiple R-squared:  0.07062,   Adjusted R-squared:  0.01899 
F-statistic: 1.368 on 1 and 18 DF,  p-value: 0.2574
```


```
anova(m_speed_b1_placebo)
```


```
Analysis of Variance Table

Response: Speed
          Df Sum Sq Mean Sq F value Pr(>F)
AG         1   20.0  20.000  1.3678 0.2574
Residuals 18  263.2  14.622
```


```
eta_squared(m_speed_b1_placebo)
```


```
For one-way between subjects designs, partial eta squared is equivalent to eta squared. Returning eta squared.
```


```
# Effect Size for ANOVA

Parameter | Eta2 |       95% CI
-------------------------------
AG        | 0.07 | [0.00, 1.00]

- One-sided CIs: upper bound fixed at [1.00].
```


```
t.test(formula = Speed ~ AG, data=data_subset)
```


```
    Welch Two Sample t-test

data:  Speed by AG
t = 1.1695, df = 17.852, p-value = 0.2576
alternative hypothesis: true difference in means between group 2 and group 3 is not equal to 0
95 percent confidence interval:
 -1.594931  5.594931
sample estimates:
mean in group 2 mean in group 3 
           11.8             9.8
```


```
cohens_d(Speed ~ AG, data=data_subset)
```


```
Cohen's d |        95% CI
-------------------------
0.52      | [-0.38, 1.41]

- Estimated using pooled SD.
```

###### Initial speed comparison among the stimulation groups in middle-aged adults


```
data_subset <- subset(df_speed, D == 1 & B == 1 & AG == 2)
data_subset <- droplevels(data_subset)
levels(data_subset$D)
```


```
[1] "1"
```


```
levels(data_subset$AG)
```


```
[1] "2"
```


```
levels(data_subset$S)
```


```
[1] "1" "2"
```


```
levels(data_subset$Group)
```


```
[1] "Middle-Placebo" "Middle-Verum"
```


```
m_speed_b1_middle <- lm(formula = Speed ~ S, data=data_subset)
summary(m_speed_b1_middle)
```


```
Call:
lm(formula = Speed ~ S, data = data_subset)

Residuals:
   Min     1Q Median     3Q    Max 
-6.800 -1.825 -0.400  2.200  5.200 

Coefficients:
            Estimate Std. Error t value Pr(>|t|)    
(Intercept)    9.900      1.023   9.674 1.48e-08 ***
S2             1.900      1.447   1.313    0.206    
---
Signif. codes:  0 ‘***’ 0.001 ‘**’ 0.01 ‘*’ 0.05 ‘.’ 0.1 ‘ ’ 1

Residual standard error: 3.236 on 18 degrees of freedom
Multiple R-squared:  0.08739,   Adjusted R-squared:  0.03669 
F-statistic: 1.724 on 1 and 18 DF,  p-value: 0.2057
```


```
anova(m_speed_b1_middle)
```


```
Analysis of Variance Table

Response: Speed
          Df Sum Sq Mean Sq F value Pr(>F)
S          1  18.05  18.050  1.7236 0.2057
Residuals 18 188.50  10.472
```


```
eta_squared(m_speed_b1_middle)
```


```
For one-way between subjects designs, partial eta squared is equivalent to eta squared. Returning eta squared.
```


```
# Effect Size for ANOVA

Parameter | Eta2 |       95% CI
-------------------------------
S         | 0.09 | [0.00, 1.00]

- One-sided CIs: upper bound fixed at [1.00].
```


```
t.test(formula = Speed ~ S, data=data_subset)
```


```
    Welch Two Sample t-test

data:  Speed by S
t = -1.3129, df = 16.786, p-value = 0.2069
alternative hypothesis: true difference in means between group 1 and group 2 is not equal to 0
95 percent confidence interval:
 -4.956339  1.156339
sample estimates:
mean in group 1 mean in group 2 
            9.9            11.8
```


```
cohens_d(Speed ~ S, data=data_subset)
```


```
Cohen's d |        95% CI
-------------------------
-0.59     | [-1.48, 0.32]

- Estimated using pooled SD.
```

###### Initial speed comparison among the stimulation groups in older adults


```
data_subset <- subset(df_speed, D == 1 & B == 1 & AG == 3)
data_subset <- droplevels(data_subset)
levels(data_subset$D)
```


```
[1] "1"
```


```
levels(data_subset$AG)
```


```
[1] "3"
```


```
levels(data_subset$S)
```


```
[1] "1" "2"
```


```
levels(data_subset$Group)
```


```
[1] "Older-Placebo" "Older-Verum"
```


```
m_speed_b1_older <- lm(formula = Speed ~ S, data=data_subset)
summary(m_speed_b1_older)
```


```
Call:
lm(formula = Speed ~ S, data = data_subset)

Residuals:
   Min     1Q Median     3Q    Max 
-4.800 -1.425 -0.050  0.700 10.200 

Coefficients:
            Estimate Std. Error t value Pr(>|t|)    
(Intercept)   7.3000     0.9595   7.608 4.97e-07 ***
S2            2.5000     1.3569   1.842   0.0819 .  
---
Signif. codes:  0 ‘***’ 0.001 ‘**’ 0.01 ‘*’ 0.05 ‘.’ 0.1 ‘ ’ 1

Residual standard error: 3.034 on 18 degrees of freedom
Multiple R-squared:  0.1587,    Adjusted R-squared:  0.1119 
F-statistic: 3.395 on 1 and 18 DF,  p-value: 0.08194
```


```
anova(m_speed_b1_older)
```


```
Analysis of Variance Table

Response: Speed
          Df Sum Sq Mean Sq F value  Pr(>F)  
S          1  31.25 31.2500  3.3947 0.08194 .
Residuals 18 165.70  9.2056                  
---
Signif. codes:  0 ‘***’ 0.001 ‘**’ 0.01 ‘*’ 0.05 ‘.’ 0.1 ‘ ’ 1
```


```
eta_squared(m_speed_b1_older)
```


```
For one-way between subjects designs, partial eta squared is equivalent to eta squared. Returning eta squared.
```


```
# Effect Size for ANOVA

Parameter | Eta2 |       95% CI
-------------------------------
S         | 0.16 | [0.00, 1.00]

- One-sided CIs: upper bound fixed at [1.00].
```


```
t.test(formula = Speed ~ S, data=data_subset)
```


```
    Welch Two Sample t-test

data:  Speed by S
t = -1.8425, df = 11.706, p-value = 0.09086
alternative hypothesis: true difference in means between group 1 and group 2 is not equal to 0
95 percent confidence interval:
 -5.4646309  0.4646309
sample estimates:
mean in group 1 mean in group 2 
            7.3             9.8
```


```
cohens_d(Speed ~ S, data=data_subset)
```


```
Cohen's d |        95% CI
-------------------------
-0.82     | [-1.73, 0.10]

- Estimated using pooled SD.
```

###### Build models to compare the speed over the course of training

For these tests, we are interested in the rate of improvement (i.e.,
the slopes), which is why we are not using corrected (i.e., centered)
data. We are not so interested in comparing the speed of the
participants in absolute terms, as we have found stimulation not to have
a direct effect on the speed of execution (please see Maceira-Elvira et
al., 2022. Sci. Adv.). However, stimulation may have an indirect effect
on speed, as it enables the early storage of the spatial coordinates in
memory, which we detect as the stabilization of the accuracy (i.e.,
reaching a plateau). Our previous findings show that once accuracy
stabilizes, participants can focus on improving speed, and a higher
prime placed on speed drives the early emergence of motor chunks, which
in turn result in more pronounced speed increases (i.e., steeper
slopes). For this reason, here we build linear models on the uncentered
data and focus on the rate of performance change over time. This is also
why we do not build models gradually, starting off from simple models
(e.g., including only the group averages). Instead, we include the
blocks directly to fit lines to each training day. However, we will
still assess whether we should include random effects or not.


```
data_subset <- subset(df_speed, D != 0)
data_subset <- na.omit(data_subset)
data_subset <- droplevels(data_subset)
levels(data_subset$D)
```


```
 [1] "1"  "2"  "3"  "4"  "5"  "6"  "7"  "8"  "9"  "10"
```


```
levels(data_subset$Group)
```


```
[1] "Middle-Placebo" "Middle-Verum"   "Older-Placebo"  "Older-Verum"
```


```
m1 <- lm(formula = Speed ~ AG*S*D*B, data=data_subset)
```

###### Include random intercepts per subject


```
m2 <- lmer(formula = Speed ~ AG*S*D*B + (1 | Code), data=data_subset)
```

###### Compare models to justify the inclusion of a random intercept.

We cannot do this comparison through a regular ANOVA, as the function
does not support comparing linear to mixed-effects models. However, we
will use tha Bayes Information Criterion (BIC), thought to be a good
criterion for model selection due to its consideration of sample sizes
(Chakrabarti and Ghosh, 2011).


```
BIC(m1, m2)
```


###### Note:

The model including random intercepts for individual subjects greatly
decreases the magnitude of the BIC, which indicates that the model
explains more of the variability of the data. Next, we will compare this
model to a model including also random slopes. Please note that for this
model, we set the Restricted Maximum Likelihood (REML) parameter to
“False”. The reason is that the model failed to converge when fitting
the model by REML.


```
m3 <- lmer(formula = Speed ~ AG*S*D*B + (1 + B|Code), data=data_subset, REML=FALSE)
```

###### Compare the models.


```
anova(m2, m3)
```


```
refitting model(s) with ML (instead of REML)
```


```
Data: data_subset
Models:
m2: Speed ~ AG * S * D * B + (1 | Code)
m3: Speed ~ AG * S * D * B + (1 + B | Code)
   npar   AIC   BIC  logLik deviance  Chisq Df Pr(>Chisq)   
m2   82 10400 10872 -5117.8    10236                        
m3   84 10392 10876 -5112.1    10224 11.431  2   0.003294 **
---
Signif. codes:  0 ‘***’ 0.001 ‘**’ 0.01 ‘*’ 0.05 ‘.’ 0.1 ‘ ’ 1
```

###### Model selection

Based on the BIC, the model including random slopes (i.e., m3) is
significantly worse than the one including only random intercepts (i.e.,
m2). Therefore, we will keep m2 as the model for speed.


```
m_speed <- lmer(formula = Speed ~ AG*S*D*B + (1 | Code), data=data_subset)
summary(m_speed)
```


```
Linear mixed model fit by REML. t-tests use Satterthwaite's method ['lmerModLmerTest']
Formula: Speed ~ AG * S * D * B + (1 | Code)
   Data: data_subset

REML criterion at convergence: 10315.7

Scaled residuals: 
    Min      1Q  Median      3Q     Max 
-5.9787 -0.5240  0.0138  0.5487  3.5313 

Random effects:
 Groups   Name        Variance Std.Dev.
 Code     (Intercept) 51.216   7.157   
 Residual              4.287   2.070   
Number of obs: 2342, groups:  Code, 40

Fixed effects:
               Estimate Std. Error         df t value Pr(>|t|)    
(Intercept)   9.587e+00  2.344e+00  4.129e+01   4.090 0.000195 ***
AG3          -2.700e+00  3.315e+00  4.129e+01  -0.815 0.419981    
S2            1.713e+00  3.315e+00  4.129e+01   0.517 0.607975    
D2            3.284e+00  8.861e-01  2.226e+03   3.706 0.000216 ***
D3            8.020e+00  8.620e-01  2.226e+03   9.304  < 2e-16 ***
D4            1.115e+01  8.620e-01  2.226e+03  12.931  < 2e-16 ***
D5            1.350e+01  8.620e-01  2.226e+03  15.661  < 2e-16 ***
D6            1.394e+01  8.861e-01  2.226e+03  15.730  < 2e-16 ***
D7            1.581e+01  8.620e-01  2.226e+03  18.345  < 2e-16 ***
D8            1.722e+01  8.620e-01  2.226e+03  19.976  < 2e-16 ***
D9            1.877e+01  8.620e-01  2.226e+03  21.771  < 2e-16 ***
D10           1.838e+01  9.154e-01  2.226e+03  20.082  < 2e-16 ***
B             1.180e+00  1.565e-01  2.226e+03   7.539 6.84e-14 ***
AG3:S2        5.600e-01  4.687e+00  4.129e+01   0.119 0.905485    
AG3:D2       -1.930e+00  1.236e+00  2.226e+03  -1.561 0.118572    
AG3:D3       -5.060e+00  1.219e+00  2.226e+03  -4.151 3.44e-05 ***
AG3:D4       -7.200e+00  1.219e+00  2.226e+03  -5.906 4.04e-09 ***
AG3:D5       -9.033e+00  1.219e+00  2.226e+03  -7.410 1.78e-13 ***
AG3:D6       -7.639e+00  1.253e+00  2.226e+03  -6.096 1.28e-09 ***
AG3:D7       -8.493e+00  1.219e+00  2.226e+03  -6.967 4.25e-12 ***
AG3:D8       -9.240e+00  1.219e+00  2.226e+03  -7.580 5.06e-14 ***
AG3:D9       -9.907e+00  1.219e+00  2.226e+03  -8.126 7.25e-16 ***
AG3:D10      -8.636e+00  1.257e+00  2.226e+03  -6.868 8.41e-12 ***
S2:D2        -5.436e-01  1.236e+00  2.226e+03  -0.440 0.660182    
S2:D3        -6.667e-03  1.219e+00  2.226e+03  -0.005 0.995637    
S2:D4         1.267e-01  1.219e+00  2.226e+03   0.104 0.917255    
S2:D5        -1.800e-01  1.219e+00  2.226e+03  -0.148 0.882630    
S2:D6         2.427e+00  1.236e+00  2.226e+03   1.963 0.049715 *  
S2:D7         1.413e+00  1.219e+00  2.226e+03   1.159 0.246437    
S2:D8         1.340e+00  1.219e+00  2.226e+03   1.099 0.271801    
S2:D9         2.800e-01  1.219e+00  2.226e+03   0.230 0.818359    
S2:D10        1.291e+00  1.257e+00  2.226e+03   1.027 0.304741    
AG3:B        -7.571e-01  2.213e-01  2.226e+03  -3.421 0.000636 ***
S2:B         -1.943e-01  2.213e-01  2.226e+03  -0.878 0.380173    
D2:B         -7.841e-02  2.274e-01  2.226e+03  -0.345 0.730272    
D3:B         -5.343e-01  2.213e-01  2.226e+03  -2.414 0.015867 *  
D4:B         -8.371e-01  2.213e-01  2.226e+03  -3.782 0.000160 ***
D5:B         -1.100e+00  2.213e-01  2.226e+03  -4.970 7.22e-07 ***
D6:B         -7.673e-01  2.274e-01  2.226e+03  -3.374 0.000753 ***
D7:B         -1.109e+00  2.213e-01  2.226e+03  -5.008 5.92e-07 ***
D8:B         -1.149e+00  2.213e-01  2.226e+03  -5.189 2.31e-07 ***
D9:B         -1.357e+00  2.213e-01  2.226e+03  -6.131 1.03e-09 ***
D10:B        -1.173e+00  2.348e-01  2.226e+03  -4.996 6.32e-07 ***
AG3:S2:D2     1.178e+00  1.745e+00  2.226e+03   0.675 0.499692    
AG3:S2:D3     2.847e+00  1.726e+00  2.226e+03   1.650 0.099148 .  
AG3:S2:D4     2.980e+00  1.724e+00  2.226e+03   1.729 0.084036 .  
AG3:S2:D5     3.567e+00  1.724e+00  2.226e+03   2.069 0.038680 *  
AG3:S2:D6     1.673e+00  1.748e+00  2.226e+03   0.957 0.338801    
AG3:S2:D7     1.511e+00  1.736e+00  2.226e+03   0.870 0.384226    
AG3:S2:D8     2.576e+00  1.736e+00  2.226e+03   1.484 0.138012    
AG3:S2:D9     3.327e+00  1.736e+00  2.226e+03   1.916 0.055504 .  
AG3:S2:D10    2.644e+00  1.763e+00  2.226e+03   1.499 0.133919    
AG3:S2:B      4.971e-01  3.130e-01  2.226e+03   1.588 0.112390    
AG3:D2:B      1.584e-01  3.173e-01  2.226e+03   0.499 0.617703    
AG3:D3:B      4.314e-01  3.130e-01  2.226e+03   1.378 0.168268    
AG3:D4:B      7.571e-01  3.130e-01  2.226e+03   2.419 0.015653 *  
AG3:D5:B      1.100e+00  3.130e-01  2.226e+03   3.514 0.000450 ***
AG3:D6:B      6.937e-01  3.216e-01  2.226e+03   2.157 0.031126 *  
AG3:D7:B      9.886e-01  3.130e-01  2.226e+03   3.158 0.001609 ** 
AG3:D8:B      9.971e-01  3.130e-01  2.226e+03   3.185 0.001465 ** 
AG3:D9:B      1.154e+00  3.130e-01  2.226e+03   3.687 0.000232 ***
AG3:D10:B     9.643e-01  3.227e-01  2.226e+03   2.989 0.002834 ** 
S2:D2:B       3.670e-01  3.173e-01  2.226e+03   1.156 0.247636    
S2:D3:B       1.971e-01  3.130e-01  2.226e+03   0.630 0.528896    
S2:D4:B       3.400e-01  3.130e-01  2.226e+03   1.086 0.277525    
S2:D5:B       5.514e-01  3.130e-01  2.226e+03   1.762 0.078276 .  
S2:D6:B      -1.841e-02  3.173e-01  2.226e+03  -0.058 0.953737    
S2:D7:B       3.771e-01  3.130e-01  2.226e+03   1.205 0.228402    
S2:D8:B       1.886e-01  3.130e-01  2.226e+03   0.602 0.546963    
S2:D9:B       4.343e-01  3.130e-01  2.226e+03   1.387 0.165469    
S2:D10:B      3.757e-01  3.227e-01  2.226e+03   1.164 0.244379    
AG3:S2:D2:B  -4.078e-01  4.472e-01  2.226e+03  -0.912 0.361988    
AG3:S2:D3:B  -4.952e-01  4.442e-01  2.226e+03  -1.115 0.265113    
AG3:S2:D4:B  -6.371e-01  4.427e-01  2.226e+03  -1.439 0.150219    
AG3:S2:D5:B  -8.857e-01  4.427e-01  2.226e+03  -2.001 0.045540 *  
AG3:S2:D6:B  -5.508e-01  4.488e-01  2.226e+03  -1.227 0.219851    
AG3:S2:D7:B  -6.749e-01  4.458e-01  2.226e+03  -1.514 0.130139    
AG3:S2:D8:B  -7.787e-01  4.458e-01  2.226e+03  -1.747 0.080776 .  
AG3:S2:D9:B  -8.397e-01  4.458e-01  2.226e+03  -1.884 0.059731 .  
AG3:S2:D10:B -9.532e-01  4.526e-01  2.226e+03  -2.106 0.035304 *  
---
Signif. codes:  0 ‘***’ 0.001 ‘**’ 0.01 ‘*’ 0.05 ‘.’ 0.1 ‘ ’ 1
```


```
Correlation matrix not shown by default, as p = 80 > 12.
Use print(x, correlation=TRUE)  or
    vcov(x)        if you need it
```


```
anova(m_speed)
```


```
Type III Analysis of Variance Table with Satterthwaite's method
          Sum Sq Mean Sq NumDF   DenDF  F value    Pr(>F)    
AG          53.4   53.42     1   36.44  12.4616  0.001146 ** 
S           11.5   11.52     1   36.44   2.6867  0.109799    
D        11218.2 1246.47     9 2226.01 290.7594 < 2.2e-16 ***
B          909.0  909.02     1 2226.00 212.0446 < 2.2e-16 ***
AG:S         1.6    1.62     1   36.44   0.3772  0.542905    
AG:D       874.6   97.18     9 2226.01  22.6683 < 2.2e-16 ***
S:D        112.7   12.53     9 2226.01   2.9218  0.001906 ** 
AG:B        15.4   15.43     1 2226.00   3.5999  0.057911 .  
S:B          1.0    1.01     1 2226.00   0.2366  0.626709    
D:B        550.9   61.21     9 2226.00  14.2788 < 2.2e-16 ***
AG:S:D      31.7    3.52     9 2226.01   0.8220  0.595799    
AG:S:B       6.7    6.66     1 2226.00   1.5546  0.212582    
AG:D:B     119.4   13.27     9 2226.00   3.0943  0.001062 ** 
S:D:B       28.8    3.20     9 2226.00   0.7462  0.666679    
AG:S:D:B    30.4    3.38     9 2226.00   0.7877  0.627834    
---
Signif. codes:  0 ‘***’ 0.001 ‘**’ 0.01 ‘*’ 0.05 ‘.’ 0.1 ‘ ’ 1
```


```
eta_squared(m_speed)
```


```
# Effect Size for ANOVA (Type III)

Parameter | Eta2 (partial) |       95% CI
-----------------------------------------
AG        |           0.25 | [0.08, 1.00]
S         |           0.07 | [0.00, 1.00]
D         |           0.54 | [0.52, 1.00]
B         |           0.09 | [0.07, 1.00]
AG:S      |           0.01 | [0.00, 1.00]
AG:D      |           0.08 | [0.06, 1.00]
S:D       |           0.01 | [0.00, 1.00]
AG:B      |       1.61e-03 | [0.00, 1.00]
S:B       |       1.06e-04 | [0.00, 1.00]
D:B       |           0.05 | [0.04, 1.00]
AG:S:D    |       3.31e-03 | [0.00, 1.00]
AG:S:B    |       6.98e-04 | [0.00, 1.00]
AG:D:B    |           0.01 | [0.00, 1.00]
S:D:B     |       3.01e-03 | [0.00, 1.00]
AG:S:D:B  |       3.17e-03 | [0.00, 1.00]

- One-sided CIs: upper bound fixed at [1.00].
```


###### Note:

There are significant differences among the age groups and there is a
significant interaction between days and stimulation. The difference
among the age groups is expected by design and based on our previous
results (Maceira-Elvira et al., 2022. Sci. Adv.) Based on our
hypotheses, we are interested in a series of specific comparisons,
namely: 1. Is there a difference in speed between middle-aged and older
adults in the absence of stimulation? In previous studies, we have found
older adults to be generally slower than middle-aged adults, but the
cohort of older adults in the control condition of the present study
appear to be faster then previous cohorts we have recruited (please see
Figure 1 in the main manuscript). 2. Is there an effect of stimulation
in the rate of change of the accuracy within each group?

###### 1. Compare placebo groups


```
data_subset <- subset(df_speed, D != 0 & S == 2)
data_subset <- na.omit(data_subset)
data_subset <- droplevels(data_subset)
levels(data_subset$D)
```


```
 [1] "1"  "2"  "3"  "4"  "5"  "6"  "7"  "8"  "9"  "10"
```


```
levels(data_subset$S)
```


```
[1] "2"
```


```
levels(data_subset$Group)
```


```
[1] "Middle-Placebo" "Older-Placebo"
```


```
m_speed_placebo <- lmer(formula = Speed ~ AG*D*B + (1 | Code), data=data_subset)
summary(m_speed_placebo)
```


```
Linear mixed model fit by REML. t-tests use Satterthwaite's method ['lmerModLmerTest']
Formula: Speed ~ AG * D * B + (1 | Code)
   Data: data_subset

REML criterion at convergence: 5357.4

Scaled residuals: 
    Min      1Q  Median      3Q     Max 
-5.4915 -0.5490  0.0143  0.5687  3.2442 

Random effects:
 Groups   Name        Variance Std.Dev.
 Code     (Intercept) 69.654   8.346   
 Residual              5.083   2.255   
Number of obs: 1172, groups:  Code, 20

Fixed effects:
              Estimate Std. Error         df t value Pr(>|t|)    
(Intercept)   11.30000    2.72139   20.29794   4.152 0.000480 ***
AG3           -2.14000    3.84862   20.29793  -0.556 0.584259    
D2             2.74000    0.93865 1114.00174   2.919 0.003581 ** 
D3             8.01333    0.93865 1114.00174   8.537  < 2e-16 ***
D4            11.27333    0.93865 1114.00174  12.010  < 2e-16 ***
D5            13.32000    0.93865 1114.00174  14.191  < 2e-16 ***
D6            16.36667    0.93865 1114.00174  17.436  < 2e-16 ***
D7            17.22667    0.93865 1114.00174  18.353  < 2e-16 ***
D8            18.56000    0.93865 1114.00174  19.773  < 2e-16 ***
D9            19.04667    0.93865 1114.00174  20.292  < 2e-16 ***
D10           19.67333    0.93865 1114.00174  20.959  < 2e-16 ***
B              0.98571    0.17043 1114.00174   5.784 9.48e-09 ***
AG3:D2        -0.75220    1.34162 1114.00365  -0.561 0.575140    
AG3:D3        -2.21286    1.33025 1114.00267  -1.663 0.096494 .  
AG3:D4        -4.22000    1.32745 1114.00173  -3.179 0.001518 ** 
AG3:D5        -5.46667    1.32745 1114.00173  -4.118 4.10e-05 ***
AG3:D6        -5.96667    1.32745 1114.00173  -4.495 7.69e-06 ***
AG3:D7        -6.98223    1.34637 1114.00610  -5.186 2.55e-07 ***
AG3:D8        -6.66371    1.34637 1114.00610  -4.949 8.59e-07 ***
AG3:D9        -6.58000    1.34637 1114.00610  -4.887 1.17e-06 ***
AG3:D10       -5.99185    1.34637 1114.00610  -4.450 9.43e-06 ***
AG3:B         -0.26000    0.24102 1114.00174  -1.079 0.280938    
D2:B           0.28857    0.24102 1114.00174   1.197 0.231452    
D3:B          -0.33714    0.24102 1114.00174  -1.399 0.162151    
D4:B          -0.49714    0.24102 1114.00174  -2.063 0.039378 *  
D5:B          -0.54857    0.24102 1114.00174  -2.276 0.023034 *  
D6:B          -0.78571    0.24102 1114.00174  -3.260 0.001148 ** 
D7:B          -0.73143    0.24102 1114.00174  -3.035 0.002464 ** 
D8:B          -0.96000    0.24102 1114.00174  -3.983 7.25e-05 ***
D9:B          -0.92286    0.24102 1114.00174  -3.829 0.000136 ***
D10:B         -0.79714    0.24102 1114.00174  -3.307 0.000972 ***
AG3:D2:B      -0.24933    0.34317 1114.00293  -0.727 0.467648    
AG3:D3:B      -0.06375    0.34322 1114.00510  -0.186 0.852680    
AG3:D4:B       0.12000    0.34086 1114.00173   0.352 0.724865    
AG3:D5:B       0.21429    0.34086 1114.00173   0.629 0.529696    
AG3:D6:B       0.14286    0.34086 1114.00173   0.419 0.675216    
AG3:D7:B       0.31365    0.34556 1114.00173   0.908 0.364253    
AG3:D8:B       0.21841    0.34556 1114.00173   0.632 0.527480    
AG3:D9:B       0.31460    0.34556 1114.00173   0.910 0.362799    
AG3:D10:B      0.01111    0.34556 1114.00173   0.032 0.974355    
---
Signif. codes:  0 ‘***’ 0.001 ‘**’ 0.01 ‘*’ 0.05 ‘.’ 0.1 ‘ ’ 1
```


```
Correlation matrix not shown by default, as p = 40 > 12.
Use print(x, correlation=TRUE)  or
    vcov(x)        if you need it
```


```
anova(m_speed_placebo)
```


```
Type III Analysis of Variance Table with Satterthwaite's method
       Sum Sq Mean Sq NumDF   DenDF  F value    Pr(>F)    
AG       15.9   15.90     1   18.19   3.1289   0.09368 .  
D      6665.5  740.61     9 1114.01 145.7021 < 2.2e-16 ***
B       485.7  485.72     1 1114.00  95.5567 < 2.2e-16 ***
AG:D    336.7   37.41     9 1114.01   7.3607 1.763e-10 ***
AG:B     21.2   21.20     1 1114.00   4.1715   0.04134 *  
D:B     354.1   39.35     9 1114.00   7.7413 4.122e-11 ***
AG:D:B   24.3    2.70     9 1114.00   0.5313   0.85249    
---
Signif. codes:  0 ‘***’ 0.001 ‘**’ 0.01 ‘*’ 0.05 ‘.’ 0.1 ‘ ’ 1
```


```
eta_squared(m_speed_placebo)
```


```
# Effect Size for ANOVA (Type III)

Parameter | Eta2 (partial) |       95% CI
-----------------------------------------
AG        |           0.15 | [0.00, 1.00]
D         |           0.54 | [0.51, 1.00]
B         |           0.08 | [0.06, 1.00]
AG:D      |           0.06 | [0.03, 1.00]
AG:B      |       3.73e-03 | [0.00, 1.00]
D:B       |           0.06 | [0.03, 1.00]
AG:D:B    |       4.27e-03 | [0.00, 1.00]

- One-sided CIs: upper bound fixed at [1.00].
```

###### Note:

There is not enough evidence for a significant difference in speed
between the groups. However, the rate of speed change seems to differ
among the groups (i.e., the interaction between age group and the blocks
of training). We will compare the slopes next. Please note we do not
include the factor “Day” in this comparison, as we are not interested on
a day-by-day comparison. We would only like to know whether the general
difference in slope described in the previous model is derived from
middle-aged adults improving at a generally steeper rate compared to
older adults.


```
slope_comp <- emtrends(m_speed_placebo, pairwise ~ AG, var = "B")$contrasts
```


```
NOTE: Results may be misleading due to involvement in interactions
```


```
summary(slope_comp)
```


```
 contrast  estimate     SE   df t.ratio p.value
 AG2 - AG3    0.158 0.0773 1114   2.042  0.0413

Results are averaged over the levels of: D 
Degrees-of-freedom method: kenward-roger
```

###### Note:

The results suggest that, overall, the slopes of middle-aged adults
are significantly steeper than those in older adults, which is also
apparent in the plots of the behavioral data, especially in the first
two days of training.

###### 2. Effect of stimulation within each age group

###### Middle-aged adults


```
data_subset <- subset(df_speed, D != 0 & AG == 2)
data_subset <- na.omit(data_subset)
data_subset <- droplevels(data_subset)
levels(data_subset$D)
```


```
 [1] "1"  "2"  "3"  "4"  "5"  "6"  "7"  "8"  "9"  "10"
```


```
levels(data_subset$AG)
```


```
[1] "2"
```


```
levels(data_subset$Group)
```


```
[1] "Middle-Placebo" "Middle-Verum"
```


```
m_speed_middle <- lmer(formula = Speed ~ S*D*B + (1 | Code), data=data_subset)
anova(m_speed_middle)
```


```
Type III Analysis of Variance Table with Satterthwaite's method
      Sum Sq Mean Sq NumDF   DenDF  F value Pr(>F)    
S        2.4    2.36     1   18.23   0.4351 0.5177    
D     9097.0 1010.78     9 1118.00 186.4917 <2e-16 ***
B      583.4  583.42     1 1118.00 107.6432 <2e-16 ***
S:D     44.4    4.93     9 1118.00   0.9098 0.5157    
S:B      6.5    6.47     1 1118.00   1.1938 0.2748    
D:B    569.4   63.26     9 1118.00  11.6721 <2e-16 ***
S:D:B   26.6    2.95     9 1118.00   0.5449 0.8422    
---
Signif. codes:  0 ‘***’ 0.001 ‘**’ 0.01 ‘*’ 0.05 ‘.’ 0.1 ‘ ’ 1
```


```
eta_squared(m_speed_middle)
```


```
# Effect Size for ANOVA (Type III)

Parameter | Eta2 (partial) |       95% CI
-----------------------------------------
S         |           0.02 | [0.00, 1.00]
D         |           0.60 | [0.57, 1.00]
B         |           0.09 | [0.06, 1.00]
S:D       |       7.27e-03 | [0.00, 1.00]
S:B       |       1.07e-03 | [0.00, 1.00]
D:B       |           0.09 | [0.06, 1.00]
S:D:B     |       4.37e-03 | [0.00, 1.00]

- One-sided CIs: upper bound fixed at [1.00].
```

###### Older adults


```
data_subset <- subset(df_speed, D != 0 & AG == 3)
data_subset <- na.omit(data_subset)
data_subset <- droplevels(data_subset)
levels(data_subset$D)
```


```
 [1] "1"  "2"  "3"  "4"  "5"  "6"  "7"  "8"  "9"  "10"
```


```
levels(data_subset$AG)
```


```
[1] "3"
```


```
levels(data_subset$Group)
```


```
[1] "Older-Placebo" "Older-Verum"
```


```
m_speed_older <- lmer(formula = Speed ~ S*D*B + (1 | Code), data=data_subset)
anova(m_speed_older)
```


```
Type III Analysis of Variance Table with Satterthwaite's method
       Sum Sq Mean Sq NumDF  DenDF  F value    Pr(>F)    
S       10.06   10.06     1   18.2   3.2016 0.0902194 .  
D     3024.65  336.07     9 1108.0 106.9042 < 2.2e-16 ***
B      342.17  342.17     1 1108.0 108.8444 < 2.2e-16 ***
S:D    100.26   11.14     9 1108.0   3.5438 0.0002388 ***
S:B      1.23    1.23     1 1108.0   0.3924 0.5311796    
D:B    102.17   11.35     9 1108.0   3.6111 0.0001889 ***
S:D:B   32.77    3.64     9 1108.0   1.1582 0.3184684    
---
Signif. codes:  0 ‘***’ 0.001 ‘**’ 0.01 ‘*’ 0.05 ‘.’ 0.1 ‘ ’ 1
```


```
eta_squared(m_speed_older)
```


```
# Effect Size for ANOVA (Type III)

Parameter | Eta2 (partial) |       95% CI
-----------------------------------------
S         |           0.15 | [0.00, 1.00]
D         |           0.46 | [0.43, 1.00]
B         |           0.09 | [0.06, 1.00]
S:D       |           0.03 | [0.01, 1.00]
S:B       |       3.54e-04 | [0.00, 1.00]
D:B       |           0.03 | [0.01, 1.00]
S:D:B     |       9.32e-03 | [0.00, 1.00]

- One-sided CIs: upper bound fixed at [1.00].
```

##### Normalized speed of execution in the finger-tapping task (i.e., speed dynamics)

The following statistical tests concern the normalized speed of
execution in the finger-tapping task, quantified as the total number of
sequences generated in each training block, divided by the speed of the
first training block.


```
speed_norm_file_name <-  "AP3_speed_norm_MS_27_03_2023.txt"
file_path <- file.path(paste(python_data_directory, speed_norm_file_name, sep='/'))
df_speed_norm = read.delim(file_path, header = TRUE, na.strings = "NN")
head(df_speed_norm)
```


```
df_speed_norm$Code <- as.factor(df_speed_norm$Code)
df_speed_norm$AG <- as.factor(df_speed_norm$AG)
df_speed_norm$S <- as.factor(df_speed_norm$S)
df_speed_norm$Group <- as.factor(df_speed_norm$Group)
df_speed_norm$Speed_n <- as.numeric(df_speed_norm$Speed_n)
df_speed_norm$D <- as.factor(df_speed_norm$D)
df_speed_norm$B <- as.numeric(df_speed_norm$B)
levels(df_speed_norm$Code)
```


```
 [1] "217201" "217202" "217203" "217205" "217206" "217207" "217208" "217209" "217210" "217251" "217252" "217253" "217254" "217255" "217257" "217258" "217259" "217260"
[19] "217301" "217302" "217303" "217304" "217305" "217306" "217307" "217308" "217309" "217310" "217311" "217312" "217351" "217352" "217353" "217354" "217355" "217356"
[37] "217357" "217358" "217359" "217360"
```


```
levels(df_speed_norm$AG)
```


```
[1] "2" "3"
```


```
levels(df_speed_norm$S)
```


```
[1] "1" "2"
```


```
levels(df_speed_norm$Group)
```


```
[1] "Middle-Placebo" "Middle-Verum"   "Older-Placebo"  "Older-Verum"
```


```
levels(df_speed_norm$D)
```


```
 [1] "1"  "2"  "3"  "4"  "5"  "6"  "7"  "8"  "9"  "10"
```


###### Choose a model to approximate the data


```
m1 <- lm(formula = Speed_n ~ AG*S*D*B, data=df_speed_norm)
m2 <- lmer(formula = Speed_n ~ AG*S*D*B + (1 | Code), data=df_speed_norm)
BIC(m1, m2)
```


```
m3 <- lmer(formula = Speed_n ~ AG*S*D*B + (1 + B|Code), data=df_speed_norm)
anova(m2, m3)
```


```
refitting model(s) with ML (instead of REML)
```


```
Data: df_speed_norm
Models:
m2: Speed_n ~ AG * S * D * B + (1 | Code)
m3: Speed_n ~ AG * S * D * B + (1 + B | Code)
   npar     AIC     BIC logLik deviance  Chisq Df Pr(>Chisq)    
m2   82 -533.05 -60.828 348.52  -697.05                         
m3   84 -542.88 -59.148 355.44  -710.88 13.838  2  0.0009891 ***
---
Signif. codes:  0 ‘***’ 0.001 ‘**’ 0.01 ‘*’ 0.05 ‘.’ 0.1 ‘ ’ 1
```

###### Note:

The model including a random intercept and a random slope is the best
among the tested models. We will then use m3 to test the speed dynamics
among the groups, following the same rationale we described when testing
the speed.

###### Speed dynamics model


```
m_speed_dynamics <- lmer(formula = Speed_n ~ AG*S*D*B + (1 + B|Code), data=df_speed_norm)
summary(m_speed_dynamics)
```


```
Linear mixed model fit by REML. t-tests use Satterthwaite's method ['lmerModLmerTest']
Formula: Speed_n ~ AG * S * D * B + (1 + B | Code)
   Data: df_speed_norm

REML criterion at convergence: -255.5

Scaled residuals: 
    Min      1Q  Median      3Q     Max 
-5.9599 -0.5751  0.0001  0.5798  4.1681 

Random effects:
 Groups   Name        Variance  Std.Dev. Corr
 Code     (Intercept) 0.1271613 0.35660      
          B           0.0002804 0.01675  0.06
 Residual             0.0403824 0.20095      
Number of obs: 2342, groups:  Code, 40

Fixed effects:
               Estimate Std. Error         df t value Pr(>|t|)    
(Intercept)   9.715e-01  1.273e-01  5.533e+01   7.629 3.39e-10 ***
AG3          -2.709e-02  1.801e-01  5.533e+01  -0.150 0.880987    
S2           -1.246e-02  1.801e-01  5.533e+01  -0.069 0.945084    
D2            3.131e-01  8.611e-02  2.200e+03   3.637 0.000283 ***
D3            8.083e-01  8.366e-02  2.191e+03   9.661  < 2e-16 ***
D4            1.112e+00  8.366e-02  2.191e+03  13.286  < 2e-16 ***
D5            1.359e+00  8.366e-02  2.191e+03  16.247  < 2e-16 ***
D6            1.401e+00  8.611e-02  2.200e+03  16.274  < 2e-16 ***
D7            1.612e+00  8.366e-02  2.191e+03  19.263  < 2e-16 ***
D8            1.739e+00  8.366e-02  2.191e+03  20.785  < 2e-16 ***
D9            1.951e+00  8.366e-02  2.191e+03  23.320  < 2e-16 ***
D10           1.940e+00  8.907e-02  2.208e+03  21.776  < 2e-16 ***
B             1.204e-01  1.609e-02  7.183e+02   7.484 2.11e-13 ***
AG3:S2        1.528e-02  2.547e-01  5.533e+01   0.060 0.952378    
AG3:D2       -1.349e-01  1.201e-01  2.195e+03  -1.124 0.261130    
AG3:D3       -4.254e-01  1.183e-01  2.191e+03  -3.595 0.000331 ***
AG3:D4       -5.946e-01  1.183e-01  2.191e+03  -5.025 5.44e-07 ***
AG3:D5       -7.554e-01  1.183e-01  2.191e+03  -6.384 2.10e-10 ***
AG3:D6       -5.639e-01  1.218e-01  2.200e+03  -4.631 3.85e-06 ***
AG3:D7       -6.303e-01  1.183e-01  2.191e+03  -5.327 1.10e-07 ***
AG3:D8       -6.619e-01  1.183e-01  2.191e+03  -5.594 2.50e-08 ***
AG3:D9       -7.521e-01  1.183e-01  2.191e+03  -6.356 2.51e-10 ***
AG3:D10      -6.212e-01  1.222e-01  2.201e+03  -5.083 4.02e-07 ***
S2:D2        -1.071e-01  1.201e-01  2.195e+03  -0.892 0.372351    
S2:D3        -1.590e-01  1.183e-01  2.191e+03  -1.344 0.179188    
S2:D4        -1.810e-01  1.183e-01  2.191e+03  -1.530 0.126150    
S2:D5        -2.733e-01  1.183e-01  2.191e+03  -2.310 0.020969 *  
S2:D6        -2.876e-02  1.201e-01  2.195e+03  -0.240 0.810723    
S2:D7        -1.680e-01  1.183e-01  2.191e+03  -1.420 0.155761    
S2:D8        -1.639e-01  1.183e-01  2.191e+03  -1.385 0.166044    
S2:D9        -3.212e-01  1.183e-01  2.191e+03  -2.714 0.006692 ** 
S2:D10       -2.155e-01  1.222e-01  2.201e+03  -1.764 0.077940 .  
AG3:B        -6.520e-02  2.275e-02  7.183e+02  -2.866 0.004284 ** 
S2:B         -4.455e-02  2.275e-02  7.183e+02  -1.958 0.050616 .  
D2:B         -4.302e-03  2.210e-02  2.202e+03  -0.195 0.845704    
D3:B         -5.445e-02  2.148e-02  2.191e+03  -2.535 0.011325 *  
D4:B         -8.047e-02  2.148e-02  2.191e+03  -3.746 0.000184 ***
D5:B         -1.060e-01  2.148e-02  2.191e+03  -4.936 8.56e-07 ***
D6:B         -6.671e-02  2.210e-02  2.202e+03  -3.018 0.002575 ** 
D7:B         -1.106e-01  2.148e-02  2.191e+03  -5.150 2.83e-07 ***
D8:B         -1.084e-01  2.148e-02  2.191e+03  -5.045 4.92e-07 ***
D9:B         -1.400e-01  2.148e-02  2.191e+03  -6.518 8.81e-11 ***
D10:B        -1.239e-01  2.286e-02  2.211e+03  -5.421 6.56e-08 ***
AG3:S2:D2     1.199e-01  1.695e-01  2.194e+03   0.707 0.479474    
AG3:S2:D3     3.320e-01  1.675e-01  2.191e+03   1.982 0.047608 *  
AG3:S2:D4     3.461e-01  1.673e-01  2.191e+03   2.069 0.038696 *  
AG3:S2:D5     4.400e-01  1.673e-01  2.191e+03   2.630 0.008605 ** 
AG3:S2:D6     2.404e-01  1.698e-01  2.195e+03   1.416 0.156966    
AG3:S2:D7     2.241e-01  1.686e-01  2.194e+03   1.329 0.183852    
AG3:S2:D8     2.749e-01  1.686e-01  2.194e+03   1.631 0.103083    
AG3:S2:D9     4.055e-01  1.686e-01  2.194e+03   2.406 0.016228 *  
AG3:S2:D10    3.034e-01  1.713e-01  2.199e+03   1.771 0.076725 .  
AG3:S2:B      5.591e-02  3.217e-02  7.183e+02   1.738 0.082707 .  
AG3:D2:B      1.528e-02  3.082e-02  2.196e+03   0.496 0.620105    
AG3:D3:B      4.513e-02  3.038e-02  2.191e+03   1.486 0.137532    
AG3:D4:B      7.401e-02  3.038e-02  2.191e+03   2.436 0.014933 *  
AG3:D5:B      1.081e-01  3.038e-02  2.191e+03   3.557 0.000383 ***
AG3:D6:B      5.977e-02  3.126e-02  2.202e+03   1.912 0.055998 .  
AG3:D7:B      9.798e-02  3.038e-02  2.191e+03   3.225 0.001279 ** 
AG3:D8:B      9.323e-02  3.038e-02  2.191e+03   3.069 0.002177 ** 
AG3:D9:B      1.161e-01  3.038e-02  2.191e+03   3.820 0.000137 ***
AG3:D10:B     9.764e-02  3.137e-02  2.203e+03   3.113 0.001879 ** 
S2:D2:B       3.428e-02  3.082e-02  2.196e+03   1.112 0.266210    
S2:D3:B       3.354e-02  3.038e-02  2.191e+03   1.104 0.269793    
S2:D4:B       4.698e-02  3.038e-02  2.191e+03   1.546 0.122167    
S2:D5:B       7.434e-02  3.038e-02  2.191e+03   2.447 0.014491 *  
S2:D6:B       8.547e-03  3.082e-02  2.196e+03   0.277 0.781577    
S2:D7:B       5.616e-02  3.038e-02  2.191e+03   1.849 0.064641 .  
S2:D8:B       3.623e-02  3.038e-02  2.191e+03   1.192 0.233251    
S2:D9:B       6.768e-02  3.038e-02  2.191e+03   2.228 0.025994 *  
S2:D10:B      5.516e-02  3.137e-02  2.203e+03   1.758 0.078836 .  
AG3:S2:D2:B  -3.370e-02  4.343e-02  2.194e+03  -0.776 0.437781    
AG3:S2:D3:B  -5.205e-02  4.312e-02  2.192e+03  -1.207 0.227492    
AG3:S2:D4:B  -6.323e-02  4.297e-02  2.191e+03  -1.472 0.141254    
AG3:S2:D5:B  -9.642e-02  4.297e-02  2.191e+03  -2.244 0.024931 *  
AG3:S2:D6:B  -5.513e-02  4.359e-02  2.196e+03  -1.265 0.206110    
AG3:S2:D7:B  -7.888e-02  4.328e-02  2.195e+03  -1.822 0.068525 .  
AG3:S2:D8:B  -8.049e-02  4.328e-02  2.195e+03  -1.860 0.063069 .  
AG3:S2:D9:B  -9.527e-02  4.328e-02  2.195e+03  -2.201 0.027836 *  
AG3:S2:D10:B -9.440e-02  4.398e-02  2.201e+03  -2.146 0.031965 *  
---
Signif. codes:  0 ‘***’ 0.001 ‘**’ 0.01 ‘*’ 0.05 ‘.’ 0.1 ‘ ’ 1
```


```
Correlation matrix not shown by default, as p = 80 > 12.
Use print(x, correlation=TRUE)  or
    vcov(x)        if you need it
```


```
anova(m_speed_dynamics)
```


```
Type III Analysis of Variance Table with Satterthwaite's method
          Sum Sq Mean Sq NumDF   DenDF  F value    Pr(>F)    
AG         0.492  0.4918     1   36.02  12.1786  0.001295 ** 
S          0.003  0.0032     1   36.02   0.0797  0.779314    
D        118.513 13.1681     9 2194.97 326.0853 < 2.2e-16 ***
B          4.927  4.9268     1   36.44 122.0046 3.488e-13 ***
AG:S       0.062  0.0622     1   36.02   1.5410  0.222492    
AG:D       4.042  0.4491     9 2194.97  11.1220 < 2.2e-16 ***
S:D        0.308  0.0342     9 2194.97   0.8474  0.572299    
AG:B       0.001  0.0008     1   36.44   0.0191  0.890901    
S:B        0.047  0.0471     1   36.44   1.1669  0.287141    
D:B        4.881  0.5423     9 2195.84  13.4299 < 2.2e-16 ***
AG:S:D     0.446  0.0496     9 2194.97   1.2271  0.273423    
AG:S:B     0.016  0.0159     1   36.44   0.3948  0.533728    
AG:D:B     0.960  0.1067     9 2195.84   2.6418  0.004827 ** 
S:D:B      0.275  0.0305     9 2195.84   0.7556  0.657829    
AG:S:D:B   0.372  0.0413     9 2195.84   1.0231  0.418705    
---
Signif. codes:  0 ‘***’ 0.001 ‘**’ 0.01 ‘*’ 0.05 ‘.’ 0.1 ‘ ’ 1
```


```
eta_squared(m_speed_dynamics)
```


```
# Effect Size for ANOVA (Type III)

Parameter | Eta2 (partial) |       95% CI
-----------------------------------------
AG        |           0.25 | [0.07, 1.00]
S         |       2.21e-03 | [0.00, 1.00]
D         |           0.57 | [0.55, 1.00]
B         |           0.77 | [0.65, 1.00]
AG:S      |           0.04 | [0.00, 1.00]
AG:D      |           0.04 | [0.03, 1.00]
S:D       |       3.46e-03 | [0.00, 1.00]
AG:B      |       5.23e-04 | [0.00, 1.00]
S:B       |           0.03 | [0.00, 1.00]
D:B       |           0.05 | [0.03, 1.00]
AG:S:D    |       5.01e-03 | [0.00, 1.00]
AG:S:B    |           0.01 | [0.00, 1.00]
AG:D:B    |           0.01 | [0.00, 1.00]
S:D:B     |       3.09e-03 | [0.00, 1.00]
AG:S:D:B  |       4.18e-03 | [0.00, 1.00]

- One-sided CIs: upper bound fixed at [1.00].
```

###### Note:

There is a significant difference among the age groups, but no
significant effect of stimulation. We will compare the placebo groups
next.

###### Age-related differences in speed dynamics between unstimulated groups


```
data_subset <- subset(df_speed_norm, S == 2)
data_subset <- na.omit(data_subset)
data_subset <- droplevels(data_subset)
levels(data_subset$D)
```


```
 [1] "1"  "2"  "3"  "4"  "5"  "6"  "7"  "8"  "9"  "10"
```


```
levels(data_subset$AG)
```


```
[1] "2" "3"
```


```
levels(data_subset$S)
```


```
[1] "2"
```


```
levels(data_subset$Group)
```


```
[1] "Middle-Placebo" "Older-Placebo"
```


```
m_speed_dynamics_placebo <- lmer(formula = Speed_n ~ AG*D*B + (1 + B|Code), data=data_subset)
```


```
Warning: Model failed to converge with max|grad| = 0.00706887 (tol = 0.002, component 1)
```


```
anova(m_speed_dynamics_placebo)
```


```
Type III Analysis of Variance Table with Satterthwaite's method
       Sum Sq Mean Sq NumDF   DenDF  F value    Pr(>F)    
AG      0.123  0.1230     1   18.05   3.1354 0.0934994 .  
D      57.460  6.3845     9 1098.21 162.7028 < 2.2e-16 ***
B       1.946  1.9456     1   18.31  49.5831 1.302e-06 ***
AG:D    1.170  0.1300     9 1098.21   3.3125 0.0005317 ***
AG:B    0.005  0.0046     1   18.31   0.1175 0.7356816    
D:B     2.636  0.2929     9 1098.49   7.4631 1.206e-10 ***
AG:D:B  0.111  0.0123     9 1098.49   0.3137 0.9706880    
---
Signif. codes:  0 ‘***’ 0.001 ‘**’ 0.01 ‘*’ 0.05 ‘.’ 0.1 ‘ ’ 1
```


```
eta_squared(m_speed_dynamics_placebo)
```


```
# Effect Size for ANOVA (Type III)

Parameter | Eta2 (partial) |       95% CI
-----------------------------------------
AG        |           0.15 | [0.00, 1.00]
D         |           0.57 | [0.54, 1.00]
B         |           0.73 | [0.52, 1.00]
AG:D      |           0.03 | [0.01, 1.00]
AG:B      |       6.38e-03 | [0.00, 1.00]
D:B       |           0.06 | [0.03, 1.00]
AG:D:B    |       2.56e-03 | [0.00, 1.00]

- One-sided CIs: upper bound fixed at [1.00].
```

###### Note:

There is only a trend towards a significant difference among the age
groups, and a significant interaction suggesting different magnitudes
between age groups across days. However, there is no evidence for a
difference in the rate of improvement, which is the parameter of
interest when discussing dynamics.

###### Stimulation-related differences in speed dynamics in middle-aged adults


```
data_subset <- subset(df_speed_norm, AG == 2)
data_subset <- na.omit(data_subset)
data_subset <- droplevels(data_subset)
levels(data_subset$D)
```


```
 [1] "1"  "2"  "3"  "4"  "5"  "6"  "7"  "8"  "9"  "10"
```


```
levels(data_subset$AG)
```


```
[1] "2"
```


```
levels(data_subset$S)
```


```
[1] "1" "2"
```


```
levels(data_subset$Group)
```


```
[1] "Middle-Placebo" "Middle-Verum"
```


```
m_speed_dynamics_middle <- lmer(formula = Speed_n ~ S*D*B + (1 + B|Code), data=data_subset)
```


```
Warning: Model failed to converge with max|grad| = 0.0241171 (tol = 0.002, component 1)
```


```
anova(m_speed_dynamics_middle)
```


```
Type III Analysis of Variance Table with Satterthwaite's method
      Sum Sq Mean Sq NumDF   DenDF  F value    Pr(>F)    
S      0.048  0.0478     1   18.17   1.0755    0.3133    
D     81.709  9.0788     9 1101.78 204.2348 < 2.2e-16 ***
B      2.182  2.1825     1   18.10  49.0970 1.486e-06 ***
S:D    0.500  0.0556     9 1101.78   1.2502    0.2603    
S:B    0.004  0.0038     1   18.10   0.0847    0.7743    
D:B    4.870  0.5411     9 1102.11  12.1717 < 2.2e-16 ***
S:D:B  0.444  0.0493     9 1102.11   1.1098    0.3526    
---
Signif. codes:  0 ‘***’ 0.001 ‘**’ 0.01 ‘*’ 0.05 ‘.’ 0.1 ‘ ’ 1
```


```
eta_squared(m_speed_dynamics_middle)
```


```
# Effect Size for ANOVA (Type III)

Parameter | Eta2 (partial) |       95% CI
-----------------------------------------
S         |           0.06 | [0.00, 1.00]
D         |           0.63 | [0.60, 1.00]
B         |           0.73 | [0.52, 1.00]
S:D       |           0.01 | [0.00, 1.00]
S:B       |       4.66e-03 | [0.00, 1.00]
D:B       |           0.09 | [0.06, 1.00]
S:D:B     |       8.98e-03 | [0.00, 1.00]

- One-sided CIs: upper bound fixed at [1.00].
```

###### Note:

The model failed to converge, so we will use a model including only
random intercepts for this test.


```
m_speed_dynamics_middle <- lmer(formula = Speed_n ~ S*D*B + (1|Code), data=data_subset)
anova(m_speed_dynamics_middle)
```


```
Type III Analysis of Variance Table with Satterthwaite's method
      Sum Sq Mean Sq NumDF   DenDF  F value Pr(>F)    
S      0.047  0.0472     1   18.84   1.0391 0.3209    
D     81.808  9.0898     9 1118.01 199.9231 <2e-16 ***
B      5.297  5.2972     1 1118.00 116.5065 <2e-16 ***
S:D    0.491  0.0546     9 1118.01   1.2002 0.2908    
S:B    0.010  0.0099     1 1118.00   0.2188 0.6400    
D:B    4.878  0.5420     9 1118.00  11.9204 <2e-16 ***
S:D:B  0.433  0.0481     9 1118.00   1.0587 0.3909    
---
Signif. codes:  0 ‘***’ 0.001 ‘**’ 0.01 ‘*’ 0.05 ‘.’ 0.1 ‘ ’ 1
```


```
eta_squared(m_speed_dynamics_middle)
```


```
# Effect Size for ANOVA (Type III)

Parameter | Eta2 (partial) |       95% CI
-----------------------------------------
S         |           0.05 | [0.00, 1.00]
D         |           0.62 | [0.59, 1.00]
B         |           0.09 | [0.07, 1.00]
S:D       |       9.57e-03 | [0.00, 1.00]
S:B       |       1.96e-04 | [0.00, 1.00]
D:B       |           0.09 | [0.06, 1.00]
S:D:B     |       8.45e-03 | [0.00, 1.00]

- One-sided CIs: upper bound fixed at [1.00].
```

###### Note:

The data, as described by the model, does not provide evidence for a
difference in speed dynamics related to stimulation in middle-aged
adults.

###### Stimulation-related differences in speed dynamics in older adults


```
data_subset <- subset(df_speed_norm, AG == 3)
data_subset <- na.omit(data_subset)
data_subset <- droplevels(data_subset)
levels(data_subset$D)
```


```
 [1] "1"  "2"  "3"  "4"  "5"  "6"  "7"  "8"  "9"  "10"
```


```
levels(data_subset$AG)
```


```
[1] "3"
```


```
levels(data_subset$S)
```


```
[1] "1" "2"
```


```
levels(data_subset$Group)
```


```
[1] "Older-Placebo" "Older-Verum"
```


```
m_speed_dynamics_older <- lmer(formula = Speed_n ~ S*D*B + (1 + B|Code), data=data_subset)
anova(m_speed_dynamics_older)
```


```
Type III Analysis of Variance Table with Satterthwaite's method
      Sum Sq Mean Sq NumDF   DenDF  F value    Pr(>F)    
S      0.018  0.0182     1   18.02   0.5023  0.487583    
D     41.117  4.5686     9 1092.95 125.9285 < 2.2e-16 ***
B      2.892  2.8915     1   18.40  79.7022 4.142e-08 ***
S:D    0.259  0.0287     9 1092.95   0.7923  0.623517    
S:B    0.067  0.0670     1   18.40   1.8462  0.190647    
D:B    0.984  0.1093     9 1093.40   3.0124  0.001472 ** 
S:D:B  0.205  0.0228     9 1093.40   0.6279  0.773966    
---
Signif. codes:  0 ‘***’ 0.001 ‘**’ 0.01 ‘*’ 0.05 ‘.’ 0.1 ‘ ’ 1
```


```
eta_squared(m_speed_dynamics_older)
```


```
# Effect Size for ANOVA (Type III)

Parameter | Eta2 (partial) |       95% CI
-----------------------------------------
S         |           0.03 | [0.00, 1.00]
D         |           0.51 | [0.48, 1.00]
B         |           0.81 | [0.66, 1.00]
S:D       |       6.48e-03 | [0.00, 1.00]
S:B       |           0.09 | [0.00, 1.00]
D:B       |           0.02 | [0.01, 1.00]
S:D:B     |       5.14e-03 | [0.00, 1.00]

- One-sided CIs: upper bound fixed at [1.00].
```

###### Note:

The data, as described by the model, does not provide evidence for a
difference in speed dynamics related to stimulation in older adults.

##### Accuracy of execution in the finger-tapping task

The following statistical tests concern the speed of execution in the
finger-tapping task, quantified as the total number of sequences
generated in each training block.


```
acc_file_name <-  "AP3_acc_MS_27_03_2023.txt"
file_path <- file.path(paste(python_data_directory, acc_file_name, sep='/'))
df_acc = read.delim(file_path, header = TRUE, na.strings = "NN")
head(df_acc)
```


```
df_acc$Code <- as.factor(df_acc$Code)
df_acc$AG <- as.factor(df_acc$AG)
df_acc$S <- as.factor(df_acc$S)
df_acc$Group <- as.factor(df_acc$Group)
df_acc$Acc <- as.numeric(df_acc$Acc)
df_acc$D <- as.factor(df_acc$D)
df_acc$B <- as.numeric(df_acc$B)
```


```
Warning: NAs introduced by coercion
```


```
levels(df_acc$Code)
```


```
 [1] "217201" "217202" "217203" "217205" "217206" "217207" "217208" "217209" "217210" "217251" "217252" "217253" "217254" "217255" "217257" "217258" "217259" "217260"
[19] "217301" "217302" "217303" "217304" "217305" "217306" "217307" "217308" "217309" "217310" "217311" "217312" "217351" "217352" "217353" "217354" "217355" "217356"
[37] "217357" "217358" "217359" "217360"
```


```
levels(df_acc$AG)
```


```
[1] "2" "3"
```


```
levels(df_acc$S)
```


```
[1] "1" "2"
```


```
levels(df_acc$Group)
```


```
[1] "Middle-Placebo" "Middle-Verum"   "Older-Placebo"  "Older-Verum"
```


```
levels(df_acc$D)
```


```
 [1] "0"  "1"  "2"  "3"  "4"  "5"  "6"  "7"  "8"  "9"  "10"
```


###### Compare initial training accuracy


```
data_subset <- subset(df_acc, D == 1 & B == 1)
data_subset <- droplevels(data_subset)
levels(data_subset$D)
```


```
[1] "1"
```


```
levels(data_subset$Group)
```


```
[1] "Middle-Placebo" "Middle-Verum"   "Older-Placebo"  "Older-Verum"
```


```
m_acc_b1 <- lm(formula = Acc ~ AG*S, data=data_subset)
summary(m_acc_b1)
```


```
Call:
lm(formula = Acc ~ AG * S, data = data_subset)

Residuals:
     Min       1Q   Median       3Q      Max 
-0.34810 -0.09213  0.00905  0.10413  0.50905 

Coefficients:
            Estimate Std. Error t value Pr(>|t|)    
(Intercept)  0.78580    0.05144  15.275  < 2e-16 ***
AG3         -0.29485    0.07275  -4.053 0.000258 ***
S2           0.10633    0.07275   1.462 0.152540    
AG3:S2       0.18437    0.10289   1.792 0.081549 .  
---
Signif. codes:  0 ‘***’ 0.001 ‘**’ 0.01 ‘*’ 0.05 ‘.’ 0.1 ‘ ’ 1

Residual standard error: 0.1627 on 36 degrees of freedom
Multiple R-squared:  0.4829,    Adjusted R-squared:  0.4398 
F-statistic: 11.21 on 3 and 36 DF,  p-value: 2.436e-05
```


```
anova(m_acc_b1)
```


```
Analysis of Variance Table

Response: Acc
          Df  Sum Sq Mean Sq F value    Pr(>F)    
AG         1 0.41073 0.41073 15.5203 0.0003595 ***
S          1 0.39407 0.39407 14.8911 0.0004541 ***
AG:S       1 0.08498 0.08498  3.2111 0.0815488 .  
Residuals 36 0.95270 0.02646                      
---
Signif. codes:  0 ‘***’ 0.001 ‘**’ 0.01 ‘*’ 0.05 ‘.’ 0.1 ‘ ’ 1
```


```
eta_squared(m_acc_b1)
```


```
# Effect Size for ANOVA (Type I)

Parameter | Eta2 (partial) |       95% CI
-----------------------------------------
AG        |           0.30 | [0.11, 1.00]
S         |           0.29 | [0.10, 1.00]
AG:S      |           0.08 | [0.00, 1.00]

- One-sided CIs: upper bound fixed at [1.00].
```

###### Differences in initial accuracy between the groups receiving placebo stimulation


```
data_subset <- subset(df_acc, D == 1 & B == 1 & S == 2)
data_subset <- droplevels(data_subset)
levels(data_subset$D)
```


```
[1] "1"
```


```
levels(data_subset$S)
```


```
[1] "2"
```


```
levels(data_subset$Group)
```


```
[1] "Middle-Placebo" "Older-Placebo"
```


```
m_acc_b1_placebo <- lm(formula = Acc ~ AG, data=data_subset)
summary(m_acc_b1_placebo)
```


```
Call:
lm(formula = Acc ~ AG, data = data_subset)

Residuals:
      Min        1Q    Median        3Q       Max 
-0.236194 -0.080134  0.007871  0.083431  0.218351 

Coefficients:
            Estimate Std. Error t value Pr(>|t|)    
(Intercept)  0.89213    0.03484  25.609  1.3e-15 ***
AG3         -0.11048    0.04927  -2.243   0.0378 *  
---
Signif. codes:  0 ‘***’ 0.001 ‘**’ 0.01 ‘*’ 0.05 ‘.’ 0.1 ‘ ’ 1

Residual standard error: 0.1102 on 18 degrees of freedom
Multiple R-squared:  0.2184,    Adjusted R-squared:  0.175 
F-statistic: 5.029 on 1 and 18 DF,  p-value: 0.03776
```


```
anova(m_acc_b1_placebo)
```


```
Analysis of Variance Table

Response: Acc
          Df  Sum Sq  Mean Sq F value  Pr(>F)  
AG         1 0.06103 0.061030   5.029 0.03776 *
Residuals 18 0.21844 0.012136                  
---
Signif. codes:  0 ‘***’ 0.001 ‘**’ 0.01 ‘*’ 0.05 ‘.’ 0.1 ‘ ’ 1
```


```
eta_squared(m_acc_b1_placebo)
```


```
For one-way between subjects designs, partial eta squared is equivalent to eta squared. Returning eta squared.
```


```
# Effect Size for ANOVA

Parameter | Eta2 |       95% CI
-------------------------------
AG        | 0.22 | [0.01, 1.00]

- One-sided CIs: upper bound fixed at [1.00].
```


```
t.test(Acc ~ AG, data=data_subset)
```


```
    Welch Two Sample t-test

data:  Acc by AG
t = 2.2425, df = 13.934, p-value = 0.04172
alternative hypothesis: true difference in means between group 2 and group 3 is not equal to 0
95 percent confidence interval:
 0.004769313 0.216191671
sample estimates:
mean in group 2 mean in group 3 
      0.8921291       0.7816486
```


```
cohens_d(Acc ~ AG, data=data_subset)
```


```
Cohen's d |       95% CI
------------------------
1.00      | [0.06, 1.93]

- Estimated using pooled SD.
```

###### Note:

In this case, the group of middle-aged adults receiving placebo
stimulation started off at a significantly higher accuracy level
compared to older adults. In addition to this test, we will compare the
middle-aged adults receiving verum stimulation to the older adults
receiving the placebo. The reason is that in the main manuscript we
discuss that the older-placebo group was initially more accurate than
other groups of older adults we have recruited, and that their initial
accuracy was more similar to the one seen in middle-aged adults.


```
data_subset <- subset(df_acc, D == 1 & B == 1 & (Group == 'Middle-Verum' | Group == 'Older-Placebo'))
data_subset <- droplevels(data_subset)
levels(data_subset$D)
```


```
[1] "1"
```


```
levels(data_subset$S)
```


```
[1] "1" "2"
```


```
levels(data_subset$Group)
```


```
[1] "Middle-Verum"  "Older-Placebo"
```


```
m_acc_b1_middle_older <- lm(formula = Acc ~ S, data=data_subset)
summary(m_acc_b1_middle_older)
```


```
Call:
lm(formula = Acc ~ S, data = data_subset)

Residuals:
     Min       1Q   Median       3Q      Max 
-0.28580 -0.10040  0.04753  0.10413  0.21835 

Coefficients:
             Estimate Std. Error t value Pr(>|t|)    
(Intercept)  0.785800   0.048816   16.10 3.93e-12 ***
S2          -0.004151   0.069036   -0.06    0.953    
---
Signif. codes:  0 ‘***’ 0.001 ‘**’ 0.01 ‘*’ 0.05 ‘.’ 0.1 ‘ ’ 1

Residual standard error: 0.1544 on 18 degrees of freedom
Multiple R-squared:  0.0002008, Adjusted R-squared:  -0.05534 
F-statistic: 0.003616 on 1 and 18 DF,  p-value: 0.9527
```


```
anova(m_acc_b1_middle_older)
```


```
Analysis of Variance Table

Response: Acc
          Df  Sum Sq   Mean Sq F value Pr(>F)
S          1 0.00009 0.0000862  0.0036 0.9527
Residuals 18 0.42893 0.0238297
```


```
eta_squared(m_acc_b1_middle_older)
```


```
For one-way between subjects designs, partial eta squared is equivalent to eta squared. Returning eta squared.
```


```
# Effect Size for ANOVA

Parameter |     Eta2 |       95% CI
-----------------------------------
S         | 2.01e-04 | [0.00, 1.00]

- One-sided CIs: upper bound fixed at [1.00].
```

###### Note:

As suspected from visual inspection, the initial accuracy is quite
similar among the groups. Frequentist statistics cannot typically be
used to prove equality, however, we can claim that if, in truth, the
average of both groups was the same, the likelihood of finding at least
the observed difference would be above 95%, which is highly likely.

###### Comparison of initial accuracy in middle-aged groups


```
data_subset <- subset(df_acc, D == 1 & B == 1 & AG == 2)
data_subset <- droplevels(data_subset)
levels(data_subset$D)
```


```
[1] "1"
```


```
levels(data_subset$AG)
```


```
[1] "2"
```


```
levels(data_subset$S)
```


```
[1] "1" "2"
```


```
levels(data_subset$Group)
```


```
[1] "Middle-Placebo" "Middle-Verum"
```


```
m_acc_b1_middle <- lm(formula = Acc ~ S, data=data_subset)
summary(m_acc_b1_middle)
```


```
Call:
lm(formula = Acc ~ S, data = data_subset)

Residuals:
     Min       1Q   Median       3Q      Max 
-0.28580 -0.08275  0.03370  0.08642  0.21420 

Coefficients:
            Estimate Std. Error t value Pr(>|t|)    
(Intercept)  0.78580    0.04156  18.906 2.54e-13 ***
S2           0.10633    0.05878   1.809   0.0872 .  
---
Signif. codes:  0 ‘***’ 0.001 ‘**’ 0.01 ‘*’ 0.05 ‘.’ 0.1 ‘ ’ 1

Residual standard error: 0.1314 on 18 degrees of freedom
Multiple R-squared:  0.1538,    Adjusted R-squared:  0.1068 
F-statistic: 3.272 on 1 and 18 DF,  p-value: 0.08719
```


```
anova(m_acc_b1_middle)
```


```
Analysis of Variance Table

Response: Acc
          Df  Sum Sq  Mean Sq F value  Pr(>F)  
S          1 0.05653 0.056530  3.2724 0.08719 .
Residuals 18 0.31094 0.017274                  
---
Signif. codes:  0 ‘***’ 0.001 ‘**’ 0.01 ‘*’ 0.05 ‘.’ 0.1 ‘ ’ 1
```


```
eta_squared(m_acc_b1_middle)
```


```
For one-way between subjects designs, partial eta squared is equivalent to eta squared. Returning eta squared.
```


```
# Effect Size for ANOVA

Parameter | Eta2 |       95% CI
-------------------------------
S         | 0.15 | [0.00, 1.00]

- One-sided CIs: upper bound fixed at [1.00].
```


```
t.test(Acc ~ S, data=data_subset)
```


```
    Welch Two Sample t-test

data:  Acc by S
t = -1.809, df = 12.343, p-value = 0.09486
alternative hypothesis: true difference in means between group 1 and group 2 is not equal to 0
95 percent confidence interval:
 -0.23400237  0.02134364
sample estimates:
mean in group 1 mean in group 2 
      0.7857998       0.8921291
```


```
cohens_d(Acc ~ S, data=data_subset)
```


```
Cohen's d |        95% CI
-------------------------
-0.81     | [-1.71, 0.12]

- Estimated using pooled SD.
```

###### Comparison of initial accuracy in older groups


```
data_subset <- subset(df_acc, D == 1 & B == 1 & AG == 3)
data_subset <- droplevels(data_subset)
levels(data_subset$D)
```


```
[1] "1"
```


```
levels(data_subset$AG)
```


```
[1] "3"
```


```
levels(data_subset$S)
```


```
[1] "1" "2"
```


```
levels(data_subset$Group)
```


```
[1] "Older-Placebo" "Older-Verum"
```


```
m_acc_b1_older <- lm(formula = Acc ~ S, data=data_subset)
summary(m_acc_b1_older)
```


```
Call:
lm(formula = Acc ~ S, data = data_subset)

Residuals:
     Min       1Q   Median       3Q      Max 
-0.34810 -0.12613  0.00905  0.10769  0.50905 

Coefficients:
            Estimate Std. Error t value Pr(>|t|)    
(Intercept)  0.49095    0.05971   8.222 1.66e-07 ***
S2           0.29070    0.08444   3.443   0.0029 ** 
---
Signif. codes:  0 ‘***’ 0.001 ‘**’ 0.01 ‘*’ 0.05 ‘.’ 0.1 ‘ ’ 1

Residual standard error: 0.1888 on 18 degrees of freedom
Multiple R-squared:  0.397, Adjusted R-squared:  0.3635 
F-statistic: 11.85 on 1 and 18 DF,  p-value: 0.002905
```


```
anova(m_acc_b1_older)
```


```
Analysis of Variance Table

Response: Acc
          Df  Sum Sq Mean Sq F value   Pr(>F)   
S          1 0.42252 0.42252  11.851 0.002905 **
Residuals 18 0.64176 0.03565                    
---
Signif. codes:  0 ‘***’ 0.001 ‘**’ 0.01 ‘*’ 0.05 ‘.’ 0.1 ‘ ’ 1
```


```
eta_squared(m_acc_b1_older)
```


```
For one-way between subjects designs, partial eta squared is equivalent to eta squared. Returning eta squared.
```


```
# Effect Size for ANOVA

Parameter | Eta2 |       95% CI
-------------------------------
S         | 0.40 | [0.11, 1.00]

- One-sided CIs: upper bound fixed at [1.00].
```


```
t.test(Acc ~ S, data=data_subset)
```


```
    Welch Two Sample t-test

data:  Acc by S
t = -3.4425, df = 14.678, p-value = 0.003729
alternative hypothesis: true difference in means between group 1 and group 2 is not equal to 0
95 percent confidence interval:
 -0.4710269 -0.1103656
sample estimates:
mean in group 1 mean in group 2 
      0.4909524       0.7816486
```


```
cohens_d(Acc ~ S, data=data_subset)
```


```
Cohen's d |         95% CI
--------------------------
-1.54     | [-2.53, -0.51]

- Estimated using pooled SD.
```

###### Accuracy over the course of training


```
data_subset <- subset(df_acc, D != 0)
data_subset <- droplevels(data_subset)
levels(data_subset$D)
```


```
 [1] "1"  "2"  "3"  "4"  "5"  "6"  "7"  "8"  "9"  "10"
```


```
levels(data_subset$AG)
```


```
[1] "2" "3"
```


```
levels(data_subset$S)
```


```
[1] "1" "2"
```


```
levels(data_subset$Group)
```


```
[1] "Middle-Placebo" "Middle-Verum"   "Older-Placebo"  "Older-Verum"
```


```
m1 <- lm(formula = Acc ~ AG*S*D*B, data=data_subset)
m2 <- lmer(formula = Acc ~ AG*S*D*B + (1|Code), data=data_subset)
BIC(m1, m2)
```

###### Note:

In this case, the linear model is better at explaining the data’s
variance than the mixed-effects model including random intercepts. Next,
we will compare m1 to a model includig also random slopes.


```
m3 <- lmer(formula = Acc ~ AG*S*D*B + (1 + B|Code), data=data_subset, REML=FALSE)
BIC(m1, m3)
```

###### Note:

The mixed-effects model including both random intercepts and slopes
is better, so that’s the one we will use for the tests. We set REML to
“False” because the model did not converge otherwise.


```
m_acc <- lmer(formula = Acc ~ AG*S*D*B + (1 + B|Code), data=data_subset, REML=FALSE)
summary(m_acc)
```


```
Linear mixed model fit by maximum likelihood . t-tests use Satterthwaite's method ['lmerModLmerTest']
Formula: Acc ~ AG * S * D * B + (1 + B | Code)
   Data: data_subset

     AIC      BIC   logLik deviance df.resid 
 -3787.9  -3304.2   1978.0  -3955.9     2257 

Scaled residuals: 
    Min      1Q  Median      3Q     Max 
-6.1749 -0.4879  0.0982  0.5744  5.2688 

Random effects:
 Groups   Name        Variance  Std.Dev. Corr
 Code     (Intercept) 2.358e-03 0.048557     
          B           3.304e-05 0.005748 0.06
 Residual             1.022e-02 0.101115     
Number of obs: 2341, groups:  Code, 40

Fixed effects:
               Estimate Std. Error         df t value Pr(>|t|)    
(Intercept)   7.643e-01  3.349e-02  4.267e+02  22.818  < 2e-16 ***
AG3          -2.262e-01  4.737e-02  4.267e+02  -4.775 2.47e-06 ***
S2            1.375e-01  4.737e-02  4.267e+02   2.904 0.003881 ** 
D2           -1.048e-02  4.330e-02  2.276e+03  -0.242 0.808692    
D3            1.363e-01  4.210e-02  2.262e+03   3.239 0.001219 ** 
D4            1.009e-01  4.210e-02  2.262e+03   2.397 0.016629 *  
D5            8.204e-02  4.210e-02  2.262e+03   1.949 0.051440 .  
D6            9.443e-02  4.363e-02  2.276e+03   2.164 0.030551 *  
D7            1.480e-01  4.210e-02  2.262e+03   3.516 0.000446 ***
D8            9.447e-02  4.210e-02  2.262e+03   2.244 0.024930 *  
D9            1.507e-01  4.210e-02  2.262e+03   3.581 0.000350 ***
D10           1.403e-01  4.476e-02  2.288e+03   3.135 0.001740 ** 
B             1.456e-02  7.857e-03  1.136e+03   1.853 0.064197 .  
AG3:S2        9.416e-02  6.699e-02  4.267e+02   1.406 0.160585    
AG3:D2        1.491e-01  6.039e-02  2.269e+03   2.469 0.013627 *  
AG3:D3       -7.869e-02  5.953e-02  2.262e+03  -1.322 0.186366    
AG3:D4        6.694e-02  5.953e-02  2.262e+03   1.124 0.260980    
AG3:D5        1.196e-01  5.953e-02  2.262e+03   2.009 0.044606 *  
AG3:D6        6.382e-02  6.147e-02  2.276e+03   1.038 0.299265    
AG3:D7        1.021e-01  5.953e-02  2.262e+03   1.714 0.086609 .  
AG3:D8        1.745e-01  5.953e-02  2.262e+03   2.932 0.003403 ** 
AG3:D9        4.516e-02  5.953e-02  2.262e+03   0.759 0.448211    
AG3:D10       1.593e-01  6.144e-02  2.277e+03   2.592 0.009602 ** 
S2:D2        -2.349e-02  6.039e-02  2.269e+03  -0.389 0.697287    
S2:D3        -1.517e-01  5.953e-02  2.262e+03  -2.547 0.010921 *  
S2:D4        -9.897e-02  5.953e-02  2.262e+03  -1.662 0.096564 .  
S2:D5        -7.134e-02  5.953e-02  2.262e+03  -1.198 0.230921    
S2:D6        -8.816e-02  6.063e-02  2.269e+03  -1.454 0.146076    
S2:D7        -1.461e-01  5.953e-02  2.262e+03  -2.455 0.014171 *  
S2:D8        -8.506e-02  5.953e-02  2.262e+03  -1.429 0.153239    
S2:D9        -1.318e-01  5.953e-02  2.262e+03  -2.214 0.026930 *  
S2:D10       -1.132e-01  6.144e-02  2.277e+03  -1.843 0.065521 .  
AG3:B         1.823e-02  1.111e-02  1.136e+03   1.640 0.101196    
S2:B         -1.751e-02  1.111e-02  1.136e+03  -1.576 0.115379    
D2:B          1.700e-02  1.112e-02  2.277e+03   1.529 0.126292    
D3:B         -1.329e-02  1.081e-02  2.262e+03  -1.230 0.218974    
D4:B         -6.295e-03  1.081e-02  2.262e+03  -0.582 0.560376    
D5:B         -2.547e-03  1.081e-02  2.262e+03  -0.236 0.813776    
D6:B         -5.712e-03  1.116e-02  2.277e+03  -0.512 0.608776    
D7:B         -1.582e-02  1.081e-02  2.262e+03  -1.464 0.143436    
D8:B         -6.532e-03  1.081e-02  2.262e+03  -0.604 0.545694    
D9:B         -1.692e-02  1.081e-02  2.262e+03  -1.565 0.117712    
D10:B        -1.311e-02  1.149e-02  2.290e+03  -1.141 0.253942    
AG3:S2:D2    -1.327e-01  8.525e-02  2.267e+03  -1.556 0.119827    
AG3:S2:D3     1.621e-01  8.429e-02  2.262e+03   1.923 0.054642 .  
AG3:S2:D4    -2.813e-02  8.420e-02  2.262e+03  -0.334 0.738317    
AG3:S2:D5    -6.703e-02  8.420e-02  2.262e+03  -0.796 0.426081    
AG3:S2:D6     2.359e-02  8.557e-02  2.269e+03   0.276 0.782855    
AG3:S2:D7    -1.368e-02  8.481e-02  2.266e+03  -0.161 0.871890    
AG3:S2:D8    -5.852e-02  8.481e-02  2.266e+03  -0.690 0.490270    
AG3:S2:D9     5.041e-02  8.481e-02  2.266e+03   0.594 0.552323    
AG3:S2:D10   -1.002e-01  8.616e-02  2.274e+03  -1.163 0.244899    
AG3:S2:B      1.478e-03  1.571e-02  1.136e+03   0.094 0.925095    
AG3:D2:B     -2.730e-02  1.550e-02  2.270e+03  -1.761 0.078443 .  
AG3:D3:B      2.060e-02  1.529e-02  2.262e+03   1.348 0.177859    
AG3:D4:B     -1.090e-02  1.529e-02  2.262e+03  -0.713 0.475718    
AG3:D5:B     -2.058e-02  1.529e-02  2.262e+03  -1.346 0.178335    
AG3:D6:B     -9.219e-03  1.575e-02  2.277e+03  -0.585 0.558387    
AG3:D7:B     -1.892e-02  1.529e-02  2.262e+03  -1.237 0.216079    
AG3:D8:B     -2.032e-02  1.529e-02  2.262e+03  -1.329 0.183973    
AG3:D9:B      6.690e-03  1.529e-02  2.262e+03   0.438 0.661719    
AG3:D10:B    -1.957e-02  1.577e-02  2.279e+03  -1.241 0.214710    
S2:D2:B      -2.556e-03  1.550e-02  2.270e+03  -0.165 0.869054    
S2:D3:B       2.508e-02  1.529e-02  2.262e+03   1.641 0.101006    
S2:D4:B       1.354e-02  1.529e-02  2.262e+03   0.886 0.375858    
S2:D5:B       2.943e-03  1.529e-02  2.262e+03   0.193 0.847358    
S2:D6:B       1.136e-02  1.554e-02  2.270e+03   0.731 0.464758    
S2:D7:B       2.537e-02  1.529e-02  2.262e+03   1.660 0.097092 .  
S2:D8:B       1.004e-02  1.529e-02  2.262e+03   0.657 0.511380    
S2:D9:B       2.069e-02  1.529e-02  2.262e+03   1.353 0.176071    
S2:D10:B      1.551e-02  1.577e-02  2.279e+03   0.983 0.325480    
AG3:S2:D2:B   1.684e-02  2.185e-02  2.267e+03   0.771 0.440799    
AG3:S2:D3:B  -4.229e-02  2.170e-02  2.263e+03  -1.949 0.051390 .  
AG3:S2:D4:B  -1.034e-03  2.162e-02  2.262e+03  -0.048 0.961866    
AG3:S2:D5:B   1.490e-02  2.162e-02  2.262e+03   0.689 0.490743    
AG3:S2:D6:B  -1.058e-02  2.195e-02  2.270e+03  -0.482 0.629961    
AG3:S2:D7:B   2.611e-03  2.177e-02  2.267e+03   0.120 0.904549    
AG3:S2:D8:B  -2.898e-03  2.177e-02  2.267e+03  -0.133 0.894133    
AG3:S2:D9:B  -2.572e-02  2.177e-02  2.267e+03  -1.181 0.237691    
AG3:S2:D10:B  1.276e-02  2.212e-02  2.276e+03   0.577 0.564095    
---
Signif. codes:  0 ‘***’ 0.001 ‘**’ 0.01 ‘*’ 0.05 ‘.’ 0.1 ‘ ’ 1
```


```
Correlation matrix not shown by default, as p = 80 > 12.
Use print(x, correlation=TRUE)  or
    vcov(x)        if you need it
```


```
anova(m_acc)
```


```
Type III Analysis of Variance Table with Satterthwaite's method
          Sum Sq Mean Sq NumDF   DenDF F value    Pr(>F)    
AG       0.35867 0.35867     1   40.05 35.0804 6.041e-07 ***
S        0.22802 0.22802     1   40.05 22.3014 2.848e-05 ***
D        0.85854 0.09539     9 2267.87  9.3301 4.957e-14 ***
B        0.37106 0.37106     1   40.50 36.2916 4.176e-07 ***
AG:S     0.04721 0.04721     1   40.05  4.6175   0.03775 *  
AG:D     0.20414 0.02268     9 2267.87  2.2184   0.01848 *  
S:D      0.21081 0.02342     9 2267.87  2.2909   0.01477 *  
AG:B     0.05750 0.05750     1   40.50  5.6242   0.02255 *  
S:B      0.04405 0.04405     1   40.50  4.3088   0.04431 *  
D:B      0.23042 0.02560     9 2268.53  2.5041   0.00753 ** 
AG:S:D   0.17753 0.01973     9 2267.87  1.9292   0.04388 *  
AG:S:B   0.00117 0.00117     1   40.50  0.1140   0.73740    
AG:D:B   0.08453 0.00939     9 2268.53  0.9186   0.50770    
S:D:B    0.10177 0.01131     9 2268.53  1.1060   0.35475    
AG:S:D:B 0.13317 0.01480     9 2268.53  1.4472   0.16228    
---
Signif. codes:  0 ‘***’ 0.001 ‘**’ 0.01 ‘*’ 0.05 ‘.’ 0.1 ‘ ’ 1
```


```
eta_squared(m_acc)
```


```
# Effect Size for ANOVA (Type III)

Parameter | Eta2 (partial) |       95% CI
-----------------------------------------
AG        |           0.47 | [0.28, 1.00]
S         |           0.36 | [0.17, 1.00]
D         |           0.04 | [0.02, 1.00]
B         |           0.47 | [0.28, 1.00]
AG:S      |           0.10 | [0.00, 1.00]
AG:D      |       8.73e-03 | [0.00, 1.00]
S:D       |       9.01e-03 | [0.00, 1.00]
AG:B      |           0.12 | [0.01, 1.00]
S:B       |           0.10 | [0.00, 1.00]
D:B       |       9.84e-03 | [0.00, 1.00]
AG:S:D    |       7.60e-03 | [0.00, 1.00]
AG:S:B    |       2.81e-03 | [0.00, 1.00]
AG:D:B    |       3.63e-03 | [0.00, 1.00]
S:D:B     |       4.37e-03 | [0.00, 1.00]
AG:S:D:B  |       5.71e-03 | [0.00, 1.00]

- One-sided CIs: upper bound fixed at [1.00].
```

###### Note:

As there we significant differences related to the age groups and
also due to our original hypothesis, we will test the effect of age
among the placebo groups and the effect of stimulation within each age
group.

###### Age-related differences in accuracy


```
data_subset <- subset(df_acc, D != 0 & S == 2)
data_subset <- droplevels(data_subset)
levels(data_subset$D)
```


```
 [1] "1"  "2"  "3"  "4"  "5"  "6"  "7"  "8"  "9"  "10"
```


```
levels(data_subset$AG)
```


```
[1] "2" "3"
```


```
levels(data_subset$S)
```


```
[1] "2"
```


```
levels(data_subset$Group)
```


```
[1] "Middle-Placebo" "Older-Placebo"
```


```
m_acc_placebo <- lmer(formula = Acc ~ AG*D*B + (1 + B|Code), data=data_subset)
summary(m_acc_placebo)
```


```
Linear mixed model fit by REML. t-tests use Satterthwaite's method ['lmerModLmerTest']
Formula: Acc ~ AG * D * B + (1 + B | Code)
   Data: data_subset

REML criterion at convergence: -2173.8

Scaled residuals: 
    Min      1Q  Median      3Q     Max 
-7.5053 -0.4983  0.1143  0.6325  2.9431 

Random effects:
 Groups   Name        Variance  Std.Dev. Corr 
 Code     (Intercept) 1.701e-03 0.041244      
          B           1.104e-05 0.003323 -0.14
 Residual             6.958e-03 0.083412      
Number of obs: 1172, groups:  Code, 20

Fixed effects:
              Estimate Std. Error         df t value Pr(>|t|)    
(Intercept)  9.018e-01  2.780e-02  1.837e+02  32.434  < 2e-16 ***
AG3         -1.320e-01  3.932e-02  1.837e+02  -3.358 0.000955 ***
D2          -3.398e-02  3.473e-02  1.096e+03  -0.978 0.328067    
D3          -1.532e-02  3.473e-02  1.096e+03  -0.441 0.659220    
D4           1.918e-03  3.473e-02  1.096e+03   0.055 0.955965    
D5           1.070e-02  3.473e-02  1.096e+03   0.308 0.758070    
D6           6.272e-03  3.473e-02  1.096e+03   0.181 0.856711    
D7           1.883e-03  3.473e-02  1.096e+03   0.054 0.956759    
D8           9.411e-03  3.473e-02  1.096e+03   0.271 0.786447    
D9           1.893e-02  3.473e-02  1.096e+03   0.545 0.585864    
D10          2.709e-02  3.473e-02  1.096e+03   0.780 0.435431    
B           -2.952e-03  6.392e-03  6.342e+02  -0.462 0.644380    
AG3:D2       1.650e-02  4.964e-02  1.098e+03   0.332 0.739661    
AG3:D3       8.326e-02  4.922e-02  1.097e+03   1.692 0.090987 .  
AG3:D4       3.881e-02  4.911e-02  1.096e+03   0.790 0.429594    
AG3:D5       5.261e-02  4.911e-02  1.096e+03   1.071 0.284309    
AG3:D6       8.741e-02  4.911e-02  1.096e+03   1.780 0.075389 .  
AG3:D7       8.813e-02  4.982e-02  1.101e+03   1.769 0.077187 .  
AG3:D8       1.158e-01  4.982e-02  1.101e+03   2.324 0.020316 *  
AG3:D9       9.531e-02  4.982e-02  1.101e+03   1.913 0.055997 .  
AG3:D10      5.879e-02  4.982e-02  1.101e+03   1.180 0.238204    
AG3:B        1.970e-02  9.040e-03  6.342e+02   2.180 0.029649 *  
D2:B         1.444e-02  8.917e-03  1.096e+03   1.620 0.105558    
D3:B         1.179e-02  8.917e-03  1.096e+03   1.322 0.186402    
D4:B         7.245e-03  8.917e-03  1.096e+03   0.812 0.416687    
D5:B         3.964e-04  8.917e-03  1.096e+03   0.044 0.964554    
D6:B         5.647e-03  8.917e-03  1.096e+03   0.633 0.526672    
D7:B         9.553e-03  8.917e-03  1.096e+03   1.071 0.284293    
D8:B         3.508e-03  8.917e-03  1.096e+03   0.393 0.694094    
D9:B         3.772e-03  8.917e-03  1.096e+03   0.423 0.672402    
D10:B        2.405e-03  8.917e-03  1.096e+03   0.270 0.787439    
AG3:D2:B    -1.047e-02  1.270e-02  1.098e+03  -0.824 0.409893    
AG3:D3:B    -2.164e-02  1.270e-02  1.098e+03  -1.704 0.088685 .  
AG3:D4:B    -1.194e-02  1.261e-02  1.096e+03  -0.947 0.344002    
AG3:D5:B    -5.680e-03  1.261e-02  1.096e+03  -0.450 0.652475    
AG3:D6:B    -1.980e-02  1.261e-02  1.096e+03  -1.570 0.116763    
AG3:D7:B    -1.622e-02  1.279e-02  1.102e+03  -1.268 0.204970    
AG3:D8:B    -2.313e-02  1.279e-02  1.102e+03  -1.809 0.070780 .  
AG3:D9:B    -1.894e-02  1.279e-02  1.102e+03  -1.481 0.138828    
AG3:D10:B   -6.731e-03  1.279e-02  1.102e+03  -0.526 0.598753    
---
Signif. codes:  0 ‘***’ 0.001 ‘**’ 0.01 ‘*’ 0.05 ‘.’ 0.1 ‘ ’ 1
```


```
Correlation matrix not shown by default, as p = 40 > 12.
Use print(x, correlation=TRUE)  or
    vcov(x)        if you need it
```


```
anova(m_acc_placebo)
```


```
Type III Analysis of Variance Table with Satterthwaite's method
         Sum Sq  Mean Sq NumDF   DenDF F value   Pr(>F)   
AG     0.070019 0.070019     1   18.03 10.0637 0.005265 **
D      0.177761 0.019751     9 1098.75  2.8388 0.002620 **
B      0.097901 0.097901     1   18.16 14.0711 0.001443 **
AG:D   0.068963 0.007663     9 1098.75  1.1013 0.358739   
AG:B   0.026129 0.026129     1   18.16  3.7555 0.068341 . 
D:B    0.069109 0.007679     9 1099.04  1.1036 0.357030   
AG:D:B 0.045916 0.005102     9 1099.04  0.7333 0.678631   
---
Signif. codes:  0 ‘***’ 0.001 ‘**’ 0.01 ‘*’ 0.05 ‘.’ 0.1 ‘ ’ 1
```


```
eta_squared(m_acc_placebo)
```


```
# Effect Size for ANOVA (Type III)

Parameter | Eta2 (partial) |       95% CI
-----------------------------------------
AG        |           0.36 | [0.08, 1.00]
D         |           0.02 | [0.00, 1.00]
B         |           0.44 | [0.15, 1.00]
AG:D      |       8.94e-03 | [0.00, 1.00]
AG:B      |           0.17 | [0.00, 1.00]
D:B       |       8.96e-03 | [0.00, 1.00]
AG:D:B    |       5.97e-03 | [0.00, 1.00]

- One-sided CIs: upper bound fixed at [1.00].
```

###### Accuracy in groups of middle-aged adults


```
data_subset <- subset(df_acc, D != 0 & AG == 2)
data_subset <- droplevels(data_subset)
levels(data_subset$D)
```


```
 [1] "1"  "2"  "3"  "4"  "5"  "6"  "7"  "8"  "9"  "10"
```


```
levels(data_subset$AG)
```


```
[1] "2"
```


```
levels(data_subset$S)
```


```
[1] "1" "2"
```


```
levels(data_subset$Group)
```


```
[1] "Middle-Placebo" "Middle-Verum"
```


```
m_acc_middle <- lmer(formula = Acc ~ S*D*B + (1 + B|Code), data=data_subset)
summary(m_acc_middle)
```


```
Linear mixed model fit by REML. t-tests use Satterthwaite's method ['lmerModLmerTest']
Formula: Acc ~ S * D * B + (1 + B | Code)
   Data: data_subset

REML criterion at convergence: -2223

Scaled residuals: 
    Min      1Q  Median      3Q     Max 
-6.6728 -0.4582  0.1088  0.5747  3.1238 

Random effects:
 Groups   Name        Variance  Std.Dev. Corr 
 Code     (Intercept) 3.031e-03 0.055055      
          B           3.118e-05 0.005584 -0.37
 Residual             6.613e-03 0.081318      
Number of obs: 1175, groups:  Code, 20

Fixed effects:
              Estimate Std. Error         df t value Pr(>|t|)    
(Intercept)  7.643e-01  2.960e-02  1.018e+02  25.820  < 2e-16 ***
S2           1.375e-01  4.186e-02  1.018e+02   3.286  0.00140 ** 
D2          -1.038e-02  3.484e-02  1.104e+03  -0.298  0.76575    
D3           1.363e-01  3.386e-02  1.099e+03   4.027 6.04e-05 ***
D4           1.009e-01  3.386e-02  1.099e+03   2.980  0.00295 ** 
D5           8.204e-02  3.386e-02  1.099e+03   2.423  0.01554 *  
D6           9.315e-02  3.511e-02  1.104e+03   2.653  0.00808 ** 
D7           1.480e-01  3.386e-02  1.099e+03   4.372 1.35e-05 ***
D8           9.447e-02  3.386e-02  1.099e+03   2.790  0.00536 ** 
D9           1.507e-01  3.386e-02  1.099e+03   4.452 9.36e-06 ***
D10          1.427e-01  3.603e-02  1.109e+03   3.962 7.92e-05 ***
B            1.456e-02  6.396e-03  4.422e+02   2.276  0.02333 *  
S2:D2       -2.360e-02  4.858e-02  1.102e+03  -0.486  0.62725    
S2:D3       -1.517e-01  4.788e-02  1.099e+03  -3.167  0.00158 ** 
S2:D4       -9.897e-02  4.788e-02  1.099e+03  -2.067  0.03895 *  
S2:D5       -7.134e-02  4.788e-02  1.099e+03  -1.490  0.13650    
S2:D6       -8.688e-02  4.877e-02  1.102e+03  -1.781  0.07513 .  
S2:D7       -1.461e-01  4.788e-02  1.099e+03  -3.052  0.00232 ** 
S2:D8       -8.506e-02  4.788e-02  1.099e+03  -1.776  0.07593 .  
S2:D9       -1.318e-01  4.788e-02  1.099e+03  -2.753  0.00600 ** 
S2:D10      -1.156e-01  4.944e-02  1.105e+03  -2.339  0.01952 *  
S2:B        -1.751e-02  9.045e-03  4.422e+02  -1.936  0.05355 .  
D2:B         1.698e-02  8.943e-03  1.105e+03   1.898  0.05789 .  
D3:B        -1.329e-02  8.693e-03  1.099e+03  -1.529  0.12656    
D4:B        -6.295e-03  8.693e-03  1.099e+03  -0.724  0.46913    
D5:B        -2.547e-03  8.693e-03  1.099e+03  -0.293  0.76962    
D6:B        -5.407e-03  8.978e-03  1.105e+03  -0.602  0.54716    
D7:B        -1.582e-02  8.693e-03  1.099e+03  -1.820  0.06904 .  
D8:B        -6.532e-03  8.693e-03  1.099e+03  -0.751  0.45255    
D9:B        -1.692e-02  8.693e-03  1.099e+03  -1.946  0.05190 .  
D10:B       -1.368e-02  9.245e-03  1.111e+03  -1.480  0.13910    
S2:D2:B     -2.533e-03  1.247e-02  1.102e+03  -0.203  0.83907    
S2:D3:B      2.508e-02  1.229e-02  1.099e+03   2.040  0.04158 *  
S2:D4:B      1.354e-02  1.229e-02  1.099e+03   1.101  0.27098    
S2:D5:B      2.943e-03  1.229e-02  1.099e+03   0.239  0.81086    
S2:D6:B      1.105e-02  1.250e-02  1.102e+03   0.885  0.37661    
S2:D7:B      2.537e-02  1.229e-02  1.099e+03   2.064  0.03926 *  
S2:D8:B      1.004e-02  1.229e-02  1.099e+03   0.817  0.41428    
S2:D9:B      2.069e-02  1.229e-02  1.099e+03   1.683  0.09269 .  
S2:D10:B     1.609e-02  1.269e-02  1.106e+03   1.268  0.20512    
---
Signif. codes:  0 ‘***’ 0.001 ‘**’ 0.01 ‘*’ 0.05 ‘.’ 0.1 ‘ ’ 1
```


```
Correlation matrix not shown by default, as p = 40 > 12.
Use print(x, correlation=TRUE)  or
    vcov(x)        if you need it
```


```
anova(m_acc_middle)
```


```
Type III Analysis of Variance Table with Satterthwaite's method
        Sum Sq  Mean Sq NumDF   DenDF F value   Pr(>F)    
S     0.019693 0.019693     1   17.94  2.9781 0.101581    
D     0.240611 0.026735     9 1101.02  4.0430 4.11e-05 ***
B     0.058554 0.058554     1   17.82  8.8548 0.008168 ** 
S:D   0.126867 0.014096     9 1101.02  2.1317 0.024516 *  
S:B   0.013182 0.013182     1   17.82  1.9934 0.175214    
D:B   0.111251 0.012361     9 1101.40  1.8693 0.052782 .  
S:D:B 0.076313 0.008479     9 1101.40  1.2823 0.241934    
---
Signif. codes:  0 ‘***’ 0.001 ‘**’ 0.01 ‘*’ 0.05 ‘.’ 0.1 ‘ ’ 1
```


```
eta_squared(m_acc_middle)
```


```
# Effect Size for ANOVA (Type III)

Parameter | Eta2 (partial) |       95% CI
-----------------------------------------
S         |           0.14 | [0.00, 1.00]
D         |           0.03 | [0.01, 1.00]
B         |           0.33 | [0.06, 1.00]
S:D       |           0.02 | [0.00, 1.00]
S:B       |           0.10 | [0.00, 1.00]
D:B       |           0.02 | [0.00, 1.00]
S:D:B     |           0.01 | [0.00, 1.00]

- One-sided CIs: upper bound fixed at [1.00].
```

###### Accuracy in the groups of older adults


```
data_subset <- subset(df_acc, D != 0 & AG == 3)
data_subset <- droplevels(data_subset)
levels(data_subset$D)
```


```
 [1] "1"  "2"  "3"  "4"  "5"  "6"  "7"  "8"  "9"  "10"
```


```
levels(data_subset$AG)
```


```
[1] "3"
```


```
levels(data_subset$S)
```


```
[1] "1" "2"
```


```
levels(data_subset$Group)
```


```
[1] "Older-Placebo" "Older-Verum"
```


```
m_acc_older <- lmer(formula = Acc ~ S*D*B + (1 + B|Code), data=data_subset)
summary(m_acc_older)
```


```
Linear mixed model fit by REML. t-tests use Satterthwaite's method ['lmerModLmerTest']
Formula: Acc ~ S * D * B + (1 + B | Code)
   Data: data_subset

REML criterion at convergence: -1329.1

Scaled residuals: 
    Min      1Q  Median      3Q     Max 
-5.2052 -0.5173  0.1006  0.6102  4.3695 

Random effects:
 Groups   Name        Variance  Std.Dev. Corr
 Code     (Intercept) 2.372e-03 0.048698     
          B           5.212e-05 0.007219 0.33
 Residual             1.454e-02 0.120579     
Number of obs: 1166, groups:  Code, 20

Fixed effects:
              Estimate Std. Error         df t value Pr(>|t|)    
(Intercept)  5.381e-01  3.869e-02  2.606e+02  13.907  < 2e-16 ***
S2           2.317e-01  5.472e-02  2.606e+02   4.234 3.18e-05 ***
D2           1.386e-01  5.020e-02  1.091e+03   2.761 0.005856 ** 
D3           5.764e-02  5.020e-02  1.091e+03   1.148 0.251124    
D4           1.678e-01  5.020e-02  1.091e+03   3.343 0.000856 ***
D5           2.017e-01  5.020e-02  1.091e+03   4.017 6.29e-05 ***
D6           1.577e-01  5.162e-02  1.098e+03   3.056 0.002300 ** 
D7           2.501e-01  5.020e-02  1.091e+03   4.982 7.32e-07 ***
D8           2.690e-01  5.020e-02  1.091e+03   5.359 1.02e-07 ***
D9           1.959e-01  5.020e-02  1.091e+03   3.902 0.000101 ***
D10          2.996e-01  5.020e-02  1.091e+03   5.967 3.25e-09 ***
B            3.278e-02  9.397e-03  5.060e+02   3.489 0.000528 ***
S2:D2       -1.556e-01  7.176e-02  1.093e+03  -2.169 0.030296 *  
S2:D3        1.047e-02  7.115e-02  1.091e+03   0.147 0.882999    
S2:D4       -1.271e-01  7.099e-02  1.091e+03  -1.790 0.073677 .  
S2:D5       -1.384e-01  7.099e-02  1.091e+03  -1.949 0.051556 .  
S2:D6       -6.406e-02  7.201e-02  1.095e+03  -0.890 0.373854    
S2:D7       -1.595e-01  7.201e-02  1.095e+03  -2.215 0.026993 *  
S2:D8       -1.432e-01  7.201e-02  1.095e+03  -1.989 0.046958 *  
S2:D9       -8.106e-02  7.201e-02  1.095e+03  -1.126 0.260580    
S2:D10      -2.131e-01  7.201e-02  1.095e+03  -2.959 0.003153 ** 
S2:B        -1.603e-02  1.329e-02  5.060e+02  -1.206 0.228272    
D2:B        -1.030e-02  1.289e-02  1.091e+03  -0.799 0.424565    
D3:B         7.312e-03  1.289e-02  1.091e+03   0.567 0.570643    
D4:B        -1.720e-02  1.289e-02  1.091e+03  -1.334 0.182377    
D5:B        -2.313e-02  1.289e-02  1.091e+03  -1.794 0.073058 .  
D6:B        -1.480e-02  1.325e-02  1.098e+03  -1.117 0.264366    
D7:B        -3.474e-02  1.289e-02  1.091e+03  -2.695 0.007151 ** 
D8:B        -2.685e-02  1.289e-02  1.091e+03  -2.083 0.037492 *  
D9:B        -1.023e-02  1.289e-02  1.091e+03  -0.793 0.427683    
D10:B       -3.268e-02  1.289e-02  1.091e+03  -2.535 0.011370 *  
S2:D2:B      1.418e-02  1.835e-02  1.092e+03   0.773 0.439877    
S2:D3:B     -1.724e-02  1.836e-02  1.092e+03  -0.939 0.347840    
S2:D4:B      1.251e-02  1.823e-02  1.091e+03   0.686 0.492829    
S2:D5:B      1.784e-02  1.823e-02  1.091e+03   0.979 0.327881    
S2:D6:B      6.515e-04  1.849e-02  1.095e+03   0.035 0.971898    
S2:D7:B      2.790e-02  1.849e-02  1.096e+03   1.509 0.131616    
S2:D8:B      7.055e-03  1.849e-02  1.096e+03   0.382 0.702837    
S2:D9:B     -5.116e-03  1.849e-02  1.096e+03  -0.277 0.782075    
S2:D10:B     2.818e-02  1.849e-02  1.096e+03   1.524 0.127708    
---
Signif. codes:  0 ‘***’ 0.001 ‘**’ 0.01 ‘*’ 0.05 ‘.’ 0.1 ‘ ’ 1
```


```
Correlation matrix not shown by default, as p = 40 > 12.
Use print(x, correlation=TRUE)  or
    vcov(x)        if you need it
```


```
anova(m_acc_older)
```


```
Type III Analysis of Variance Table with Satterthwaite's method
       Sum Sq Mean Sq NumDF   DenDF F value    Pr(>F)    
S     0.30661 0.30661     1   18.16 21.0884 0.0002215 ***
D     0.82175 0.09131     9 1093.82  6.2800 1.076e-08 ***
B     0.34473 0.34473     1   18.42 23.7104 0.0001158 ***
S:D   0.26315 0.02924     9 1093.82  2.0110 0.0350738 *  
S:B   0.02905 0.02905     1   18.42  1.9978 0.1742088    
D:B   0.20220 0.02247     9 1093.99  1.5452 0.1272651    
S:D:B 0.15849 0.01761     9 1093.99  1.2112 0.2839162    
---
Signif. codes:  0 ‘***’ 0.001 ‘**’ 0.01 ‘*’ 0.05 ‘.’ 0.1 ‘ ’ 1
```


```
eta_squared(m_acc_older)
```


```
# Effect Size for ANOVA (Type III)

Parameter | Eta2 (partial) |       95% CI
-----------------------------------------
S         |           0.54 | [0.25, 1.00]
D         |           0.05 | [0.02, 1.00]
B         |           0.56 | [0.29, 1.00]
S:D       |           0.02 | [0.00, 1.00]
S:B       |           0.10 | [0.00, 1.00]
D:B       |           0.01 | [0.00, 1.00]
S:D:B     |       9.87e-03 | [0.00, 1.00]

- One-sided CIs: upper bound fixed at [1.00].
```

##### Normalized accuracy of execution in the finger-tapping task (i.e., accuracy dynamics)

The following statistical tests concern the normalized accuracy of
execution in the finger-tapping task, quantified as the ratio of correct
to total sequences generated in each training block, divided by the
accuracy of the first training block.


```
acc_norm_file_name <-  "AP3_acc_norm_MS_27_03_2023.txt"
file_path <- file.path(paste(python_data_directory, acc_norm_file_name, sep='/'))
df_acc_norm = read.delim(file_path, header = TRUE, na.strings = "NN")
head(df_acc_norm)
```


```
df_acc_norm$Code <- as.factor(df_acc_norm$Code)
df_acc_norm$AG <- as.factor(df_acc_norm$AG)
df_acc_norm$S <- as.factor(df_acc_norm$S)
df_acc_norm$Group <- as.factor(df_acc_norm$Group)
df_acc_norm$Acc_n <- as.numeric(df_acc_norm$Acc_n)
df_acc_norm$D <- as.factor(df_acc_norm$D)
df_acc_norm$B <- as.numeric(df_acc_norm$B)
levels(df_acc_norm$Code)
```


```
 [1] "217201" "217202" "217203" "217205" "217206" "217207" "217208" "217209" "217210" "217251" "217252" "217253" "217254" "217255" "217257" "217258" "217259" "217260"
[19] "217301" "217302" "217303" "217304" "217305" "217306" "217307" "217308" "217309" "217310" "217311" "217312" "217351" "217352" "217353" "217354" "217355" "217356"
[37] "217357" "217358" "217359" "217360"
```


```
levels(df_acc_norm$AG)
```


```
[1] "2" "3"
```


```
levels(df_acc_norm$S)
```


```
[1] "1" "2"
```


```
levels(df_acc_norm$Group)
```


```
[1] "Middle-Placebo" "Middle-Verum"   "Older-Placebo"  "Older-Verum"
```


```
levels(df_acc_norm$D)
```


```
 [1] "1"  "2"  "3"  "4"  "5"  "6"  "7"  "8"  "9"  "10"
```


###### Select a model to approximate the data


```
m1 <- lm(formula = Acc_n ~ AG*S*D*B, data=df_acc_norm)
m2 <- lmer(formula = Acc_n ~ AG*S*D*B + (1 | Code), data=df_acc_norm)
BIC(m1, m2)
```


```
m3 <- lmer(formula = Acc_n ~ AG*S*D*B + (1 + B|Code), data=df_acc_norm)
```


```
boundary (singular) fit: see help('isSingular')
```


```
anova(m2, m3)
```


```
refitting model(s) with ML (instead of REML)
```


```
Data: df_acc_norm
Models:
m2: Acc_n ~ AG * S * D * B + (1 | Code)
m3: Acc_n ~ AG * S * D * B + (1 + B | Code)
   npar    AIC   BIC  logLik deviance  Chisq Df Pr(>Chisq)    
m2   82 315.22 787.4 -75.608   151.22                         
m3   84 300.50 784.2 -66.252   132.50 18.713  2  8.641e-05 ***
---
Signif. codes:  0 ‘***’ 0.001 ‘**’ 0.01 ‘*’ 0.05 ‘.’ 0.1 ‘ ’ 1
```

###### Model selection

Based on the BIC, the better model is m3, including random intercepts
and slopes for individuals. However, this model was flagged as singular
and the change in BIC is small, so we will use m2.


```
m_acc_norm <- lmer(formula = Acc_n ~ AG*S*D*B + (1 | Code), data=df_acc_norm)
summary(m_acc_norm)
```


```
Linear mixed model fit by REML. t-tests use Satterthwaite's method ['lmerModLmerTest']
Formula: Acc_n ~ AG * S * D * B + (1 | Code)
   Data: df_acc_norm

REML criterion at convergence: 576.4

Scaled residuals: 
     Min       1Q   Median       3Q      Max 
-16.8067  -0.2832   0.0453   0.3227   6.0865 

Random effects:
 Groups   Name        Variance Std.Dev.
 Code     (Intercept) 0.51442  0.7172  
 Residual             0.05812  0.2411  
Number of obs: 2341, groups:  Code, 40

Fixed effects:
               Estimate Std. Error         df t value Pr(>|t|)    
(Intercept)   9.850e-01  2.377e-01  4.323e+01   4.145 0.000156 ***
AG3           2.278e-01  3.361e-01  4.323e+01   0.678 0.501518    
S2            2.622e-02  3.361e-01  4.323e+01   0.078 0.938184    
D2            1.465e-02  1.032e-01  2.225e+03   0.142 0.887073    
D3            2.275e-01  1.004e-01  2.225e+03   2.266 0.023538 *  
D4            1.750e-01  1.004e-01  2.225e+03   1.744 0.081293 .  
D5            1.390e-01  1.004e-01  2.225e+03   1.385 0.166238    
D6            1.731e-01  1.040e-01  2.225e+03   1.665 0.096065 .  
D7            2.359e-01  1.004e-01  2.225e+03   2.351 0.018828 *  
D8            1.598e-01  1.004e-01  2.225e+03   1.592 0.111474    
D9            2.364e-01  1.004e-01  2.225e+03   2.355 0.018596 *  
D10           2.251e-01  1.066e-01  2.225e+03   2.112 0.034782 *  
B             2.438e-02  1.822e-02  2.225e+03   1.338 0.181036    
AG3:S2       -2.473e-01  4.753e-01  4.323e+01  -0.520 0.605496    
AG3:D2        6.006e-01  1.439e-01  2.225e+03   4.172 3.13e-05 ***
AG3:D3        3.070e-01  1.419e-01  2.225e+03   2.163 0.030654 *  
AG3:D4        4.168e-01  1.419e-01  2.225e+03   2.936 0.003354 ** 
AG3:D5        6.550e-01  1.419e-01  2.225e+03   4.615 4.16e-06 ***
AG3:D6        4.772e-01  1.465e-01  2.225e+03   3.258 0.001140 ** 
AG3:D7        5.764e-01  1.419e-01  2.225e+03   4.061 5.05e-05 ***
AG3:D8        7.335e-01  1.419e-01  2.225e+03   5.167 2.59e-07 ***
AG3:D9        3.974e-01  1.419e-01  2.225e+03   2.800 0.005160 ** 
AG3:D10       7.396e-01  1.464e-01  2.225e+03   5.052 4.73e-07 ***
S2:D2        -4.976e-02  1.439e-01  2.225e+03  -0.346 0.729628    
S2:D3        -2.390e-01  1.419e-01  2.225e+03  -1.684 0.092333 .  
S2:D4        -1.655e-01  1.419e-01  2.225e+03  -1.166 0.243753    
S2:D5        -1.202e-01  1.419e-01  2.225e+03  -0.847 0.397232    
S2:D6        -1.635e-01  1.445e-01  2.225e+03  -1.131 0.258045    
S2:D7        -2.299e-01  1.419e-01  2.225e+03  -1.619 0.105501    
S2:D8        -1.479e-01  1.419e-01  2.225e+03  -1.042 0.297544    
S2:D9        -2.098e-01  1.419e-01  2.225e+03  -1.478 0.139588    
S2:D10       -1.879e-01  1.464e-01  2.225e+03  -1.283 0.199535    
AG3:B         9.937e-02  2.577e-02  2.225e+03   3.856 0.000119 ***
S2:B         -2.700e-02  2.577e-02  2.225e+03  -1.048 0.294857    
D2:B          1.872e-02  2.648e-02  2.225e+03   0.707 0.479740    
D3:B         -2.566e-02  2.577e-02  2.225e+03  -0.996 0.319580    
D4:B         -1.456e-02  2.577e-02  2.225e+03  -0.565 0.572290    
D5:B         -7.561e-03  2.577e-02  2.225e+03  -0.293 0.769277    
D6:B         -1.696e-02  2.658e-02  2.225e+03  -0.638 0.523560    
D7:B         -2.703e-02  2.577e-02  2.225e+03  -1.049 0.294380    
D8:B         -1.385e-02  2.577e-02  2.225e+03  -0.537 0.591089    
D9:B         -2.824e-02  2.577e-02  2.225e+03  -1.096 0.273280    
D10:B        -2.189e-02  2.734e-02  2.225e+03  -0.801 0.423280    
AG3:S2:D2    -5.711e-01  2.032e-01  2.225e+03  -2.810 0.004995 ** 
AG3:S2:D3    -1.840e-01  2.010e-01  2.225e+03  -0.915 0.360068    
AG3:S2:D4    -3.468e-01  2.007e-01  2.225e+03  -1.728 0.084162 .  
AG3:S2:D5    -5.473e-01  2.007e-01  2.225e+03  -2.726 0.006452 ** 
AG3:S2:D6    -3.338e-01  2.040e-01  2.225e+03  -1.637 0.101837    
AG3:S2:D7    -4.355e-01  2.022e-01  2.225e+03  -2.154 0.031349 *  
AG3:S2:D8    -5.536e-01  2.022e-01  2.225e+03  -2.738 0.006230 ** 
AG3:S2:D9    -2.477e-01  2.022e-01  2.225e+03  -1.225 0.220620    
AG3:S2:D10   -6.416e-01  2.053e-01  2.225e+03  -3.125 0.001804 ** 
AG3:S2:B     -7.390e-02  3.645e-02  2.225e+03  -2.028 0.042727 *  
AG3:D2:B     -8.473e-02  3.695e-02  2.225e+03  -2.293 0.021939 *  
AG3:D3:B     -2.797e-02  3.645e-02  2.225e+03  -0.767 0.442876    
AG3:D4:B     -5.807e-02  3.645e-02  2.225e+03  -1.593 0.111250    
AG3:D5:B     -1.156e-01  3.645e-02  2.225e+03  -3.172 0.001535 ** 
AG3:D6:B     -6.798e-02  3.752e-02  2.225e+03  -1.812 0.070164 .  
AG3:D7:B     -9.000e-02  3.645e-02  2.225e+03  -2.469 0.013616 *  
AG3:D8:B     -9.308e-02  3.645e-02  2.225e+03  -2.554 0.010724 *  
AG3:D9:B     -1.604e-02  3.645e-02  2.225e+03  -0.440 0.659847    
AG3:D10:B    -9.869e-02  3.757e-02  2.225e+03  -2.627 0.008676 ** 
S2:D2:B      -2.886e-03  3.695e-02  2.225e+03  -0.078 0.937755    
S2:D3:B       3.796e-02  3.645e-02  2.225e+03   1.041 0.297767    
S2:D4:B       2.136e-02  3.645e-02  2.225e+03   0.586 0.557983    
S2:D5:B       6.620e-03  3.645e-02  2.225e+03   0.182 0.855889    
S2:D6:B       2.292e-02  3.703e-02  2.225e+03   0.619 0.536047    
S2:D7:B       3.731e-02  3.645e-02  2.225e+03   1.024 0.306172    
S2:D8:B       1.761e-02  3.645e-02  2.225e+03   0.483 0.629123    
S2:D9:B       3.167e-02  3.645e-02  2.225e+03   0.869 0.385026    
S2:D10:B      2.331e-02  3.757e-02  2.225e+03   0.621 0.534967    
AG3:S2:D2:B   7.474e-02  5.208e-02  2.225e+03   1.435 0.151358    
AG3:S2:D3:B   1.035e-03  5.173e-02  2.225e+03   0.020 0.984032    
AG3:S2:D4:B   4.381e-02  5.155e-02  2.225e+03   0.850 0.395481    
AG3:S2:D5:B   1.039e-01  5.155e-02  2.225e+03   2.015 0.044035 *  
AG3:S2:D6:B   4.099e-02  5.231e-02  2.225e+03   0.784 0.433314    
AG3:S2:D7:B   6.904e-02  5.190e-02  2.225e+03   1.330 0.183603    
AG3:S2:D8:B   6.153e-02  5.190e-02  2.225e+03   1.185 0.235988    
AG3:S2:D9:B  -9.543e-03  5.190e-02  2.225e+03  -0.184 0.854140    
AG3:S2:D10:B  8.984e-02  5.270e-02  2.225e+03   1.705 0.088365 .  
---
Signif. codes:  0 ‘***’ 0.001 ‘**’ 0.01 ‘*’ 0.05 ‘.’ 0.1 ‘ ’ 1
```


```
Correlation matrix not shown by default, as p = 80 > 12.
Use print(x, correlation=TRUE)  or
    vcov(x)        if you need it
```


```
anova(m_acc_norm)
```


```
Type III Analysis of Variance Table with Satterthwaite's method
         Sum Sq Mean Sq NumDF   DenDF F value    Pr(>F)    
AG       0.1803 0.18025     1   36.59  3.1013 0.0865857 .  
S        0.2184 0.21837     1   36.59  3.7572 0.0603247 .  
D        3.9676 0.44085     9 2225.01  7.5850 5.039e-11 ***
B        2.0730 2.07298     1 2225.01 35.6667 2.718e-09 ***
AG:S     0.1122 0.11221     1   36.59  1.9306 0.1730938    
AG:D     1.7064 0.18960     9 2225.01  3.2622 0.0005965 ***
S:D      1.8792 0.20880     9 2225.01  3.5924 0.0001876 ***
AG:B     0.7478 0.74781     1 2225.01 12.8665 0.0003417 ***
S:B      0.7220 0.72202     1 2225.01 12.4227 0.0004327 ***
D:B      1.0547 0.11719     9 2225.01  2.0163 0.0340147 *  
AG:S:D   1.0394 0.11549     9 2225.01  1.9871 0.0370696 *  
AG:S:B   0.2957 0.29569     1 2225.01  5.0875 0.0241951 *  
AG:D:B   0.6422 0.07136     9 2225.01  1.2277 0.2730170    
S:D:B    0.6743 0.07493     9 2225.01  1.2892 0.2373977    
AG:S:D:B 0.6029 0.06699     9 2225.01  1.1526 0.3217352    
---
Signif. codes:  0 ‘***’ 0.001 ‘**’ 0.01 ‘*’ 0.05 ‘.’ 0.1 ‘ ’ 1
```


```
eta_squared(m_acc_norm)
```


```
# Effect Size for ANOVA (Type III)

Parameter | Eta2 (partial) |       95% CI
-----------------------------------------
AG        |           0.08 | [0.00, 1.00]
S         |           0.09 | [0.00, 1.00]
D         |           0.03 | [0.02, 1.00]
B         |           0.02 | [0.01, 1.00]
AG:S      |           0.05 | [0.00, 1.00]
AG:D      |           0.01 | [0.00, 1.00]
S:D       |           0.01 | [0.00, 1.00]
AG:B      |       5.75e-03 | [0.00, 1.00]
S:B       |       5.55e-03 | [0.00, 1.00]
D:B       |       8.09e-03 | [0.00, 1.00]
AG:S:D    |       7.97e-03 | [0.00, 1.00]
AG:S:B    |       2.28e-03 | [0.00, 1.00]
AG:D:B    |       4.94e-03 | [0.00, 1.00]
S:D:B     |       5.19e-03 | [0.00, 1.00]
AG:S:D:B  |       4.64e-03 | [0.00, 1.00]

- One-sided CIs: upper bound fixed at [1.00].
```

###### Differences in accuracy dynamics between placebo groups


```
data_subset <- subset(df_acc_norm, D != 0 & S == 2)
data_subset <- droplevels(data_subset)
levels(data_subset$D)
```


```
 [1] "1"  "2"  "3"  "4"  "5"  "6"  "7"  "8"  "9"  "10"
```


```
levels(data_subset$AG)
```


```
[1] "2" "3"
```


```
levels(data_subset$S)
```


```
[1] "2"
```


```
levels(data_subset$Group)
```


```
[1] "Middle-Placebo" "Older-Placebo"
```


```
m_acc_norm_placebo <- lmer(formula = Acc_n ~ AG*D*B + (1 | Code), data=data_subset)
summary(m_acc_norm_placebo)
```


```
Linear mixed model fit by REML. t-tests use Satterthwaite's method ['lmerModLmerTest']
Formula: Acc_n ~ AG * D * B + (1 | Code)
   Data: data_subset

REML criterion at convergence: -1613.7

Scaled residuals: 
    Min      1Q  Median      3Q     Max 
-6.1132 -0.4676  0.0782  0.5809  2.8333 

Random effects:
 Groups   Name        Variance Std.Dev.
 Code     (Intercept) 0.02468  0.1571  
 Residual             0.01107  0.1052  
Number of obs: 1172, groups:  Code, 20

Fixed effects:
              Estimate Std. Error         df t value Pr(>|t|)    
(Intercept)  1.011e+00  5.854e-02  3.408e+01  17.273  < 2e-16 ***
AG3         -1.951e-02  8.279e-02  3.408e+01  -0.236  0.81510    
D2          -3.510e-02  4.380e-02  1.114e+03  -0.801  0.42309    
D3          -1.157e-02  4.380e-02  1.114e+03  -0.264  0.79168    
D4           9.545e-03  4.380e-02  1.114e+03   0.218  0.82755    
D5           1.881e-02  4.380e-02  1.114e+03   0.429  0.66775    
D6           9.616e-03  4.380e-02  1.114e+03   0.220  0.82628    
D7           6.066e-03  4.380e-02  1.114e+03   0.138  0.88988    
D8           1.191e-02  4.380e-02  1.114e+03   0.272  0.78571    
D9           2.662e-02  4.380e-02  1.114e+03   0.608  0.54343    
D10          3.725e-02  4.380e-02  1.114e+03   0.850  0.39526    
B           -2.620e-03  7.953e-03  1.114e+03  -0.329  0.74187    
AG3:D2       2.960e-02  6.261e-02  1.114e+03   0.473  0.63650    
AG3:D3       1.232e-01  6.208e-02  1.114e+03   1.984  0.04752 *  
AG3:D4       6.998e-02  6.195e-02  1.114e+03   1.130  0.25887    
AG3:D5       1.077e-01  6.195e-02  1.114e+03   1.739  0.08235 .  
AG3:D6       1.434e-01  6.195e-02  1.114e+03   2.314  0.02083 *  
AG3:D7       1.408e-01  6.283e-02  1.114e+03   2.240  0.02528 *  
AG3:D8       1.797e-01  6.283e-02  1.114e+03   2.860  0.00431 ** 
AG3:D9       1.495e-01  6.283e-02  1.114e+03   2.379  0.01753 *  
AG3:D10      9.783e-02  6.283e-02  1.114e+03   1.557  0.11974    
AG3:B        2.547e-02  1.125e-02  1.114e+03   2.265  0.02373 *  
D2:B         1.583e-02  1.125e-02  1.114e+03   1.407  0.15957    
D3:B         1.230e-02  1.125e-02  1.114e+03   1.094  0.27430    
D4:B         6.800e-03  1.125e-02  1.114e+03   0.605  0.54556    
D5:B        -9.405e-04  1.125e-02  1.114e+03  -0.084  0.93338    
D6:B         5.956e-03  1.125e-02  1.114e+03   0.530  0.59655    
D7:B         1.027e-02  1.125e-02  1.114e+03   0.914  0.36117    
D8:B         3.757e-03  1.125e-02  1.114e+03   0.334  0.73843    
D9:B         3.426e-03  1.125e-02  1.114e+03   0.305  0.76075    
D10:B        1.420e-03  1.125e-02  1.114e+03   0.126  0.89954    
AG3:D2:B    -1.001e-02  1.601e-02  1.114e+03  -0.625  0.53192    
AG3:D3:B    -2.698e-02  1.602e-02  1.114e+03  -1.685  0.09231 .  
AG3:D4:B    -1.426e-02  1.591e-02  1.114e+03  -0.897  0.37010    
AG3:D5:B    -1.175e-02  1.591e-02  1.114e+03  -0.739  0.46012    
AG3:D6:B    -2.698e-02  1.591e-02  1.114e+03  -1.696  0.09009 .  
AG3:D7:B    -2.096e-02  1.613e-02  1.114e+03  -1.300  0.19395    
AG3:D8:B    -3.155e-02  1.613e-02  1.114e+03  -1.957  0.05063 .  
AG3:D9:B    -2.559e-02  1.613e-02  1.114e+03  -1.587  0.11287    
AG3:D10:B   -8.855e-03  1.613e-02  1.114e+03  -0.549  0.58305    
---
Signif. codes:  0 ‘***’ 0.001 ‘**’ 0.01 ‘*’ 0.05 ‘.’ 0.1 ‘ ’ 1
```


```
Correlation matrix not shown by default, as p = 40 > 12.
Use print(x, correlation=TRUE)  or
    vcov(x)        if you need it
```


```
anova(m_acc_norm_placebo)
```


```
Type III Analysis of Variance Table with Satterthwaite's method
        Sum Sq  Mean Sq NumDF   DenDF F value    Pr(>F)    
AG     0.01545 0.015449     1   19.16  1.3956 0.2519126    
D      0.34317 0.038130     9 1114.01  3.4446 0.0003368 ***
B      0.17410 0.174104     1 1114.00 15.7279  7.78e-05 ***
AG:D   0.16081 0.017868     9 1114.01  1.6141 0.1063009    
AG:B   0.05148 0.051475     1 1114.00  4.6501 0.0312652 *  
D:B    0.12717 0.014130     9 1114.00  1.2764 0.2451861    
AG:D:B 0.07988 0.008876     9 1114.00  0.8018 0.6147092    
---
Signif. codes:  0 ‘***’ 0.001 ‘**’ 0.01 ‘*’ 0.05 ‘.’ 0.1 ‘ ’ 1
```


```
eta_squared(m_acc_norm_placebo)
```


```
# Effect Size for ANOVA (Type III)

Parameter | Eta2 (partial) |       95% CI
-----------------------------------------
AG        |           0.07 | [0.00, 1.00]
D         |           0.03 | [0.01, 1.00]
B         |           0.01 | [0.00, 1.00]
AG:D      |           0.01 | [0.00, 1.00]
AG:B      |       4.16e-03 | [0.00, 1.00]
D:B       |           0.01 | [0.00, 1.00]
AG:D:B    |       6.44e-03 | [0.00, 1.00]

- One-sided CIs: upper bound fixed at [1.00].
```

###### Note:

There was a significant difference in the slopes of the accuracy
dynamics for the placebo groups. We will compare the slopes next.


```
slope_comp <- emtrends(m_acc_norm_placebo, pairwise ~ AG, var = "B")$contrasts
```


```
NOTE: Results may be misleading due to involvement in interactions
```


```
summary(slope_comp)
```


```
 contrast  estimate      SE   df t.ratio p.value
 AG2 - AG3 -0.00778 0.00361 1114  -2.156  0.0313

Results are averaged over the levels of: D 
Degrees-of-freedom method: kenward-roger
```

###### Note:

The slope in the older-placebo group was slightly steeper than in the
middle-placebo group.

###### Accuracy dynamics in the middle-aged groups


```
data_subset <- subset(df_acc_norm, D != 0 & AG == 2)
data_subset <- droplevels(data_subset)
levels(data_subset$D)
```


```
 [1] "1"  "2"  "3"  "4"  "5"  "6"  "7"  "8"  "9"  "10"
```


```
levels(data_subset$AG)
```


```
[1] "2"
```


```
levels(data_subset$S)
```


```
[1] "1" "2"
```


```
levels(data_subset$Group)
```


```
[1] "Middle-Placebo" "Middle-Verum"
```


```
m_acc_norm_middle <- lmer(formula = Acc_n ~ S*D*B + (1 | Code), data=data_subset)
summary(m_acc_norm_middle)
```


```
Linear mixed model fit by REML. t-tests use Satterthwaite's method ['lmerModLmerTest']
Formula: Acc_n ~ S * D * B + (1 | Code)
   Data: data_subset

REML criterion at convergence: -1453.5

Scaled residuals: 
    Min      1Q  Median      3Q     Max 
-8.0236 -0.3979  0.0603  0.4929  2.9363 

Random effects:
 Groups   Name        Variance Std.Dev.
 Code     (Intercept) 0.04293  0.2072  
 Residual             0.01272  0.1128  
Number of obs: 1175, groups:  Code, 20

Fixed effects:
              Estimate Std. Error         df t value Pr(>|t|)    
(Intercept)  9.850e-01  7.345e-02  2.811e+01  13.411 9.70e-14 ***
S2           2.622e-02  1.039e-01  2.811e+01   0.252 0.802580    
D2           1.453e-02  4.826e-02  1.117e+03   0.301 0.763368    
D3           2.275e-01  4.695e-02  1.117e+03   4.845 1.45e-06 ***
D4           1.750e-01  4.695e-02  1.117e+03   3.728 0.000202 ***
D5           1.390e-01  4.695e-02  1.117e+03   2.961 0.003136 ** 
D6           1.730e-01  4.863e-02  1.117e+03   3.557 0.000391 ***
D7           2.359e-01  4.695e-02  1.117e+03   5.025 5.85e-07 ***
D8           1.598e-01  4.695e-02  1.117e+03   3.404 0.000688 ***
D9           2.364e-01  4.695e-02  1.117e+03   5.035 5.56e-07 ***
D10          2.252e-01  4.986e-02  1.117e+03   4.517 6.94e-06 ***
B            2.438e-02  8.525e-03  1.117e+03   2.860 0.004310 ** 
S2:D2       -4.964e-02  6.733e-02  1.117e+03  -0.737 0.461165    
S2:D3       -2.390e-01  6.640e-02  1.117e+03  -3.600 0.000332 ***
S2:D4       -1.655e-01  6.640e-02  1.117e+03  -2.493 0.012824 *  
S2:D5       -1.202e-01  6.640e-02  1.117e+03  -1.810 0.070537 .  
S2:D6       -1.634e-01  6.760e-02  1.117e+03  -2.417 0.015825 *  
S2:D7       -2.299e-01  6.640e-02  1.117e+03  -3.462 0.000556 ***
S2:D8       -1.479e-01  6.640e-02  1.117e+03  -2.228 0.026111 *  
S2:D9       -2.098e-01  6.640e-02  1.117e+03  -3.159 0.001623 ** 
S2:D10      -1.880e-01  6.848e-02  1.117e+03  -2.745 0.006157 ** 
S2:B        -2.700e-02  1.206e-02  1.117e+03  -2.240 0.025290 *  
D2:B         1.872e-02  1.239e-02  1.117e+03   1.511 0.131046    
D3:B        -2.566e-02  1.206e-02  1.117e+03  -2.128 0.033531 *  
D4:B        -1.456e-02  1.206e-02  1.117e+03  -1.207 0.227545    
D5:B        -7.561e-03  1.206e-02  1.117e+03  -0.627 0.530693    
D6:B        -1.695e-02  1.243e-02  1.117e+03  -1.363 0.173172    
D7:B        -2.703e-02  1.206e-02  1.117e+03  -2.242 0.025146 *  
D8:B        -1.385e-02  1.206e-02  1.117e+03  -1.149 0.250913    
D9:B        -2.824e-02  1.206e-02  1.117e+03  -2.343 0.019322 *  
D10:B       -2.189e-02  1.279e-02  1.117e+03  -1.712 0.087143 .  
S2:D2:B     -2.886e-03  1.728e-02  1.117e+03  -0.167 0.867427    
S2:D3:B      3.796e-02  1.705e-02  1.117e+03   2.226 0.026180 *  
S2:D4:B      2.136e-02  1.705e-02  1.117e+03   1.253 0.210609    
S2:D5:B      6.620e-03  1.705e-02  1.117e+03   0.388 0.697872    
S2:D6:B      2.290e-02  1.732e-02  1.117e+03   1.322 0.186292    
S2:D7:B      3.731e-02  1.705e-02  1.117e+03   2.188 0.028868 *  
S2:D8:B      1.761e-02  1.705e-02  1.117e+03   1.033 0.302000    
S2:D9:B      3.167e-02  1.705e-02  1.117e+03   1.857 0.063512 .  
S2:D10:B     2.331e-02  1.757e-02  1.117e+03   1.327 0.184911    
---
Signif. codes:  0 ‘***’ 0.001 ‘**’ 0.01 ‘*’ 0.05 ‘.’ 0.1 ‘ ’ 1
```


```
Correlation matrix not shown by default, as p = 40 > 12.
Use print(x, correlation=TRUE)  or
    vcov(x)        if you need it
```


```
anova(m_acc_norm_middle)
```


```
Type III Analysis of Variance Table with Satterthwaite's method
       Sum Sq  Mean Sq NumDF   DenDF F value    Pr(>F)    
S     0.02259 0.022586     1   18.77  1.7760 0.1985889    
D     0.50261 0.055845     9 1117.01  4.3914 1.171e-05 ***
B     0.16607 0.166070     1 1117.00 13.0588 0.0003153 ***
S:D   0.30270 0.033633     9 1117.01  2.6447 0.0049334 ** 
S:B   0.04702 0.047017     1 1117.00  3.6971 0.0547608 .  
D:B   0.20286 0.022540     9 1117.00  1.7724 0.0692508 .  
S:D:B 0.16252 0.018058     9 1117.00  1.4200 0.1743834    
---
Signif. codes:  0 ‘***’ 0.001 ‘**’ 0.01 ‘*’ 0.05 ‘.’ 0.1 ‘ ’ 1
```


```
eta_squared(m_acc_norm_middle)
```


```
# Effect Size for ANOVA (Type III)

Parameter | Eta2 (partial) |       95% CI
-----------------------------------------
S         |           0.09 | [0.00, 1.00]
D         |           0.03 | [0.01, 1.00]
B         |           0.01 | [0.00, 1.00]
S:D       |           0.02 | [0.00, 1.00]
S:B       |       3.30e-03 | [0.00, 1.00]
D:B       |           0.01 | [0.00, 1.00]
S:D:B     |           0.01 | [0.00, 1.00]

- One-sided CIs: upper bound fixed at [1.00].
```

###### Accuracy dynamics in the older groups


```
data_subset <- subset(df_acc_norm, D != 0 & AG == 3)
data_subset <- droplevels(data_subset)
levels(data_subset$D)
```


```
 [1] "1"  "2"  "3"  "4"  "5"  "6"  "7"  "8"  "9"  "10"
```


```
levels(data_subset$AG)
```


```
[1] "3"
```


```
levels(data_subset$S)
```


```
[1] "1" "2"
```


```
levels(data_subset$Group)
```


```
[1] "Older-Placebo" "Older-Verum"
```


```
m_acc_norm_older <- lmer(formula = Acc_n ~ S*D*B + (1 | Code), data=data_subset)
summary(m_acc_norm_older)
```


```
Linear mixed model fit by REML. t-tests use Satterthwaite's method ['lmerModLmerTest']
Formula: Acc_n ~ S * D * B + (1 | Code)
   Data: data_subset

REML criterion at convergence: 943.2

Scaled residuals: 
     Min       1Q   Median       3Q      Max 
-12.5720  -0.3295   0.0562   0.3611   4.5509 

Random effects:
 Groups   Name        Variance Std.Dev.
 Code     (Intercept) 0.9859   0.9929  
 Residual             0.1039   0.3223  
Number of obs: 1166, groups:  Code, 20

Fixed effects:
              Estimate Std. Error         df t value Pr(>|t|)    
(Intercept)    1.21284    0.32802   21.35985   3.697 0.001307 ** 
S2            -0.22110    0.46389   21.35985  -0.477 0.638478    
D2             0.61522    0.13419 1108.00304   4.585 5.07e-06 ***
D3             0.53447    0.13419 1108.00304   3.983 7.26e-05 ***
D4             0.59187    0.13419 1108.00304   4.411 1.13e-05 ***
D5             0.79402    0.13419 1108.00304   5.917 4.37e-09 ***
D6             0.65031    0.13795 1108.00789   4.714 2.74e-06 ***
D7             0.81238    0.13419 1108.00304   6.054 1.93e-09 ***
D8             0.89330    0.13419 1108.00304   6.657 4.40e-11 ***
D9             0.63379    0.13419 1108.00304   4.723 2.62e-06 ***
D10            0.96474    0.13419 1108.00304   7.189 1.20e-12 ***
B              0.12375    0.02437 1108.00304   5.079 4.45e-07 ***
S2:D2         -0.62086    0.19181 1108.00579  -3.237 0.001244 ** 
S2:D3         -0.42299    0.19018 1108.00439  -2.224 0.026340 *  
S2:D4         -0.51235    0.18978 1108.00304  -2.700 0.007046 ** 
S2:D5         -0.66750    0.18978 1108.00304  -3.517 0.000454 ***
S2:D6         -0.49733    0.19245 1108.00553  -2.584 0.009888 ** 
S2:D7         -0.66535    0.19249 1108.00930  -3.457 0.000568 ***
S2:D8         -0.70146    0.19249 1108.00930  -3.644 0.000281 ***
S2:D9         -0.45748    0.19249 1108.00930  -2.377 0.017637 *  
S2:D10        -0.82945    0.19249 1108.00930  -4.309 1.78e-05 ***
S2:B          -0.10090    0.03446 1108.00304  -2.928 0.003478 ** 
D2:B          -0.06601    0.03446 1108.00304  -1.916 0.055653 .  
D3:B          -0.05363    0.03446 1108.00304  -1.556 0.119892    
D4:B          -0.07263    0.03446 1108.00304  -2.108 0.035283 *  
D5:B          -0.12317    0.03446 1108.00304  -3.575 0.000366 ***
D6:B          -0.08494    0.03540 1108.00304  -2.399 0.016595 *  
D7:B          -0.11703    0.03446 1108.00304  -3.396 0.000707 ***
D8:B          -0.10693    0.03446 1108.00304  -3.103 0.001963 ** 
D9:B          -0.04429    0.03446 1108.00304  -1.285 0.198984    
D10:B         -0.12059    0.03446 1108.00304  -3.500 0.000485 ***
S2:D2:B        0.07186    0.04906 1108.00475   1.465 0.143307    
S2:D3:B        0.03900    0.04907 1108.00788   0.795 0.426940    
S2:D4:B        0.06516    0.04873 1108.00305   1.337 0.181426    
S2:D5:B        0.11048    0.04873 1108.00305   2.267 0.023574 *  
S2:D6:B        0.06391    0.04940 1108.00304   1.294 0.196065    
S2:D7:B        0.10634    0.04940 1108.00304   2.153 0.031567 *  
S2:D8:B        0.07913    0.04940 1108.00304   1.602 0.109501    
S2:D9:B        0.02213    0.04940 1108.00304   0.448 0.654350    
S2:D10:B       0.11315    0.04940 1108.00304   2.290 0.022187 *  
---
Signif. codes:  0 ‘***’ 0.001 ‘**’ 0.01 ‘*’ 0.05 ‘.’ 0.1 ‘ ’ 1
```


```
Correlation matrix not shown by default, as p = 40 > 12.
Use print(x, correlation=TRUE)  or
    vcov(x)        if you need it
```


```
anova(m_acc_norm_older)
```


```
Type III Analysis of Variance Table with Satterthwaite's method
      Sum Sq Mean Sq NumDF   DenDF F value    Pr(>F)    
S     0.3003 0.30035     1   18.28  2.8909  0.106034    
D     5.1664 0.57404     9 1108.01  5.5253 1.812e-07 ***
B     2.6441 2.64409     1 1108.00 25.4499 5.302e-07 ***
S:D   2.6239 0.29154     9 1108.01  2.8062  0.002916 ** 
S:B   0.9667 0.96673     1 1108.00  9.3050  0.002340 ** 
D:B   1.4880 0.16534     9 1108.00  1.5914  0.112853    
S:D:B 1.1121 0.12356     9 1108.00  1.1893  0.297839    
---
Signif. codes:  0 ‘***’ 0.001 ‘**’ 0.01 ‘*’ 0.05 ‘.’ 0.1 ‘ ’ 1
```


```
eta_squared(m_acc_norm_older)
```


```
# Effect Size for ANOVA (Type III)

Parameter | Eta2 (partial) |       95% CI
-----------------------------------------
S         |           0.14 | [0.00, 1.00]
D         |           0.04 | [0.02, 1.00]
B         |           0.02 | [0.01, 1.00]
S:D       |           0.02 | [0.00, 1.00]
S:B       |       8.33e-03 | [0.00, 1.00]
D:B       |           0.01 | [0.00, 1.00]
S:D:B     |       9.57e-03 | [0.00, 1.00]

- One-sided CIs: upper bound fixed at [1.00].
```


```
slope_comp <- emtrends(m_acc_norm_older, pairwise ~ S, var = "B")$contrasts
```


```
NOTE: Results may be misleading due to involvement in interactions
```


```
summary(slope_comp)
```


```
 contrast estimate     SE   df t.ratio p.value
 S1 - S2    0.0338 0.0111 1108   3.050  0.0023

Results are averaged over the levels of: D 
Degrees-of-freedom method: kenward-roger
```


```
# Calculate effect size (Estimate / SE) * sqrt(N)
eff_size_manual <- 0.0338 / 0.0111 * sqrt(20)
eff_size_manual
```


```
[1] 13.61786
```

###### Note:

The slope of the dynamics in the older-verum group was steeper than
the one in the older-placebo group.

##### Stats on electrophysiological recordings (TMS)

The following tests concern the recordings using transcranial
magnetic stimulation (TMS). These recordings were done at rest and
during movement preparation (i.e., during a reaction time task). Please
refer to the Methods in the main manuscript for details.

###### SICIrest (measurements performed while participants are at rest)


```
sicirest_file_name <-  "Python_AP3_SICIrest_stack_14_03_2022.txt"
file_path <- file.path(paste(python_data_directory, sicirest_file_name, sep='/'))
df_sici_all = read.delim(file_path, header = TRUE, na.strings = "NN")
df_sici <- subset(df_sici_all, TP == "V1pe" | TP == "V2po")
df_sici <- droplevels(df_sici)
head(df_sici)
```


```
df_sici$ID <- as.factor(df_sici$ID)
df_sici$AG <- as.factor(df_sici$AG)
df_sici$S <- as.factor(df_sici$S)
df_sici$TP <- as.factor(df_sici$TP)
df_sici$SICIratio <- as.numeric(df_sici$SICIratio)
levels(df_sici$ID)
```


```
 [1] "217201" "217202" "217203" "217205" "217206" "217207" "217209" "217210" "217252" "217253" "217254" "217255" "217257" "217258" "217259" "217260" "217301" "217302"
[19] "217303" "217304" "217305" "217307" "217308" "217310" "217311" "217312" "217351" "217352" "217353" "217354" "217355" "217356" "217357" "217359" "217360"
```


```
levels(df_sici$AG)
```


```
[1] "2" "3"
```


```
levels(df_sici$S)
```


```
[1] "1" "2"
```


```
levels(df_sici$TP)
```


```
[1] "V1pe" "V2po"
```

###### Select a model for SICIrest


```
m1 <- lm(formula = SICIratio ~ AG*S*TP, data=df_sici)
m2 <- lmer(formula = SICIratio ~ AG*S*TP + (1|ID), data=df_sici)
BIC(m1, m2)
```

###### Model choice

The model including random intercepts for individuals is worse than
the plain linear model according to the BIC, so we will use m1 to model
the SICI recordings.


```
m_sici <- lm(formula = SICIratio ~ AG*S*TP, data=df_sici)
summary(m_sici)
```


```
Call:
lm(formula = SICIratio ~ AG * S * TP, data = df_sici)

Residuals:
     Min       1Q   Median       3Q      Max 
-0.45923 -0.24459 -0.08527  0.19191  0.85431 

Coefficients:
              Estimate Std. Error t value Pr(>|t|)   
(Intercept)    0.35262    0.11572   3.047  0.00343 **
AG3            0.09222    0.15526   0.594  0.55478   
S2             0.02377    0.15905   0.149  0.88171   
TPV2po        -0.05012    0.16940  -0.296  0.76837   
AG3:S2        -0.04501    0.22227  -0.202  0.84022   
AG3:TPV2po     0.12554    0.22389   0.561  0.57706   
S2:TPV2po      0.22884    0.23237   0.985  0.32866   
AG3:S2:TPV2po -0.34555    0.31970  -1.081  0.28408   
---
Signif. codes:  0 ‘***’ 0.001 ‘**’ 0.01 ‘*’ 0.05 ‘.’ 0.1 ‘ ’ 1

Residual standard error: 0.3273 on 60 degrees of freedom
  (2 observations deleted due to missingness)
Multiple R-squared:  0.06036,   Adjusted R-squared:  -0.04927 
F-statistic: 0.5506 on 7 and 60 DF,  p-value: 0.7925
```


```
anova(m_sici)
```


```
Analysis of Variance Table

Response: SICIratio
          Df Sum Sq  Mean Sq F value Pr(>F)
AG         1 0.0394 0.039375  0.3675 0.5466
S          1 0.0068 0.006798  0.0634 0.8020
TP         1 0.0363 0.036348  0.3393 0.5624
AG:S       1 0.1865 0.186494  1.7407 0.1921
AG:TP      1 0.0097 0.009720  0.0907 0.7643
S:TP       1 0.0090 0.009015  0.0841 0.7728
AG:S:TP    1 0.1252 0.125168  1.1683 0.2841
Residuals 60 6.4282 0.107137
```


```
eta_squared(m_sici)
```


```
# Effect Size for ANOVA (Type I)

Parameter | Eta2 (partial) |       95% CI
-----------------------------------------
AG        |       6.09e-03 | [0.00, 1.00]
S         |       1.06e-03 | [0.00, 1.00]
TP        |       5.62e-03 | [0.00, 1.00]
AG:S      |           0.03 | [0.00, 1.00]
AG:TP     |       1.51e-03 | [0.00, 1.00]
S:TP      |       1.40e-03 | [0.00, 1.00]
AG:S:TP   |           0.02 | [0.00, 1.00]

- One-sided CIs: upper bound fixed at [1.00].
```

###### Note:

There are no significant effects. However, by design and based on
past studies, we would expect age-related differences in intracortical
inhibition (please see e.g., Heise et al., 2013), so we will test the
age groups separately to look for effects of stimulation.


```
data_subset <- subset(df_sici, AG == 2)
data_subset <- droplevels(data_subset)
levels(data_subset$TP)
```


```
[1] "V1pe" "V2po"
```


```
levels(data_subset$AG)
```


```
[1] "2"
```


```
levels(data_subset$S)
```


```
[1] "1" "2"
```


```
m_sici_middle <- lm(formula = SICIratio ~ S*TP, data=data_subset)
summary(m_sici_middle)
```


```
Call:
lm(formula = SICIratio ~ S * TP, data = data_subset)

Residuals:
     Min       1Q   Median       3Q      Max 
-0.42398 -0.19768 -0.07667  0.12998  0.85431 

Coefficients:
            Estimate Std. Error t value Pr(>|t|)   
(Intercept)  0.35262    0.11355   3.105  0.00432 **
S2           0.02377    0.15606   0.152  0.88004   
TPV2po      -0.05012    0.16622  -0.302  0.76525   
S2:TPV2po    0.22884    0.22800   1.004  0.32411   
---
Signif. codes:  0 ‘***’ 0.001 ‘**’ 0.01 ‘*’ 0.05 ‘.’ 0.1 ‘ ’ 1

Residual standard error: 0.3212 on 28 degrees of freedom
  (2 observations deleted due to missingness)
Multiple R-squared:  0.08895,   Adjusted R-squared:  -0.008665 
F-statistic: 0.9112 on 3 and 28 DF,  p-value: 0.4481
```


```
anova(m_sici_middle)
```


```
Analysis of Variance Table

Response: SICIratio
          Df  Sum Sq  Mean Sq F value Pr(>F)
S          1 0.13730 0.137302  1.3311 0.2584
TP         1 0.04075 0.040753  0.3951 0.5347
S:TP       1 0.10391 0.103915  1.0074 0.3241
Residuals 28 2.88811 0.103147
```


```
eta_squared(m_sici_middle)
```


```
# Effect Size for ANOVA (Type I)

Parameter | Eta2 (partial) |       95% CI
-----------------------------------------
S         |           0.05 | [0.00, 1.00]
TP        |           0.01 | [0.00, 1.00]
S:TP      |           0.03 | [0.00, 1.00]

- One-sided CIs: upper bound fixed at [1.00].
```


```
data_subset <- subset(df_sici, AG == 3)
data_subset <- droplevels(data_subset)
levels(data_subset$TP)
```


```
[1] "V1pe" "V2po"
```


```
levels(data_subset$AG)
```


```
[1] "3"
```


```
levels(data_subset$S)
```


```
[1] "1" "2"
```


```
m_sici_older <- lm(formula = SICIratio ~ S*TP, data=data_subset)
summary(m_sici_older)
```


```
Call:
lm(formula = SICIratio ~ S * TP, data = data_subset)

Residuals:
    Min      1Q  Median      3Q     Max 
-0.4592 -0.2500 -0.1396  0.2632  0.6596 

Coefficients:
            Estimate Std. Error t value Pr(>|t|)    
(Intercept)  0.44484    0.10518   4.229 0.000183 ***
S2          -0.02124    0.15777  -0.135 0.893759    
TPV2po       0.07543    0.14875   0.507 0.615576    
S2:TPV2po   -0.11671    0.22312  -0.523 0.604532    
---
Signif. codes:  0 ‘***’ 0.001 ‘**’ 0.01 ‘*’ 0.05 ‘.’ 0.1 ‘ ’ 1

Residual standard error: 0.3326 on 32 degrees of freedom
Multiple R-squared:  0.02521,   Adjusted R-squared:  -0.06617 
F-statistic: 0.2759 on 3 and 32 DF,  p-value: 0.8423
```


```
anova(m_sici_older)
```


```
Analysis of Variance Table

Response: SICIratio
          Df Sum Sq  Mean Sq F value Pr(>F)
S          1 0.0563 0.056310  0.5090 0.4807
TP         1 0.0050 0.004994  0.0451 0.8331
S:TP       1 0.0303 0.030268  0.2736 0.6045
Residuals 32 3.5401 0.110629
```


```
eta_squared(m_sici_older)
```


```
# Effect Size for ANOVA (Type I)

Parameter | Eta2 (partial) |       95% CI
-----------------------------------------
S         |           0.02 | [0.00, 1.00]
TP        |       1.41e-03 | [0.00, 1.00]
S:TP      |       8.48e-03 | [0.00, 1.00]

- One-sided CIs: upper bound fixed at [1.00].
```

###### SICImove (measurements performed during movement preparation, as participants perform a reaction time task)


```
sicimove_file_name <-  "Python_AP3_SICImoveModulation_31_01_2022.txt"
file_path <- file.path(paste(python_data_directory, sicimove_file_name, sep='/'))
df_move_all = read.delim(file_path, header = TRUE, na.strings = "NN")
df_move <- subset(df_move_all, TP == 1 | TP == 2)
df_move <- droplevels(df_move)
head(df_move)
```


```
df_move$ID <- as.factor(df_move$ID)
df_move$AG <- as.factor(df_move$AG)
df_move$S <- as.factor(df_move$S)
df_move$SG <- as.factor(df_move$SG)
df_move$TP <- as.factor(df_move$TP)
df_move$Mod90.20 <- as.numeric(df_move$Mod90.20)
levels(df_move$ID)
```


```
 [1] "217201" "217202" "217203" "217205" "217206" "217207" "217209" "217210" "217252" "217253" "217254" "217255" "217257" "217258" "217259" "217260" "217301" "217302"
[19] "217303" "217304" "217305" "217307" "217308" "217310" "217311" "217312" "217351" "217352" "217353" "217354" "217355" "217356" "217357" "217359" "217360"
```


```
levels(df_move$AG)
```


```
[1] "2" "3"
```


```
levels(df_move$S)
```


```
[1] "1" "2"
```


```
levels(df_move$TP)
```


```
[1] "1" "2"
```

###### Select a model for SICImove


```
m1 <- lm(formula = Mod90.20 ~ AG*S*TP, data=df_move)
m2 <- lmer(formula = Mod90.20 ~ AG*S*TP + (1|ID), data=df_move)
BIC(m1, m2)
```

###### Model choice

The model including random intercepts for individuals is much worse
than the plain linear model according to the BIC, so we will use m1 to
model the SICImove recordings.


```
m_move <- lm(formula = Mod90.20 ~ AG*S*TP, data=df_move)
summary(m_move)
```


```
Call:
lm(formula = Mod90.20 ~ AG * S * TP, data = df_move)

Residuals:
     Min       1Q   Median       3Q      Max 
-0.63229 -0.08433 -0.00232  0.09950  0.40628 

Coefficients:
             Estimate Std. Error t value Pr(>|t|)  
(Intercept)  0.110229   0.065012   1.696   0.0952 .
AG3         -0.118780   0.087223  -1.362   0.1784  
S2          -0.008839   0.089350  -0.099   0.9215  
TP2          0.251583   0.095168   2.644   0.0105 *
AG3:S2       0.091118   0.124865   0.730   0.4684  
AG3:TP2     -0.228419   0.125775  -1.816   0.0744 .
S2:TP2      -0.306544   0.132831  -2.308   0.0245 *
AG3:S2:TP2   0.398270   0.181273   2.197   0.0320 *
---
Signif. codes:  0 ‘***’ 0.001 ‘**’ 0.01 ‘*’ 0.05 ‘.’ 0.1 ‘ ’ 1

Residual standard error: 0.1839 on 59 degrees of freedom
  (3 observations deleted due to missingness)
Multiple R-squared:  0.2758,    Adjusted R-squared:  0.1899 
F-statistic:  3.21 on 7 and 59 DF,  p-value: 0.00599
```


```
anova(m_move)
```


```
Analysis of Variance Table

Response: Mod90.20
          Df  Sum Sq  Mean Sq F value   Pr(>F)   
AG         1 0.13516 0.135164  3.9975 0.050181 . 
S          1 0.00004 0.000036  0.0011 0.974057   
TP         1 0.10638 0.106376  3.1461 0.081271 . 
AG:S       1 0.31551 0.315512  9.3313 0.003379 **
AG:TP      1 0.00380 0.003802  0.1124 0.738568   
S:TP       1 0.03556 0.035558  1.0516 0.309322   
AG:S:TP    1 0.16322 0.163218  4.8272 0.031954 * 
Residuals 59 1.99493 0.033812                    
---
Signif. codes:  0 ‘***’ 0.001 ‘**’ 0.01 ‘*’ 0.05 ‘.’ 0.1 ‘ ’ 1
```


```
eta_squared(m_move)
```


```
# Effect Size for ANOVA (Type I)

Parameter | Eta2 (partial) |       95% CI
-----------------------------------------
AG        |           0.06 | [0.00, 1.00]
S         |       1.81e-05 | [0.00, 1.00]
TP        |           0.05 | [0.00, 1.00]
AG:S      |           0.14 | [0.03, 1.00]
AG:TP     |       1.90e-03 | [0.00, 1.00]
S:TP      |           0.02 | [0.00, 1.00]
AG:S:TP   |           0.08 | [0.00, 1.00]

- One-sided CIs: upper bound fixed at [1.00].
```

###### SICImove in middle-aged adults


```
data_subset <- subset(df_move, AG == 2)
data_subset <- droplevels(data_subset)
levels(data_subset$TP)
```


```
[1] "1" "2"
```


```
levels(data_subset$AG)
```


```
[1] "2"
```


```
levels(data_subset$S)
```


```
[1] "1" "2"
```


```
m_move_middle <- lm(formula = Mod90.20 ~ S*TP, data=data_subset)
summary(m_move_middle)
```


```
Call:
lm(formula = Mod90.20 ~ S * TP, data = data_subset)

Residuals:
     Min       1Q   Median       3Q      Max 
-0.30071 -0.07079 -0.02182  0.07650  0.40628 

Coefficients:
             Estimate Std. Error t value Pr(>|t|)   
(Intercept)  0.110229   0.057096   1.931  0.06410 . 
S2          -0.008839   0.078470  -0.113  0.91115   
TP2          0.251583   0.083580   3.010  0.00561 **
S2:TP2      -0.306544   0.116657  -2.628  0.01400 * 
---
Signif. codes:  0 ‘***’ 0.001 ‘**’ 0.01 ‘*’ 0.05 ‘.’ 0.1 ‘ ’ 1

Residual standard error: 0.1615 on 27 degrees of freedom
  (3 observations deleted due to missingness)
Multiple R-squared:  0.3753,    Adjusted R-squared:  0.3059 
F-statistic: 5.407 on 3 and 27 DF,  p-value: 0.004799
```


```
anova(m_move_middle)
```


```
Analysis of Variance Table

Response: Mod90.20
          Df  Sum Sq  Mean Sq F value Pr(>F)  
S          1 0.17487 0.174867  6.7052 0.0153 *
TP         1 0.06811 0.068114  2.6118 0.1177  
S:TP       1 0.18008 0.180078  6.9050 0.0140 *
Residuals 27 0.70414 0.026079                 
---
Signif. codes:  0 ‘***’ 0.001 ‘**’ 0.01 ‘*’ 0.05 ‘.’ 0.1 ‘ ’ 1
```


```
eta_squared(m_move_middle)
```


```
# Effect Size for ANOVA (Type I)

Parameter | Eta2 (partial) |       95% CI
-----------------------------------------
S         |           0.20 | [0.02, 1.00]
TP        |           0.09 | [0.00, 1.00]
S:TP      |           0.20 | [0.03, 1.00]

- One-sided CIs: upper bound fixed at [1.00].
```


```
my_model <- m_move_middle
level_1 <- "S"

my_model.compare <- emmeans(my_model, level_1, by="TP")
my_model.compare.pairs <- pairs(my_model.compare, adjust='tukey')
test(my_model.compare.pairs, side='two-sided')
```


```
TP = 1:
 contrast estimate     SE df t.ratio p.value
 S1 - S2   0.00884 0.0785 27   0.113  0.9111

TP = 2:
 contrast estimate     SE df t.ratio p.value
 S1 - S2   0.31538 0.0863 27   3.654  0.0011
```


```
confint(my_model.compare, calc = c(n = ~.wgt.))
```


```
TP = 1:
 S emmean     SE df n lower.CL upper.CL
 1 0.1102 0.0571 27 8 -0.00692    0.227
 2 0.1014 0.0538 27 9 -0.00906    0.212

TP = 2:
 S emmean     SE df n lower.CL upper.CL
 1 0.3618 0.0610 27 7  0.23657    0.487
 2 0.0464 0.0610 27 7 -0.07881    0.172

Confidence level used: 0.95
```


```
eff_size(my_model.compare, sigma = sigma(my_model), edf = 23)
```


```
TP = 1:
 contrast effect.size    SE df lower.CL upper.CL
 S1 - S2       0.0547 0.486 27   -0.942     1.05

TP = 2:
 contrast effect.size    SE df lower.CL upper.CL
 S1 - S2       1.9529 0.607 27    0.707     3.20

sigma used for effect sizes: 0.1615 
Confidence level used: 0.95
```


```
my_model <- m_move_middle
level_1 <- "TP"

my_model.compare <- emmeans(my_model, level_1, by="S")
my_model.compare.pairs <- pairs(my_model.compare, adjust='tukey')
test(my_model.compare.pairs, side='two-sided')
```


```
S = 1:
 contrast  estimate     SE df t.ratio p.value
 TP1 - TP2   -0.252 0.0836 27  -3.010  0.0056

S = 2:
 contrast  estimate     SE df t.ratio p.value
 TP1 - TP2    0.055 0.0814 27   0.675  0.5052
```


```
confint(my_model.compare, calc = c(n = ~.wgt.))
```


```
S = 1:
 TP emmean     SE df n lower.CL upper.CL
 1  0.1102 0.0571 27 8 -0.00692    0.227
 2  0.3618 0.0610 27 7  0.23657    0.487

S = 2:
 TP emmean     SE df n lower.CL upper.CL
 1  0.1014 0.0538 27 9 -0.00906    0.212
 2  0.0464 0.0610 27 7 -0.07881    0.172

Confidence level used: 0.95
```


```
eff_size(my_model.compare, sigma = sigma(my_model), edf = 23)
```


```
S = 1:
 contrast  effect.size    SE df lower.CL upper.CL
 TP1 - TP2       -1.56 0.566 27   -2.720   -0.396

S = 2:
 contrast  effect.size    SE df lower.CL upper.CL
 TP1 - TP2        0.34 0.506 27   -0.699    1.379

sigma used for effect sizes: 0.1615 
Confidence level used: 0.95
```

###### SICImove in older adults


```
data_subset <- subset(df_move, AG == 3)
data_subset <- droplevels(data_subset)
levels(data_subset$TP)
```


```
[1] "1" "2"
```


```
levels(data_subset$AG)
```


```
[1] "3"
```


```
levels(data_subset$S)
```


```
[1] "1" "2"
```


```
m_move_older <- lm(formula = Mod90.20 ~ S*TP, data=data_subset)
summary(m_move_older)
```


```
Call:
lm(formula = Mod90.20 ~ S * TP, data = data_subset)

Residuals:
     Min       1Q   Median       3Q      Max 
-0.63229 -0.09165  0.01858  0.09951  0.33823 

Coefficients:
            Estimate Std. Error t value Pr(>|t|)
(Intercept) -0.00855    0.06351  -0.135    0.894
S2           0.08228    0.09527   0.864    0.394
TP2          0.02316    0.08982   0.258    0.798
S2:TP2       0.09173    0.13473   0.681    0.501

Residual standard error: 0.2008 on 32 degrees of freedom
Multiple R-squared:  0.135, Adjusted R-squared:  0.0539 
F-statistic: 1.665 on 3 and 32 DF,  p-value: 0.1942
```


```
anova(m_move_older)
```


```
Analysis of Variance Table

Response: Mod90.20
          Df  Sum Sq  Mean Sq F value  Pr(>F)  
S          1 0.14596 0.145960  3.6185 0.06617 .
TP         1 0.03679 0.036785  0.9119 0.34676  
S:TP       1 0.01870 0.018697  0.4635 0.50088  
Residuals 32 1.29079 0.040337                  
---
Signif. codes:  0 ‘***’ 0.001 ‘**’ 0.01 ‘*’ 0.05 ‘.’ 0.1 ‘ ’ 1
```


```
eta_squared(m_move_older)
```


```
# Effect Size for ANOVA (Type I)

Parameter | Eta2 (partial) |       95% CI
-----------------------------------------
S         |           0.10 | [0.00, 1.00]
TP        |           0.03 | [0.00, 1.00]
S:TP      |           0.01 | [0.00, 1.00]

- One-sided CIs: upper bound fixed at [1.00].
```
